# Molecular dynamics of outer membrane-embedded polysaccharide secretion porins reveals closed resting-state surface gates targetable by virtual fragment screening for drug hotspot identification

**DOI:** 10.1101/2023.12.21.572922

**Authors:** Tanos C. C. França, Fares Saïdi, Alain Ajamian, Salim T. Islam, Steven R. LaPlante

**Author notes:** Corresponding authors Steven R. LaPlante Salim T. Islam.

## Abstract

Recent advances in iterative neural-network analyses (e.g. AlphaFold2 and RoseTTA fold) have been revolutionary for protein 3D-structure prediction, especially for difficult-to-manipulate α-helical/β-barrel integral membrane proteins. These model structures are calculated based on the co-evolution of amino acids within the protein of interest and similarities to existing protein structures; local effects of the membrane on folding and stability of the calculated model structures are not considered. We recently reported the discovery, 3D modelling, and characterization of 18-β-stranded outer-membrane (OM) WzpX, WzpS, and WzpB β-barrel secretion porins for the exopolysaccharide (EPS), major spore coat polysaccharide (MASC), and biosurfactant polysaccharide (BPS) pathways (respectively) in the Gram-negative social predatory bacterium *Myxococcus xanthus* DZ2. However, information was not obtained regarding the dynamic behavior of surface-gating WzpX/S/B loop domains, nor on potential treatments to inactivate these porins. Herein, we developed a molecular dynamics (MD) protocol to study the core stability and loop dynamism of neural network-based integral membrane protein structure models embedded in an asymmetric OM bilayer, using the *M. xanthus* WzpX, WzpS, and WzpB proteins as test candidates. This was accomplished through integration of the CHARMM-graphical user interface (GUI) and Molecular Operating Environment (MOE) workflows to allow for rapid simulation system setup and facilitate data analysis. In addition to serving as a method of model-structure validation, our molecular dynamics simulations revealed minimal movement of extracellular WzpX/S/B loops in the absence of an external stimulus, as well as druggable cavities between the loops. Virtual screening of a commercial fragment library against these cavities revealed putative fragment-binding hotspots on the cell-surface face of each β-barrel, along with key interacting residues, and identified promising hits for the design of potential binders capable of plugging the β-barrels and inhibiting polysaccharide secretion.

## Introduction

Iterative neural network-based advances in protein folding such as AlphaFold2^1^ and RoseTTA fold^2^ have revolutionized the field of protein structure prediction. Nowhere has this been more apparent than with the 3D modelling of proteins that traverse membrane bilayers via α-helix or β-barrel architecture; the former are typically found in the inner (cytoplasmic) membrane of bacteria, eukaryotes, mitochondria, and chloroplasts^3^, while the latter are largely found in the outer membrane (OM) of Gram-negative bacteria, mitochondria, and chloroplasts^4^. Such integral membrane proteins are often recalcitrant to overexpression and purification, let alone *in vitro* manipulation during various experiments. As such, neural network-based structure prediction has been a boon for the study of proteins localized to these subcellular compartments.

Protein model structures from applications such as AlphaFold2 and RoseTTA fold are largely based on the analysis of co-evolving amino acids within the polypeptide of interest as well as similarities to existing protein structures^1, 2^. However, for integral membrane proteins, the local effect of the membrane environment on the relative positioning and stability of the membrane-spanning α-helical or β-stranded tracts, as well as the extra-membranous loops, is not accounted for prior to establishing the final model structure. Instead, molecular dynamics (MD) simulations of integral membrane proteins inserted in a membrane bilayer environment have proven to be a powerful complementary tool for examining potential structural rearrangements/modifications due to non-solvated hydrophobic surroundings^5^.

We recently reported the discovery, 3D modelling (via AlphaFold2), and characterization of the WzpX, WzpS, and WzpB outer-membrane (OM) β-barrel secretion porins^6^ from the respective pathways for exopolysaccharide (EPS), major spore coat polysaccharide (MASC), and biosurfactant polysaccharide (BPS)^7^ in Gram-negative^8^ *Myxococcus xanthus* DZ2, a social and predatory soil bacterium^9^. This deltaproteobacterium is used as a model organism in which to study single-cell^10, 11^ and group motility^7^, developmental progression (via formation of spore-filled fruiting bodies)^7^, drug tolerance^12^, and biofilm formation^13^, with the abovementioned secreted polysaccharides having outsized impacts on all of these physiological outcomes.

The WzpX/WzpS/WzpB OM porins^6, 14^ were found to be part of the EPS/MASC/BPS Wzx/Wzy-dependent pathways^7, 15, 16^ in which undecaprenyl pyrophosphate-linked sugar repeats are flipped across the inner membrane to its periplasmic leaflet (by Wzx)^17^, where they are then polymerized^18^ (by Wzy) and secreted across the cell envelope to the cell exterior. However, WzpX/WzpS/WzpB were identified as expanded structural homologues of the porin PgaA required for secretion of poly-*N*-acetyl-D-glucosamine (PNAG) in *Escherichia coli*. Intriguingly, PNAG is made by a synthase-dependent pathway in which sugar addition in the cytoplasm is coupled to export of the growing chain across the OM by an equivalent amount. Prior to our investigation, synthase-dependent pathway-like secretion proteins had never before been identified in Wzx/Wzy-dependent pathways^19^.

WzpX/WzpS/WzpB were also found to be genetically and functionally coupled with their respective so-called outer-membrane polysaccharide export (OPX) proteins WzaX/WzaS/WzaB (required for EPS/MASC/BPS secretion); these OPX proteins are designated as “Class 3” as they do not have membrane-spanning domains and resemble the majority of OPX proteins found across all Gram-negative and Gram-positive bacteria^6^. Given the lack of OM-spanning capacity for WzpX/WzpS/WzpB, our characterization of the integral OM β-barrels WzpX/WzpS/WzpB finally provided an explanation for how the physiologically-important EPS, MASC, and BPS polymers could be secreted across the OM in *M. xanthus*^6^. However, since the AlphaFold2-generated WzpX/WzpS/WzpB model structures (**Fig. 1**) were of a specific conformation, no information was available regarding the stability of the core model structures or the dynamic behaviour of surface-gating loop domains.

**Figure 1.**
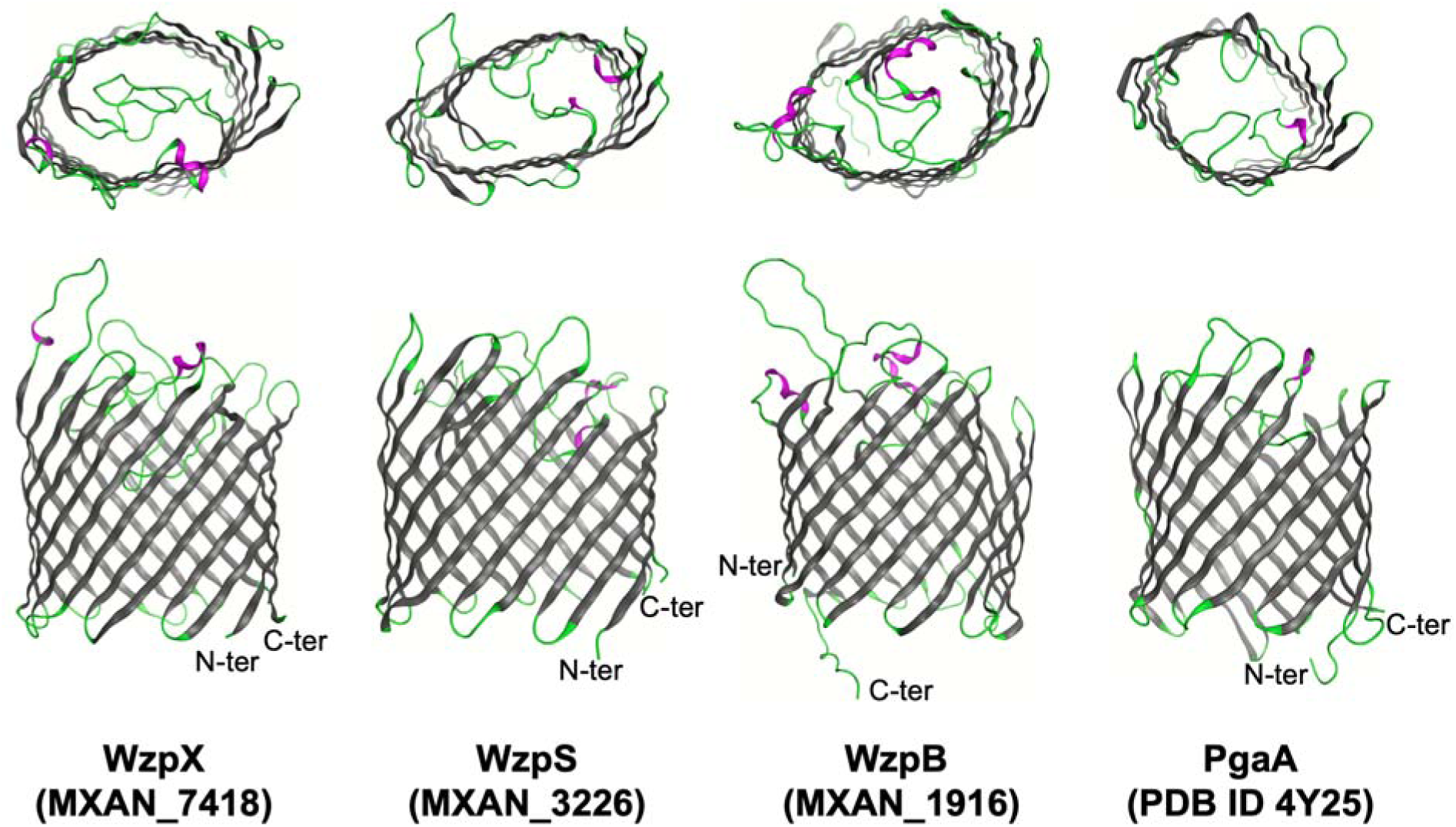
3D model structures of EPS-pathway WzpX, MASC-pathway WzpS and BPS-pathway WzpB porins^6^, as well as the X-ray crystal structure of the PNAG secretion porin PgaA (PDB 4Y25)^23^. The cell-surface (*top panels*) and side (*bottom panels*) views of the proteins are presented. Legend: β-sheets, *dark grey*; α-helices, *magenta*; unstructured loops, *green*.

A popular approach to obtaining information on macromolecular protein dynamics is use of the Molecular Operating Environment (MOE) software platform^20^; this provides a streamlined and unified environment for running MD simulations in addition to offering several other molecular modeling functionalities including (i) docking, (ii) virtual screening (VS), (iii) quantitative structure–activity relationship (QSAR), (iv) cheminformatic, and (v) computer-aided drug design (CADD) analyses. Through its versatile and user-friendly graphical user interface (GUI), molecular complexes can be constructed or imported to the MOE environment and automatically prepared for MD simulations using a set of applications for correcting molecular topologies and protonation states, generating and soaking complexes with defined periodic boundary conditions, as well as assigning partial charges and applying atomic restraints based on specified forcefield parameters. With prepared complexes in hand, MD simulation steps can be efficiently expressed using an easy MD protocol language to define temperature, pressure and energy ramps. The MOE MD framework automatically generates topology, parameter and script files for running MD simulations on parallel cluster or GPU with full preservation of force field definitions from MOE, with no need of using complex command lines. Depending on the size of the system under study this can save days of work. MD trajectories can then be imported and analyzed within the MOE environment using a set of molecular dynamics analysis methods which extract and summarize any information from the resulting trajectory files in spreadsheets from which the data can be easily plotted, making such analyses much easier to obtain by a larger segment of the scientific community. However, MOE currently lacks a tool to generate membrane-embedded protein models. Fortunately this limitation can be overcome through prior use of the membrane-builder tool of the CHARMM (Chemistry at HARvard Macromolecular Mechanics) platform^21^ and its associated GUI^22^ which enables the construction of a variety of membrane-embedded models as well as the generation of input files for further MD simulations.

In this investigation, we integrated the CHARMM-GUI and MOE workflows, and applied it towards the examination of WzpX/WzpS/WzpB model stability and dynamism, relative to that of the experimentally-derived *E. coli* PgaA X-ray crystal structure, in a simulated asymmetric OM bilayer. These analyses reveal the minimal flexibility of extracellular loop domains in the absence of an external stimulus as well as the existence of apical cavities. Virtual screening (VS) studies against these extracellular cavities using a commercial fragment library enabled determination of the cavities’ fingerprint profiles, revealing the existence of promising druggable hotspots and ranked potential hits suitable for inhibition of polysaccharide secretion via future drug design of targeted binders.

## Methodology

### Construction of the MD model systems

The MD model systems — consisting of PgaA or the related β-barrels WzpX (MXAN_7418), WzpS (MXAN_3226), and WzpB (MXAN_1916) embedded in an asymmetric *E. coli* OM bilayer — were constructed using the *Bilayer Builder* option of the *Membrane Builder* input generator tool of the CHARMM-GUI server (https://www.charmm-gui.org/)^22^. First the .pdb files of each model were uploaded to the server and the orientation option set as *“Run PPM 2.0”*, while water in the protein pore was generated with the *“Using the protein geometry”* option. Next, the *“Heterogeneous Lipid”* option was chosen since the *E. coli* OM is asymmetric. The box type chosen was *“Rectangular”* and the *“Length of Z”* was based on *“Water thickness”* with a minimum water height of 22.5 Å on the top and bottom of the system. The *“Length of XY”* was based on the *“Ratios of lipid components”* and the *“Length of X and Y”* was set to 80 Å. The composition of the lower (periplasmic) leaflet of the OM was set to previously established values^24^: 89% of 1-palmitoyl(16:0)-2-palmitoleoyl(16:1 cis-9)-phosphatidylethanolamine (PPPE), 1% of 1,10 -palmitoyl-2,20 -vacenoyl cardiolipin with a net charge of _2e (PVCL2), and 11% of 1-palmi-toyl(16:0)-2-vacenoyl(18:1 cis-11)-phosphatidylglycerol (PVPG), while the upper (surface) leaflet was 100% composed of lipopolysaccharide (LPS). The LPS chemical structure used was that from *E. coli* without *“O-units”* or *“chemical modification”*. As the behavior of the PgaA structure was to serve as the reference control, PgaA, as well as WzpX, WzpS, and WzpB, were all inserted in the same simulated OM. Though the LPS structure from *M. xanthus* is not identical to that of *E. coli*^8, 25^, integral OM proteins from *M. xanthus* have been repeatedly shown to properly display and interact when heterologously expressed in the *E. coli* OM^11, 26^, indicating a compatible environment.

The *“System Building Option”* chosen was the *“Replacement method”* with the *“Check lipid ring penetration”* option selected. The *“Component Building Options”* were as follows: *“Include Ions”*, the *“Ion replacing method”* was *“Distance”* with KCl at 0.15 M as the only *“Basic Ion Type”*. The .pdb file obtained was then uploaded into the MOE molecular modeling package (https://www.chemcomp.com/index.htm) for further preparation to run MD simulations.

### MD simulations

As mentioned above, the MD simulations were performed through the MOE molecular modeling package. To begin, each system was opened in the MOE main window and visually inspected in order to fix missing bonds and incorrect atom types and names. The systems were then optimized using the *“Protein - Structure Preparation”* tool to achieve the proper bond lengths, angles and charges compatible with the physiological environment and neutralized through the replacement of water molecules by Na^+^ or Cl^−^ ions. Once optimized, the files were prepared for the MD simulations using the MOE *“Compute/Simulations/Dynamics”* module. The files were then transferred via Linux terminal to an account at Calcul Quebec (https://www.calculquebec.ca/) where the MD simulations were run via command line according to Calcul Quebec protocols in GPU:v100:1 graphic cards with 500 MB of allocated memory. Following completion of the simulations, the output files were downloaded from Calcul Quebec to a local computer to be analyzed using MOE tools.

As previously described^27^, simulations were performed in triplicate using the AMBER10:EHT^28^ force field and NAMD software^29^ with a cut-off of 10 kcal/mol for electrostatic interactions and a range between 8 and 10 kcal/mol for van der Waals interactions. The protocol before each production step included 5000 steps of energy minimization followed by 100 ps of isothermal-isobaric ensemble (NPT) simulation and 200 ns of isothermal-isochoric ensemble (NVT). Considering that the models were quite complex owing to several different elements (protein, phospholipids, saccharides, ions and solvent), packed close to each other, it was necessary to run three short 10-ns MD simulations before the principal production step, with gradual release of the system. This was done to avoid crashing the system through many clashes caused by the sudden release of all elements at the same time. The first MD simulation was performed with position restriction (PR) of the whole system except for the membrane. In the second MD simulation, only the protein was kept restrained. For the third MD simulation, only the protein backbones were restrained. Finally, for the fourth MD simulation, a production step of 1 µs was carried out with all components unrestrained.

The trajectories obtained were analyzed using the “*md_analysis*” tool and the database viewer (DBV) menu of MOE. The Root Mean Square Deviation (RMSD, a measure of how much one atom deviates from its initial position), and Root Mean Square Fluctuation (RMSF, a measure of how much one amino acid deviates from its initial position), were calculated for the β-barrels in comparison to the first frames of each production step. Finally, plots of the MD results were created using GraphPad Prism and figures were created with MOE and PyMol ^30^.

### Virtual fragment screening

Representative frames from the MD simulations corresponding to the averaged value of RMSD after stabilization were further submitted to VS of a fragment library designed and kindly provided by NMX Research and Solutions Inc. (https://www.nmxresearch.com/) and marketed as the BIONET - 2^nd^ Generation Premium Fragment Library Pack from Key Organics (https://www.keyorganics.net/downloads-bionet-databases/), using the software Molegro Virtual Docker (MVD)^31^. This library contains 1166 fragments previously curated for NMR binding experiments; it was designed to maximize conformational diversity and the number of fragments found in approved drugs, as well as excluding potential aggregating and pan assay interference (PAIN)^32^ compounds. Fragment libraries are ideal for finding hotspots in molecular targets since fragments can serve as chemical biology probes to expose pockets and pocket features of potential interest.^33^ This facilitates the identification of hits as well as potential leads for initiating drug discovery projects. It was downloaded in the .sdf format and exported into a MOE database.mdb file where the fragments were “washed” to the appropriate bond lengths, angles, charges, and protonation states compatible with the physiological environment, and then saved in .mol2 format. To run the VS, representative MD simulation frames of WzpX, WzpS, WzpB and PgaA were opened in the MVD^31^ main window together with the washed library. Each spherical search space was set to cover the whole external surface of the model including all loops, with the central coordinates and corresponding radius summarized (**Table S1**). Thirty runs per model were performed with a maximum of 3000 interactions and the percentage of returning poses was set to 20%. All poses obtained with “*moldockscore negative*” were selected for further analysis.

### Fingerprint analysis

The protein ligand interaction fingerprint (PLIF) application of MOE was used to map the most relevant interactions in the hotspots of each β-barrel. For this we used the representative frames from the MD simulations mentioned above and databanks in the .mdb format containing only the poses observed in the hotspots. The fingerprints were prepared and generated using the PLIF set-up panel with the weak and strong energy thresholds for the 9 types of protein– ligand interaction which normally comprise fingerprints, set as indicated (**Table 1**). The resulting fingerprints results were compiled in bar-code plots.

**Table 1.** energy thresholds used to generate the fingerprints of the hotspots.

| Type of interaction |  | Weak<br>(kcal.mol<sup>-1</sup>) | Strong<br>(kcal.mol<sup>-1</sup>) |
| --- | --- | --- | --- |
| Sidechain | H-donor | 0.5 | 1.5 |
|  | H-acceptor | 0.5 | 1.5 |
| Backbone | H-donor | 0.5 | 1.5 |
|  | H-acceptor | 0.5 | 1.5 |
| Solvent | H-donor | 0.5 | 1.5 |
|  | H-acceptor | 0.5 | 1.5 |
| Ionic attraction |  | 0.5 | 3.5 |
| Metal ligation |  | 0.5 | 3.5 |
| Arene attraction |  | 0.5 | 1.0 |

## Results and Discussion

### MD simulation

Following integration of the CHARMM-GUI and MOE workflows, MD simulations of WzpX, WzpS, WzpB, and PgaA inserted in an OM bilayer yielded RMSD plots that never exceeded 4.5 Å, showing very similar profiles for the experimental structure (PgaA) and the three models (WzpX/S/B) (**Fig. 2**). These results suggest an appropriate dynamic behavior of all systems and consistency of the 3D structures of the models when compared to PgaA, serving as an additional step of structure validation for the three β-barrel models (**Fig. 1**). This also suggests a robust integration between each protein and the surrounding membrane components, corroborating the consistency of the protein–membrane models constructed herein.

**Figure 2.**
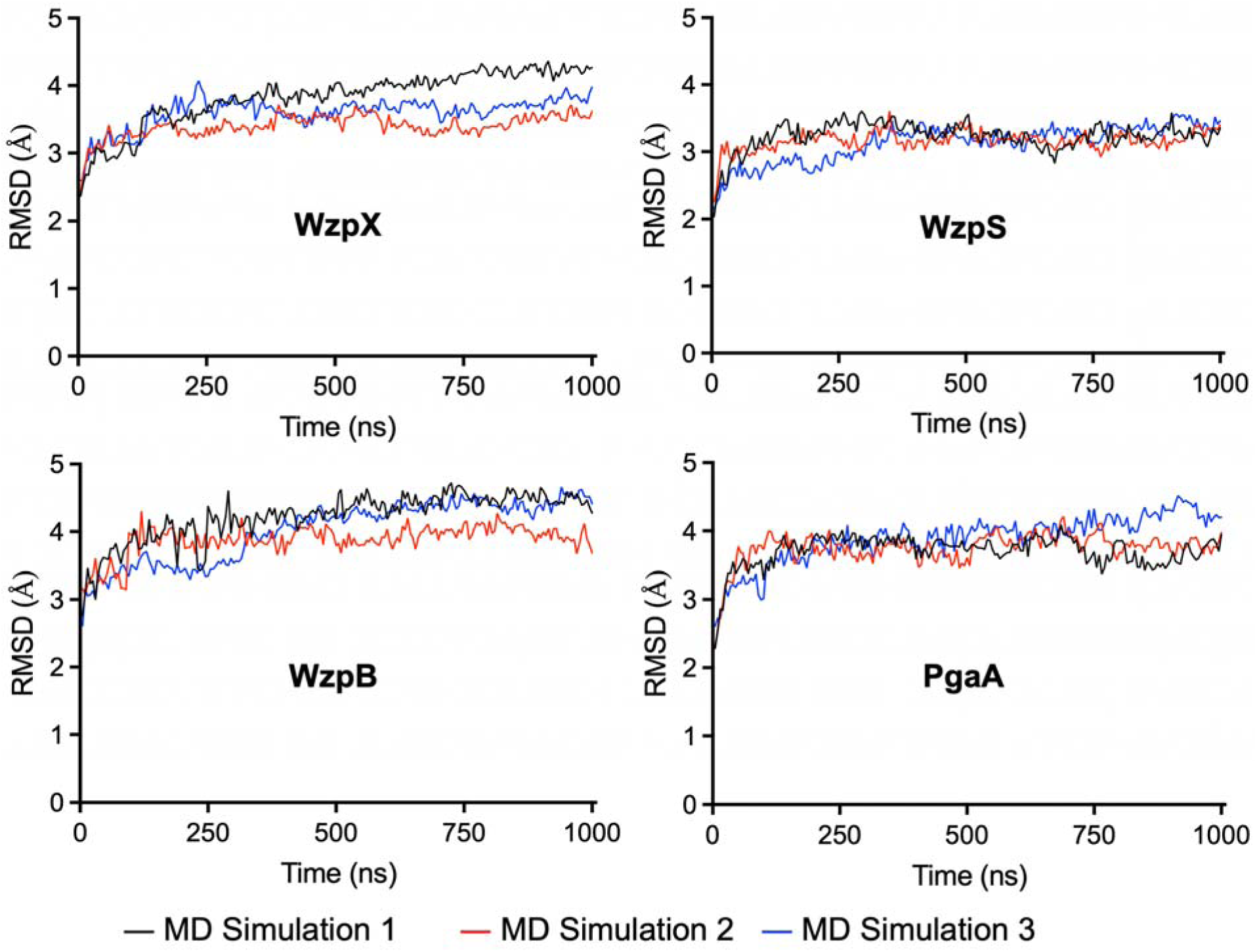
Plots of variation of RMSD of WzpX, WzpS, WzpB, and PgaA embedded in an asymmetric OM bilayer after 1 μs of MD simulation in triplicate.

Fluctuations in the position of each amino acid during the MD simulations were then probed via analysis of the average RMSF plots (**Fig. 3**). As expected, the highest fluctuations corresponded with loop regions; however, the fluctuations did not surpass 5.0 Å, suggesting an overall low mobility of the loops. The only exceptions observed were for two external loops of the WzpX model structure (amino acids 101-137, and amino acids 252-268,) which showed fluctuations up to 5.03 and 7.05 Å (respectively). The low fluctuations below 2.0 Å observed for the trans-membrane β strands reflect their stability during the MD simulations and integration with the membrane. These results are illustrated by the overlapping of frames collected during the MD simulations (**Fig. 4**), where for all systems, the loops did not move enough on their own to open the extracellular face of the β-barrels. This is consistent with the hypothesis that the opening of WzpX/S/B to the extracellular milieu is not a spontaneous motion and instead depends on the binding of some triggering factor to the β-barrel lumen (e.g. the translocating polymer) and/or periplasmic face (e.g. WzaX/S/B Class-3 OPX protein)^6, 14^ capable of causing a major conformational change in the extracellular WzpX/S/B loops.

**Figure 3.**
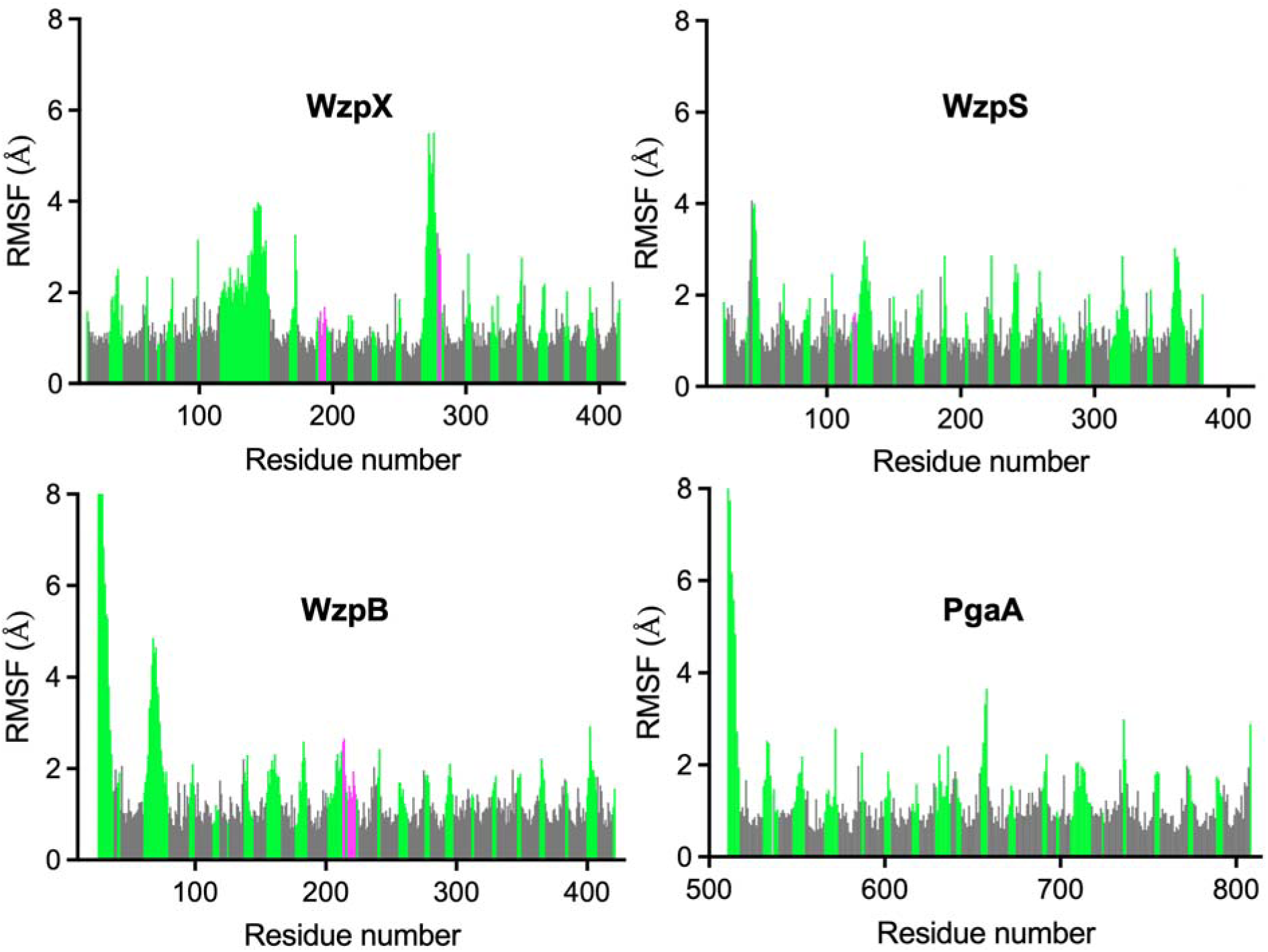
Plots of the average RMSF for WzpX, WzpS, WzpB, and PgaA after 1 μs of MD simulation in triplicate. Colors correspond to secondary-structure elements from Figure 1, i.e. β-sheets (*dark grey*), α-helices (*magenta*), and unstructured loops (*green*).

**Figure 4.**
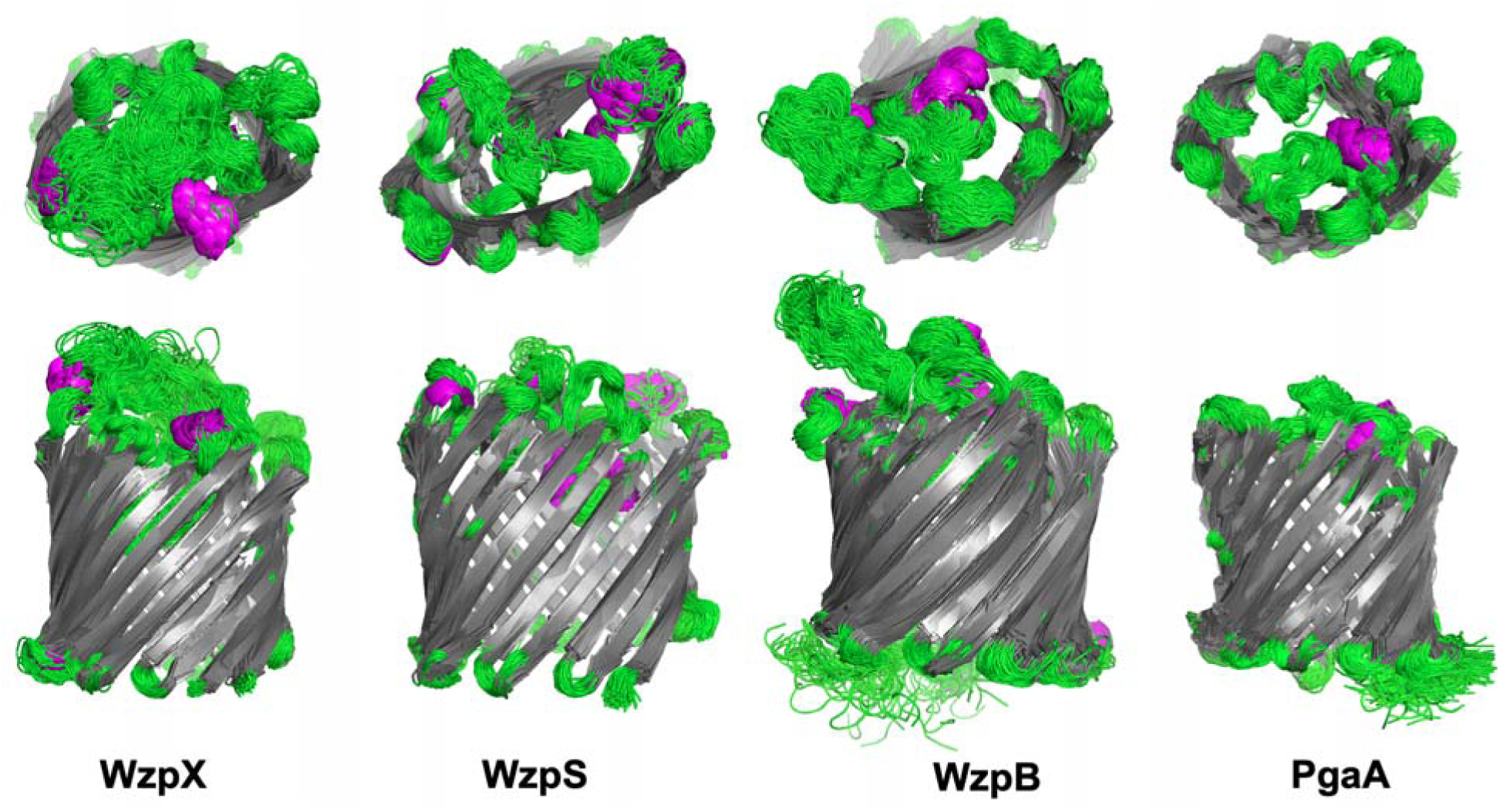
Superposition of 200 frames collected at 5-ns intervals during the MD simulations. Both the extracellular (*top row*) and side-facing (*bottom row*) views have been provided. Colors correspond to secondary-structure elements from Figures 1 and 3, *i.e.* β-sheets (*dark grey*), α-helices (*magenta*), and unstructured loops (*green*).

Analysis of the extracellular views of representative frames of the MD simulations depicted as electrostatic surfaces (**Fig. 5**) revealed the presence of several pockets among the loops which could be targeted in the future for the purpose of drug design. Moreover, each protein displayed a unique surface, showing distinct cavities and a different pattern of pockets. This suggests that it is possible to design and/or identify binding compounds capable of selectively blocking one specific β-barrel by exploring the different shapes and potential interactions with the pockets in question.

**Figure 5.**
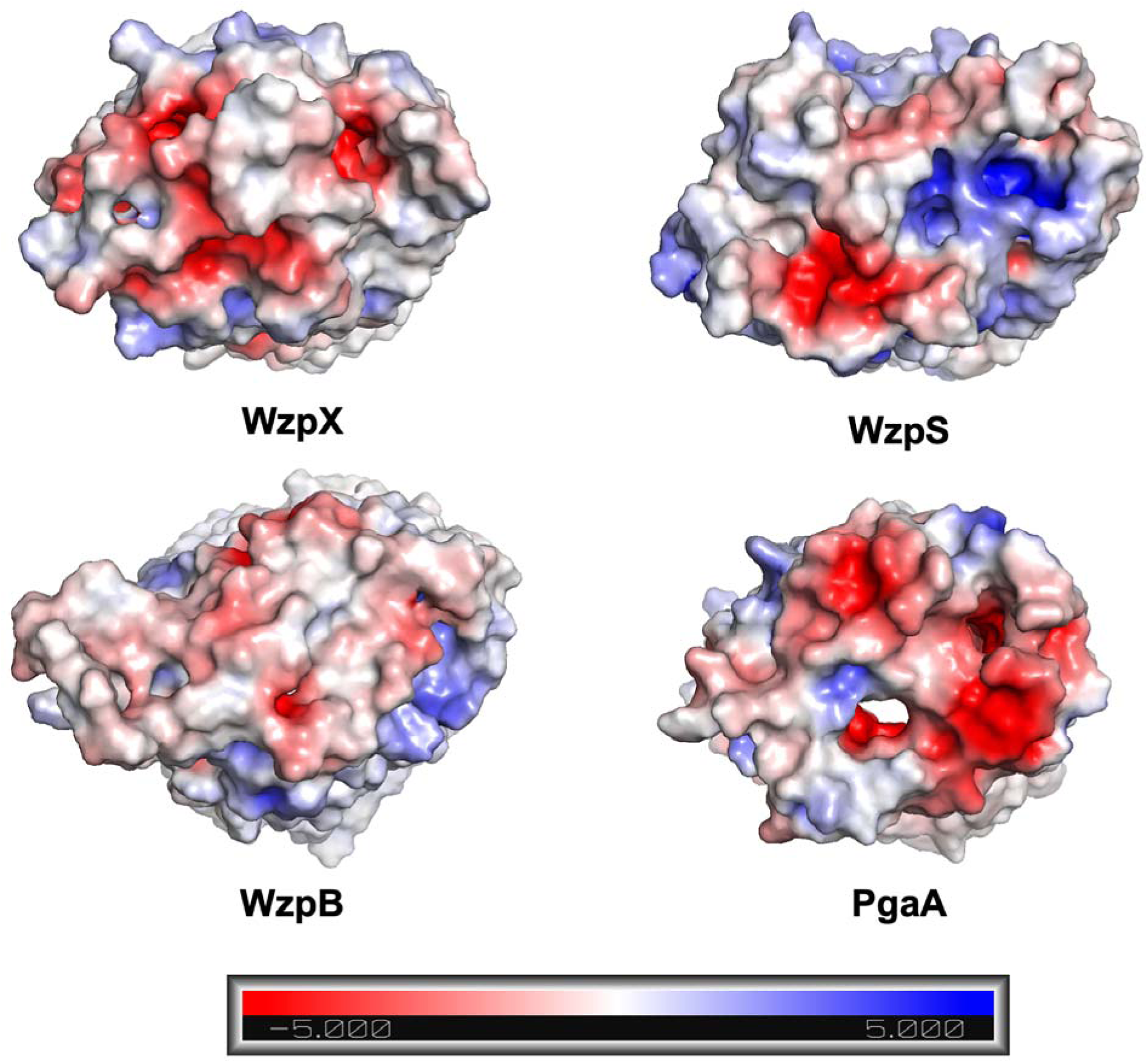
Extracellular view of electrostatic surfaces of representative frames collected during the MD simulations (± 5.000 kT/e). The regions with the most negative electrostatic potential are shown in *red* while the regions with the most positive electrostatic potential are shown in *blue*.

### Fragment Library Virtual Screening

Having uncovered druggable pockets on the extracellular faces of WzpX, WzpS, WzpB, and PgaA, we applied this information towards the identification of compound fragments via VS with the potential to bind these various pockets and provide a basis for downstream drug development. The VS returned 1082 fragments for WzpX, 1001 for WzpS, 1143 for WzpB, and 599 for PgaA (**Tables S2-S5**). The MolDockScores obtained were also significantly different among the β-barrels as shown in the plot of MolDockScore *versus* fragment number (**Fig. 6**). WzpX and WzpS were the proteins showing the best-ranked fragments with lowest energy values of −221.77 and −203.53 Kcal/mol, respectively. Conversely, for WzpB and PgaA, no fragment with energy below −110 Kcal/mol was observed and only 38 and 11 fragments (respectively) scored below −100 Kcal/mol. These results suggest a higher affinity of the fragments for WzpX and WzpS and that by extension, these two β-barrels might be more druggable compared to WzpB and PgaA.

**Figure 6.**
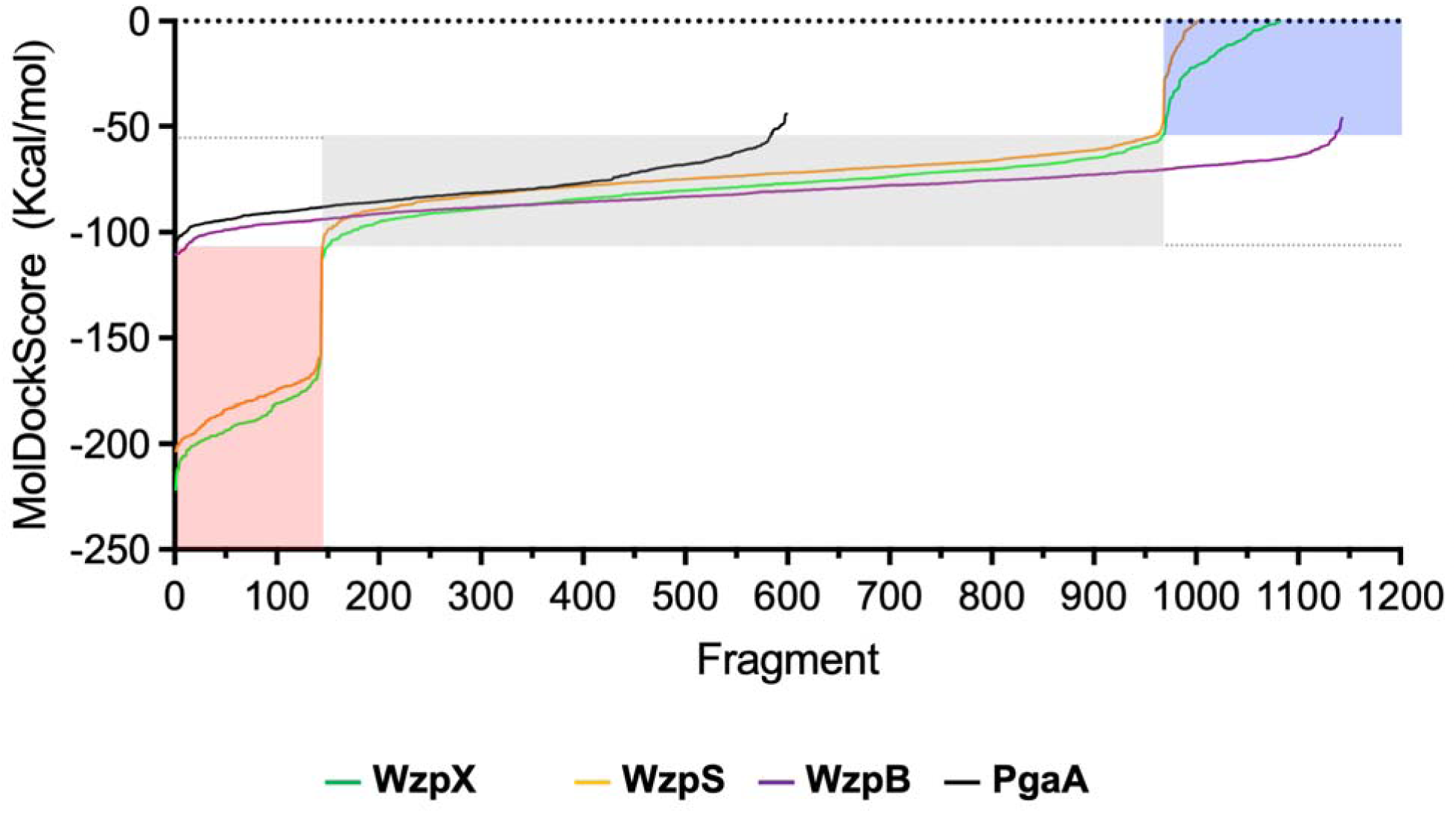
Plots of MolDockSocre (Kcal/mol) vs. fragment number obtained from VS. The colored areas correspond to the energy ranges of the negatively-charged (*red*), neutral (*gray*) and positively-charged (*blue*) fragments observed for the proteins.

The VS results also evidenced a unique charge-related energy distribution for the specific cavities of WzpX and WzpS; in both cases, negatively-charged fragments ranked best and positively-charged fragments ranked worst, while neutral fragments were distributed in between (**Fig. 6**). This was not observed for WzpB and PgaA, which showed charged and neutral fragments distributed over the whole energy curve with a slight prevalence of neutral and positively charged residues at the lowest energy regions. Interestingly, the range of energy where most of the fragment poses fell for these two proteins corresponded to the neutral range (**Fig. 6**) which is the same energy range observed for the neutral fragments over WzpX and WzpS.

The distribution of fragment poses over the β-barrels’ surfaces enabled detection of hotspots of preferred binding among the external loops. For WzpX, the most populated binding zone was located in the central cavity between the main loops (**Fig. 7**). This cavity concentrated 40.76% of the fragments and also all fragments ranked below −200 Kcal/mol. Similarly, WzpS also concentrated most of the fragments (39.56%) in a central hotspot. However, this porin also exhibited a second highly-populated hotspot which, together with the first one, concentrated the 112 best fragments, all ranking below −172 Kcal/mol. For WzpB, most fragments (75.94%) concentrated at the three hotspots around the main loop at the extracellular face of the porin (**Fig. 7**). These hotspots also hosted 33 of the 38 best-ranked fragments. Similar to WzpS, PgaA concentrated most fragments (85.47%) in the extracellular loops (see **Fig. 7**), including the 11 fragments ranked below −100 Kcal/mol.

**Figure 7.**
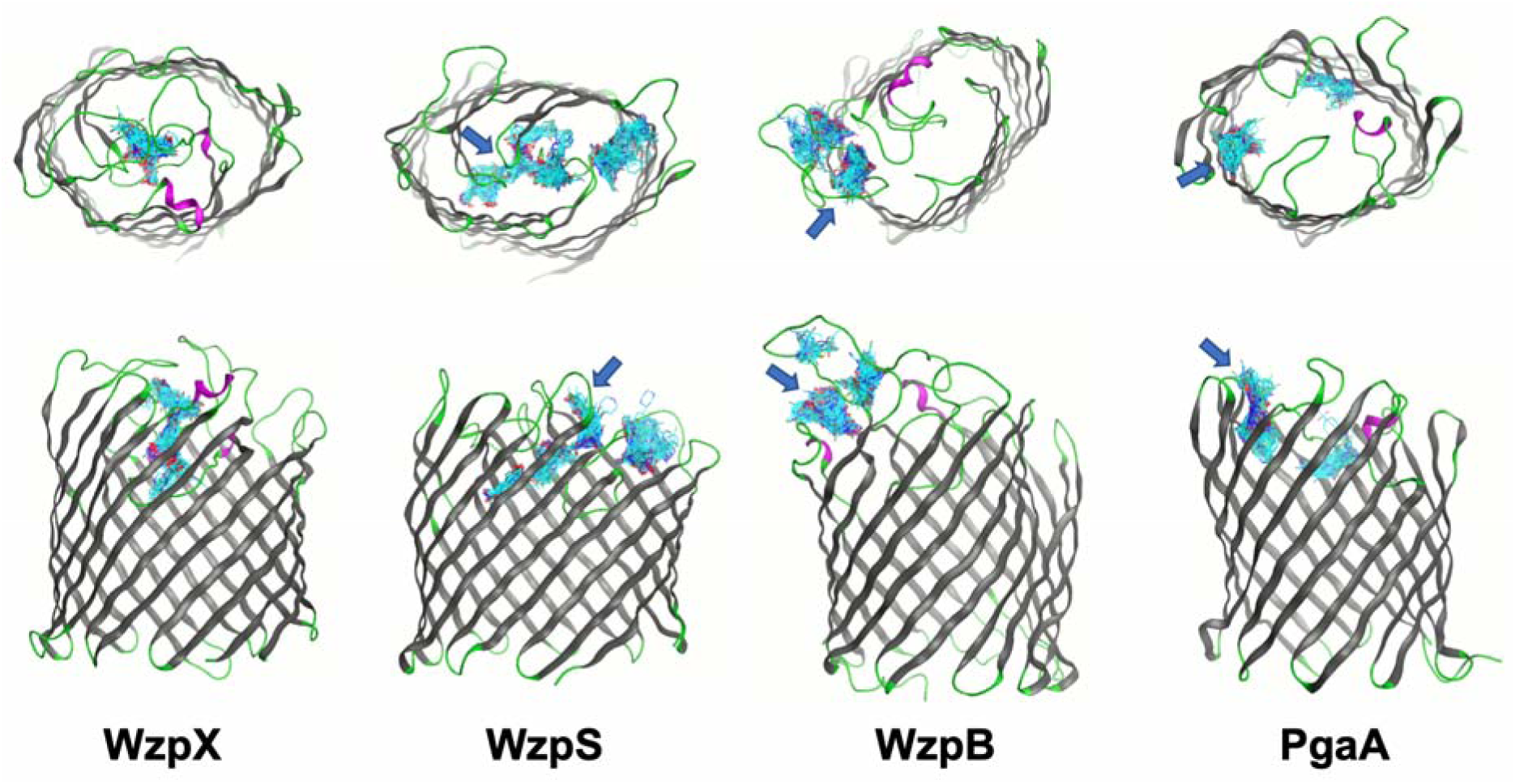
Concentration of fragment poses (*shown in cyan*) showcasing the most populated hotspots of the β-barrels. The arrows point to the hotspots concentrating the most poses.

To map hotspot residues of interest for future drug design, we analyzed the MolDockScores and corresponding amino acids involved in potential H-bonds for the 10 best-ranked fragment poses from VS (**Table S6, Figs. S1-S40**) for each β-barrel. As mentioned above, all of the 10 best-ranked fragments from VS against WzpX and WzpS were negatively charged while those for WzpB and PgaA were either neutral or positively charged (except for 3K-528S against PgaA). These data point to differences in compound fragment specificity among the β-barrels that can be further explored in the design of selective inhibitors. Importantly, only (i) one fragment was found to be a common putative binder between WzpX/WzpS and WzpB/PgaA, (ii) three fragments were common between WzpX and WzpS, (iii) only one fragment was equivalent between WzpB and PgaA, and (iv) no common fragments were detected between WzpX/WzpS and WzpB (**Table S6**). These findings point to differences in the hotspots to be explored for the downstream design of selective inhibitors.

For WzpX, up to 27 different amino acids were involved in some form of interaction with the screened fragments (**Figs. S1 - S10 and Table 2**). In particular, Arg305 stands out since this residue was observed forming H-bonds with all 10 of the best-ranked fragments. Residues Thr104, Ser108, Leu109, Asp112, Gly115, Thr116, Asp118, and Asp304, were identified as other potential sources of H-bonds while residues Ser108, Thr116, Asp118, showed potential arene-H interactions.

**Table 2.** Hotspot residues of each β-barrel.

| β-barrel | Hotspot residues |
| --- | --- |
| WzpX | Leu101, Thr104, Leu106, Gly107, Ser108, Leu109, Asp112, Thr113, Pro114, Gly115, Thr116, Pro117, Asp118, Gly119, Pro120, Leu121, Arg122, Phe141, Ser143, Ser170, Ala180, Val181, Glu257, Thr303, Asp304, Arg305, Leu306 |
| WzpS<br>(hotspot 1) | Arg103, Ala104, Gly105, Leu106, Phe115, Arg137, Phe138, Glu139, Ala140, Val141, Gly151, Tyr152, Ala153, His154, Thr155, Gln177, Phe179, Val250, Gly251, Ala252, Asn253 |
| WzpS<br>(hotspot 2) | Arg15, Asp17, Asp19, Arg21, Met31, Arg60, Thr96, Asp97, Pro98, Arg108, Ser109, Thr110, Arg347 |
| WzpB | Gly33, Val34, Tyr36, Phe37, Ser38, Pro39, Thr40, Asn95, Ser96, Ser97, Ala98, Ala99, Leu177, Ser178, Ser179, Val180 |
| PgaA | Glu704, Tyr714, Asn715, Pro716, Ile717, Lys718, Thr719, Asp721, Ser751, Trp752, Gln753, Lys754, His755, Tyr756, Val761, Arg787, Pro788, Tyr789, Asp790 |

Five of the top-10 fragments identified for WzpS docked in the most populated hotspot (herein termed “hotspot 1”) while four docked in the second most populated zone (“hotspot 2”) and one docked in the interface between the two hotspots. Twenty-one residues were observed in VS interactions with the fragments in hotspot 1, while 13 residues were identified in hotspot 2 (**Figs. S11 - S20, Table 2**). Arg103 and Asn253 formed potential H-bonds and arene-cation or arene-H interactions in hotspot 1 while for hotspot 2 the key H-bonding residues were Arg60 and Thr110, with the latter also showing potential for arene-H interactions. Fragment docking between the two hotspots (3K-528S) showed potential H-bonds and arene-H interactions with Gly105 (from hotspot 1) and Arg108 (from hotspot 2).

The 10-best fragments docked in WzpB concentrated in the most populated hotspot and showed potential interactions with 16 residues. Potential H-bonds were observed for residues Gly33, Val34, Tyr36, Thr40, Asn95, Ser96, Leu177, Ser179, while arene-H interactions were observed for residues Pro39, Ala98, and Ser179. Arg40 and Ser179 were involved in H-bonding for the majority of poses (**Figs. S21 - S30 and Table 2**).

Nineteen residues were observed to form potential interactions with the fragments in the main hotspot of PgaA; Pro716, Lys718, Gln753, Lys754, Arg787, Pro788, Tyr789, and Asp790 were implicated in H-bonding, while Lys754, Tyr789 were observed in potential arene-H interactions. Lys754, Arg787, Pro788, and Tyr789 were those residues most frequently implicated in fragment binding (**Figs. S31 - S40, Table 2**).

### Fingerprint analysis of the hotspots

The bar code plots shown in **Fig. 8** reinforces the relevance of the arginines in the hotspots of WzpX (Arg118 and Arg305) and WzpS (Arg60 and Arg108). These residues performed mostly ionic attraction (purple bars), side chain hydrogen bond acceptor (red bars), arene attraction (orange bars) and backbone hydrogen bond acceptor (salmon bars) with the fragments. This corroborates the higher affinities for negatively charged residues observed in the VS studies for these two proteins. For WzpB and PgaA on the other hand ionic attractions (purple bars) didn’t show much relevance and the most prevalent interactions were side chain hydrogen bond acceptor (red bars) and backbone hydrogen bond acceptor (cyan bars). This is aligned with the trend observed in the VS studies suggesting that these two proteins would be less selective to charged compounds.

**Figure 8.**
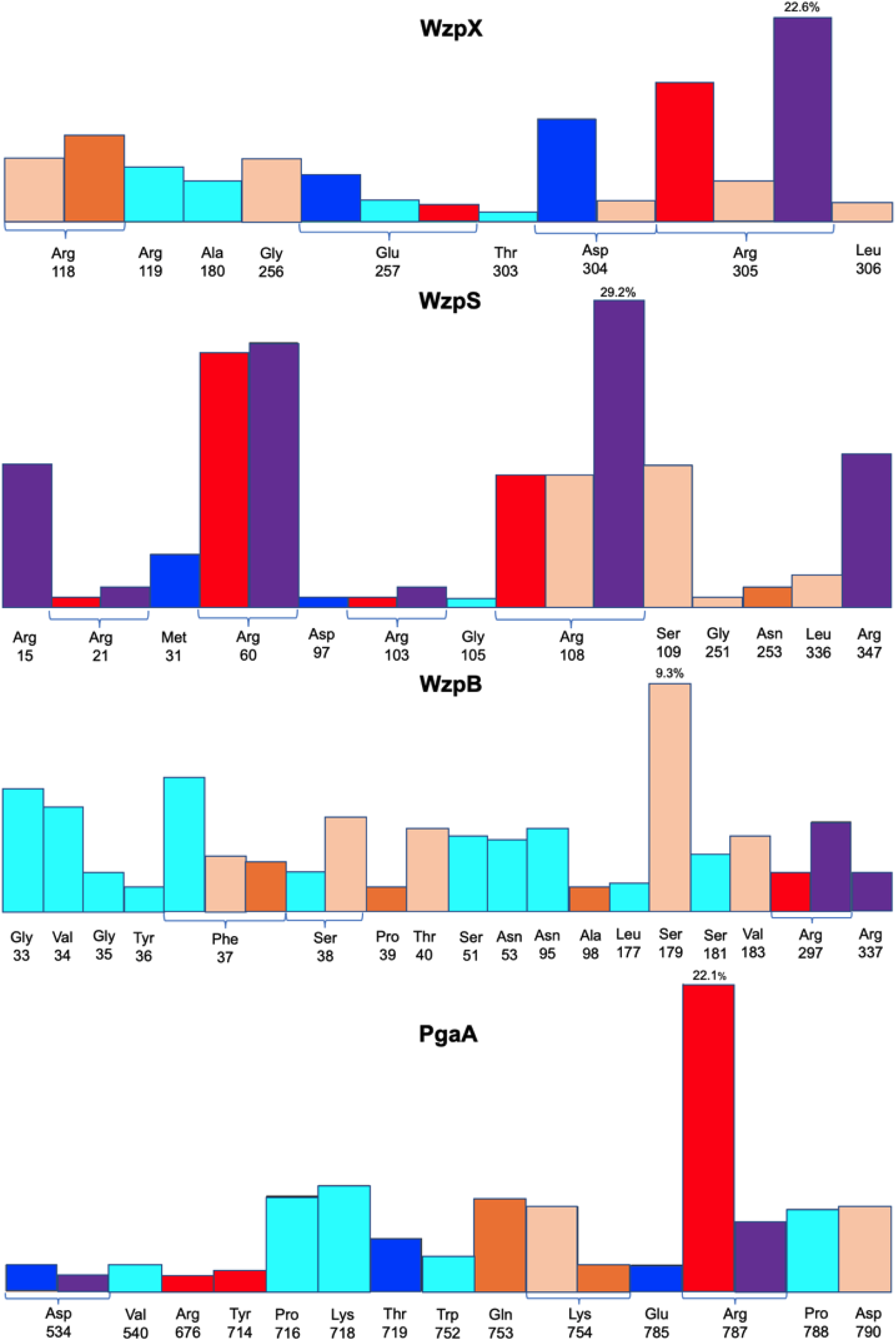
Fingerprints of the strong interactions on the hot spots of WzpX, WzpS, WzpB and PgaA. Color bars code: Red = Side chain hydrogen bond acceptor; Blue = Side chain hydrogen bond donor; Purple = Ionic attraction; Orange = Arene attraction; Salmon = Backbone hydrogen bond acceptor; Cyan = Backbone hydrogen bond donor.

## Conclusions

We have herein reported a new and straightforward protocol to run MD simulations of membrane-embedded systems merging resources from the CHARMM-GUI^22^ server and the MOE molecular modeling package^20^, in order to take advantage of the more intuitive and user-friendly environment provided by the latter package to run MD simulations. This workflow will complement other tools already reported in the literature^34^.

Results of the MD simulations performed using our protocol on the experimental structure of PgaA as well as the neural network-based model structures of WzpX, WzpS, and WzpB, corroborated the quality and consistency of the models and also revealed the low mobility of the extracellular loops of these four β-barrels. These findings support the hypothesis that an internal triggering mechanism is needed for the opening of these β-barrels; this would be consistent with nascent PNAG/EPS/MASC/BPS polymers (arriving from the periplasm) being translocated through the interior of OM-spanning PgaA/WzpX/WzpS/WzpB^6^, with this translocation step contributing to the opening of the extracellular loops of the respective porins. For WzpX/WzpS/WzpB, this opening may happen in conjunction with proposed interactions^6, 14^ between the β-barrels and the periplasmic Class-3 OPX proteins WzaX/WzaS/WzaB for the respective EPS/MASC/BPS pathways.

Subsequent VS studies against the four β-barrels using a commercial fragment library resulted in the identification of compound-interaction hotspots among the extracellular loops; going forward, these sites can be targeted by inhibitors chosen to selectively bind each barrel. Potential drugs to be developed from these analyses could then incorporate one or more of the identified binding-fragment motifs. In this manner, the secretion of EPS, MASC, BPS, and PNAG could be selectively blocked in their respective organism backgrounds.

The results also suggested that the identified hotspots of WzpX and WzpS might have more affinity for negatively-charged compounds while those of WzpB and PgaA might interact better with neutral or positively-charged compounds. Also, analysis of the binding modes of the top 10 fragment poses associated to fingerprint analysis of all poses found in the hotspots revealed the key residues of each hotspot worth being explored for future drug design.

Importantly, our investigation has demonstrated the utility of using membrane-embedded integrated MD simulations to evaluate the quality of neural network-based membrane protein model structures, which are becoming widely reported in the scientific literature. In addition, our protocol will allow for the rigorous testing of more complex questions such as proposed interaction dynamics between the EPS/MASC/BPS secretion porins (WzpX/S/B) and other proteins such as modelled octamers^14^ of the WzaX/S/B Class-3 OPX proteins^6^ required for secretion of each polymer^7, 15^.

## Data and Software Availability Statement

The following files are available as supplementary materials: 1) The .pdb files of 4Y25 and the models MXAN_7418, MXAN_3226 and MXAN_1916 plus their corresponding output files obtained from CHARMM-GUI and the MOE files used to run the MD simulations; 2) The Excel files with the VS results for the four β-barrels (Tables S2-S5); 3) Grid data for the VS (Figure S1); 4) 10 best-ranked fragments from the VS (Table S6); 5) 2D interaction maps for the 10 best-ranked fragments (Figs. S1 - S40).

## CRediT authorship contribution statement

**T.C.C.F.:** Conceptualization, Writing - original draft, Formal analysis, Writing - review & editing, Methodology

**F.S.:** Writing - original draft

**A.A.:** Software, Formal analysis

**S.T.I.:** Conceptualization, Funding acquisition, Supervision, Writing - original draft, Writing - review & editing

**S.R.L.:** Conceptualization, Funding acquisition, Supervision, Writing - review & editing.

## Acknowledgments

The authors wish to thank the Institut National de la Recherche Scientifique (INRS) and Prof Teodorico Ramalho from the Federal University of Lavras - Brazil, for software availability, and Calcul Québec for the availability of the supercomputer.

