## Supplementary material for "Molecular dynamics of outer membrane-embedded polysaccharide secretion porins reveals closed resting-state surface gates targetable by virtual fragment screening for drug hotspot identification": Suplementary material

Steven R. LaPlante^1,2,^*

^1^	Institut National de la Recherche Scientifique (INRS), Centre Armand-Frappier Santé Biotechnologie, Université du Québec, Institut Pasteur International Network, Laval, QC, H7V 1B7, Canada.

^2^	PROTEO, the Quebec Network for Research on Protein Function, Engineering, and Applications, Université Laval, Quebec, QC, G1V 0A6, Canada.

^3^	Laboratory of Molecular Modeling Applied to Chemical and Biological Defense, Military Institute of Engineering, Rio de Janeiro, 22290-270, Brazil.

^4^	Department of Chemistry, Faculty of Science, University of Hradec Kralove, Rokitanskeho 62, 50003, Hradec Kralove, Czech Republic.

^5^	Chemical Computing Group, Montreal Quebec H3A 2R7, Canada.

**Supplementary Material**

**Table S1.** Grid data for the VS.

| Protein | Grid Coordinates (Å) | | | Radius (Å) |
| --- | --- | --- | --- | --- |
|  | X | Y | Z |  |
| WzpX (model) | -2.56 | 6.17 | 17.49 | 25.00 |
| WzpS (model) | 1.19 | 4.58 | 15.49 | 26.81 |
| WzpB (model) | -3.31 | 0.42 | 26.81 | 32.00 |
| PgaA (PDB 4Y25) | -3.31 | 0.42 | 30.24 | 26.81 |

**Table S6**. Best-ranked fragments for each porin following VS. The MolDockScores are between (), H-bond residues are between [] and pi-stacking residues are between {}. Common fragments between WzpX and WzpS are shown in yellow cells, between WzpS and PgaA are in green cells, and between WzpB and PgaA are in blue cells.

| **#** | **WzpX** | **WzpS** | **WzpB** | **PgaA** |
| --- | --- | --- | --- | --- |
| 1 | 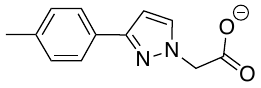 | 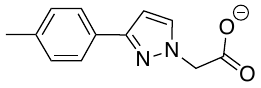 | 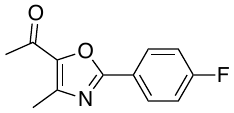 | 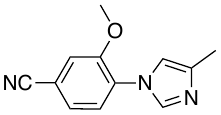 |
|  | PS-3621 (-221.77*) | PS-3621 (-203.53*) | 9T-0617 (-110.60*) | SS-4784 (-107.42*) |
|  | (Ser170, Arg305) | -- | (Pro39, Thr40, Ser179) | (Arg787, Pro788, Tyr789) |
| 2 | 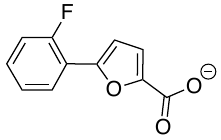 | 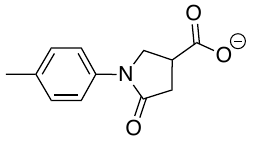 | 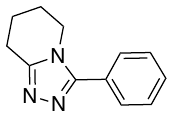 | 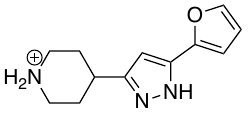 |
|  | PS-6398 (-215.21*) | AS-5481 (202.41) | 5B-098 (-110.55*) | 9L-711 (-104.31*) |
|  | (Asp118, Asp304, Arg305) | (Arg60) | (Phe37, Ser179 | (Lys754, Pro788) |
| 3 | 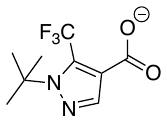 | 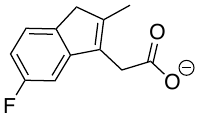 | 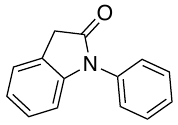 | 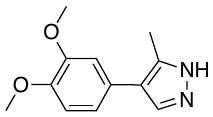 |
|  | 11W-0279 (-212.07*) | SS-2964 (-200.78*) | 12W-0033 (-110.16*) | 1G-004 (-103.44*) |
|  | (Ser108, Arg305) | (Arg103) | (Pro39) | -- |
| 4 | 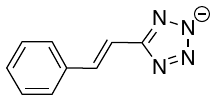 | 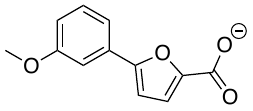 | 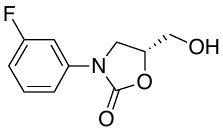 | 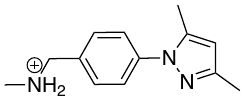 |
|  | 11T-0832 (-211.88*) | PS-5909 (-200.46*) | FS-2562 (-110.09*) | PS-3773 (-102.30*) |
|  | (Asp118, Arg305) | (Arg60) | (Thr40) | (Pro716, Lys754) |
| 5 | 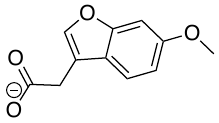 | 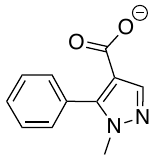 | 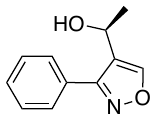 | 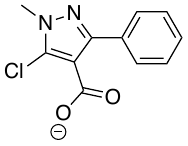 |
|  | PS-4406 (-208.66*) | PS-3429 (-199.41) | PS-5035 (-109.97*) | 3K-528S (-101.62*) |
|  | (Thr104, Thr116, Arg305) | (Arg21, Arg347) | (Pro39, Ala98, Leu177) | (Lys718, Lys754, Tyr789, Asp790) |
| 6 | 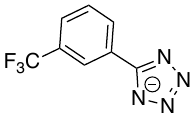 | 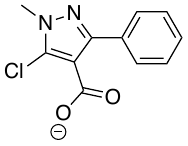 | 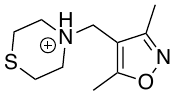 | 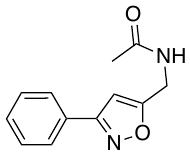 |
|  | 9W-0332 (-207.35*) | 3K-528S (-199.23) | 12L-510S (-108.99*) | 7M-374S (-101.60*) |
|  | (Gly115, Arg305) | (Gly105, Arg108) | (Tyr36, Ser179) | (Arg787, Pro788, Tyr789) |
| 7 | 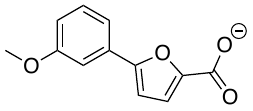 | 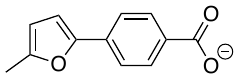 | 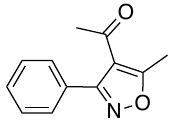 | 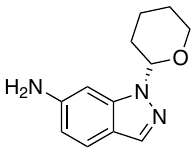 |
|  | PS-5909 (-207.19*) | PS-5916 (-197.97) | AB-0705 (-108.67*) | PS-5133 (-101.54*) |
|  | (Leu109, Arg305) | (Arg15, Arg60, Thr110, Arg347) | (Ser179) | (Tyr789, Asp790) |
| 8 | 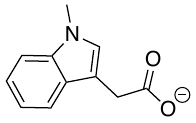 | 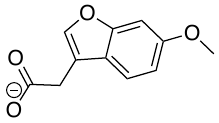 | 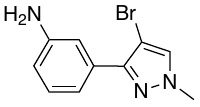 | 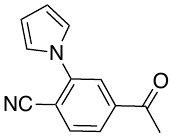 |
|  | 7L-904 (-206.00*) | PS-4406 (-197.62*) | PS-5055 (-108.61*) | 2T-1515 (-101.02*) |
|  | (Asp112, Thr116, Arg305) | (Arg103) | (Gly33, Val34) | (Arg787, Pro788) |
| 9 | 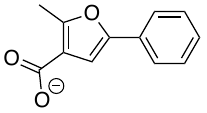 | 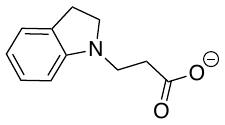 | 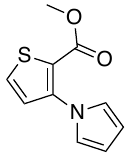 | 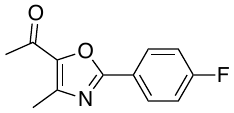 |
|  | FS-1464 (-205.99*) | PS-6198 (-197.32) | 1R-0138 (-108.49*) | 9T-0617 (-100.61*) |
|  | (Arg305) | -- | (Asn95, Ser179) | (Arg787) |
| 10 | 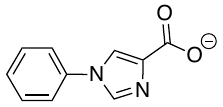 | 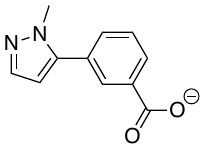 | 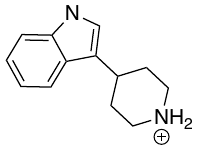 | 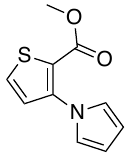 |
|  | PS-2990 (-205.87*) | FS-3666 (-197.07) | FS-1763 (-107.75*) | 1R-0138 (-100.20*) |
|  | (Thr104, Asp304, Arg305) | (Asn253) | (Thr40, Asn95) | (Gln753, Tyr789, Asp790) |

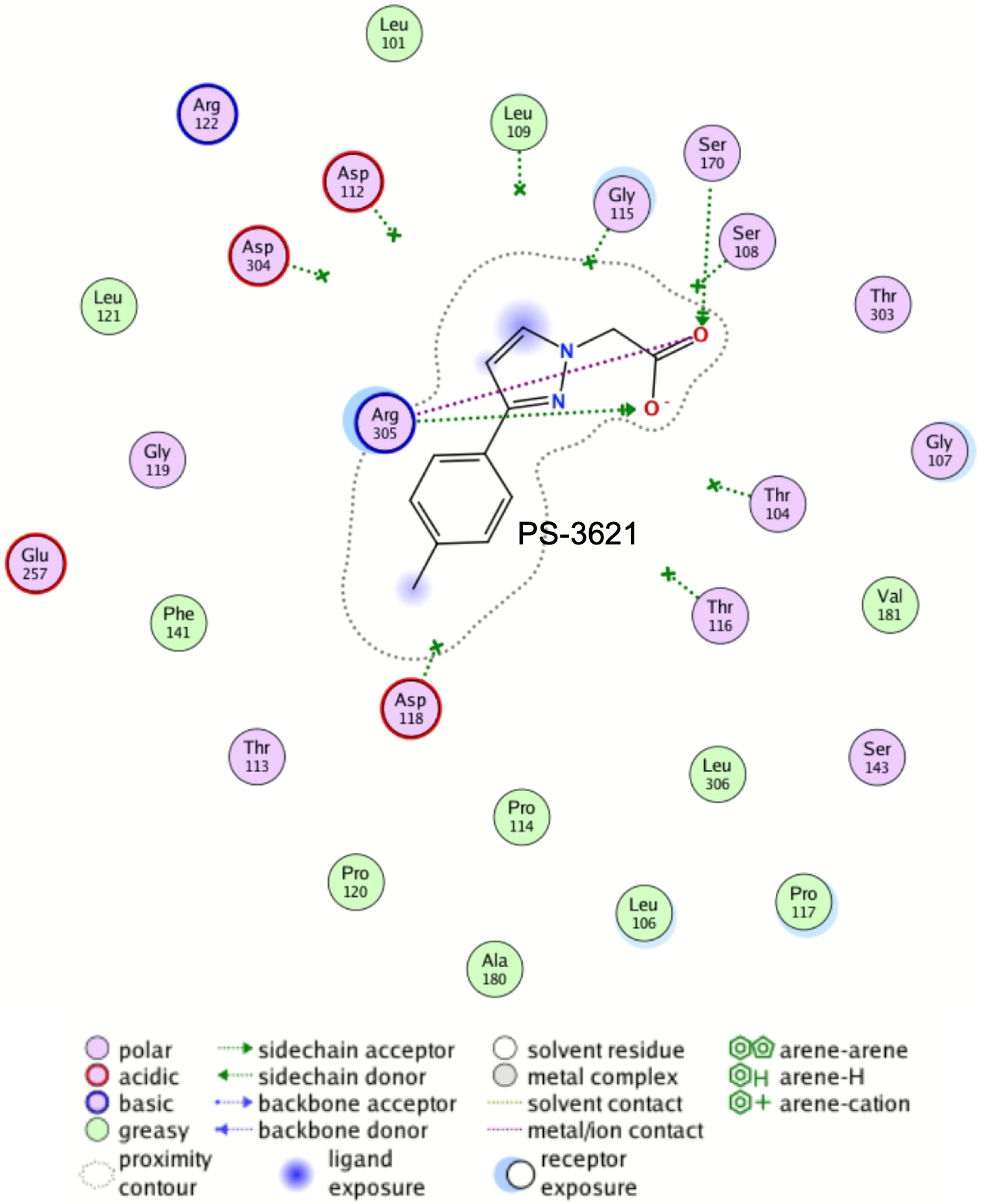

**Fig. S1.** 2D interaction map of PS-3621 in the main hotspot of WzpX.

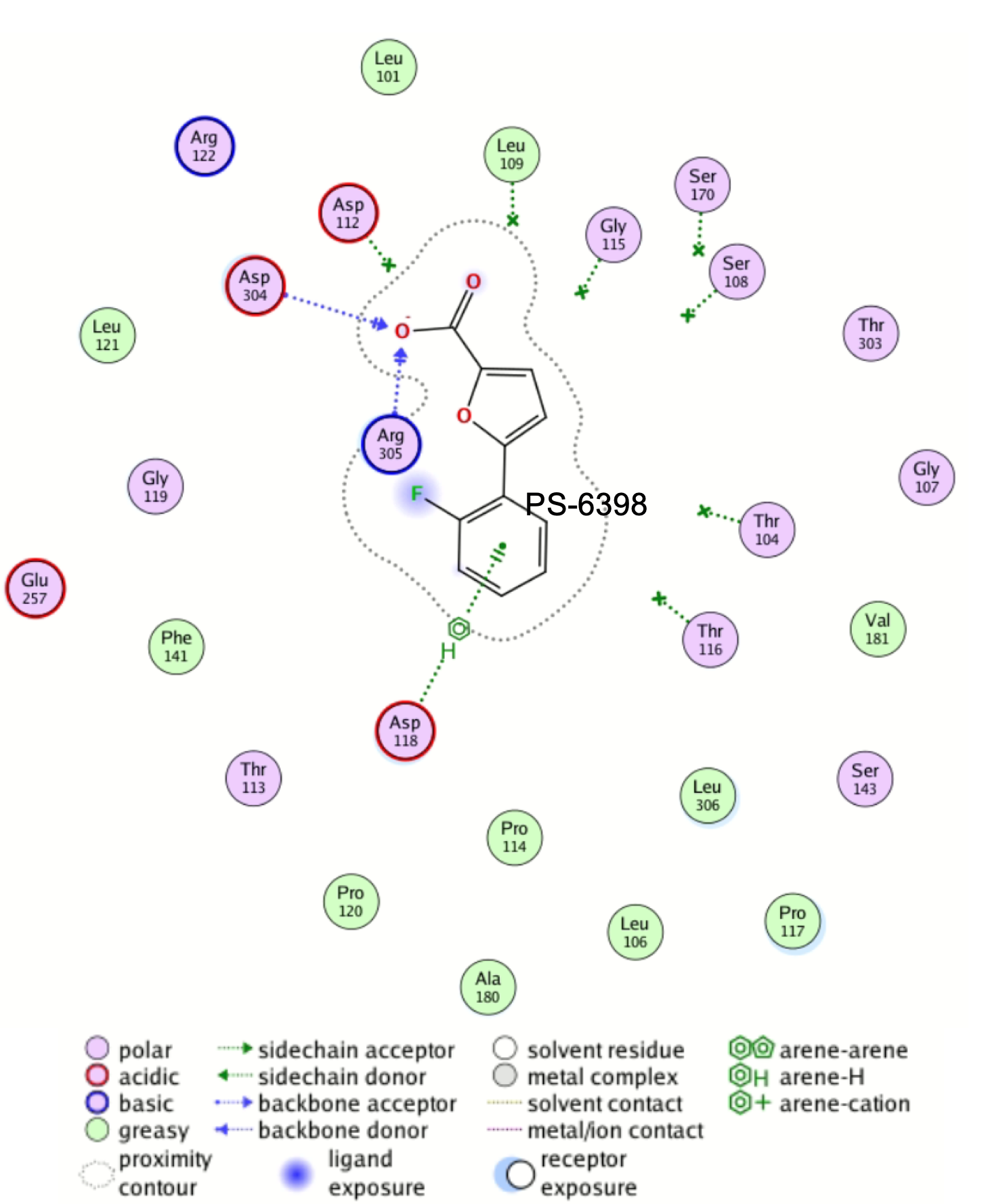

**Fig. S2.** 2D interaction map of PS-6398 in the main hotspot of WzpX.

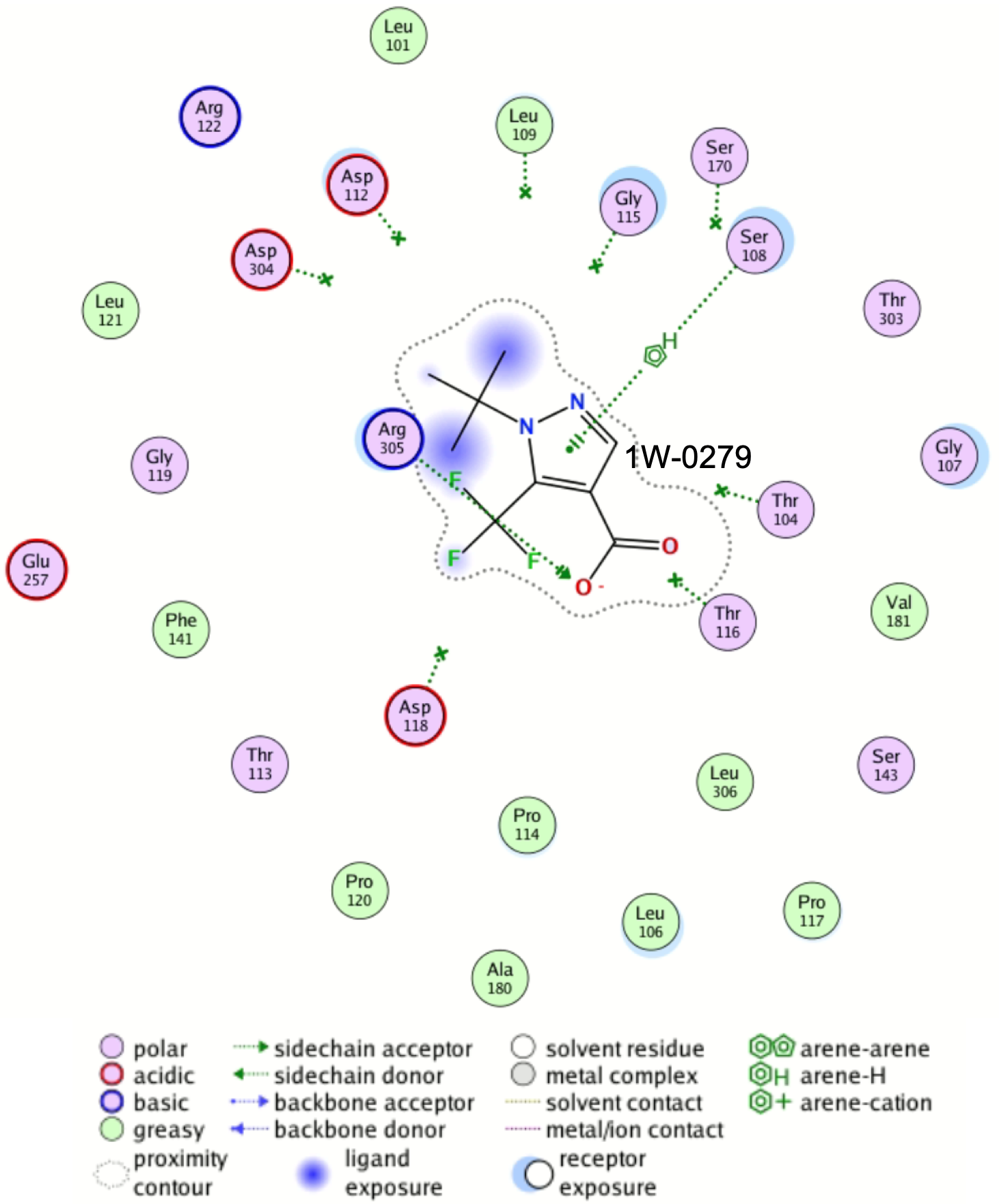

**Fig. S3.** 2D interaction map of 11W-0279 in the main hotspot of WzpX.

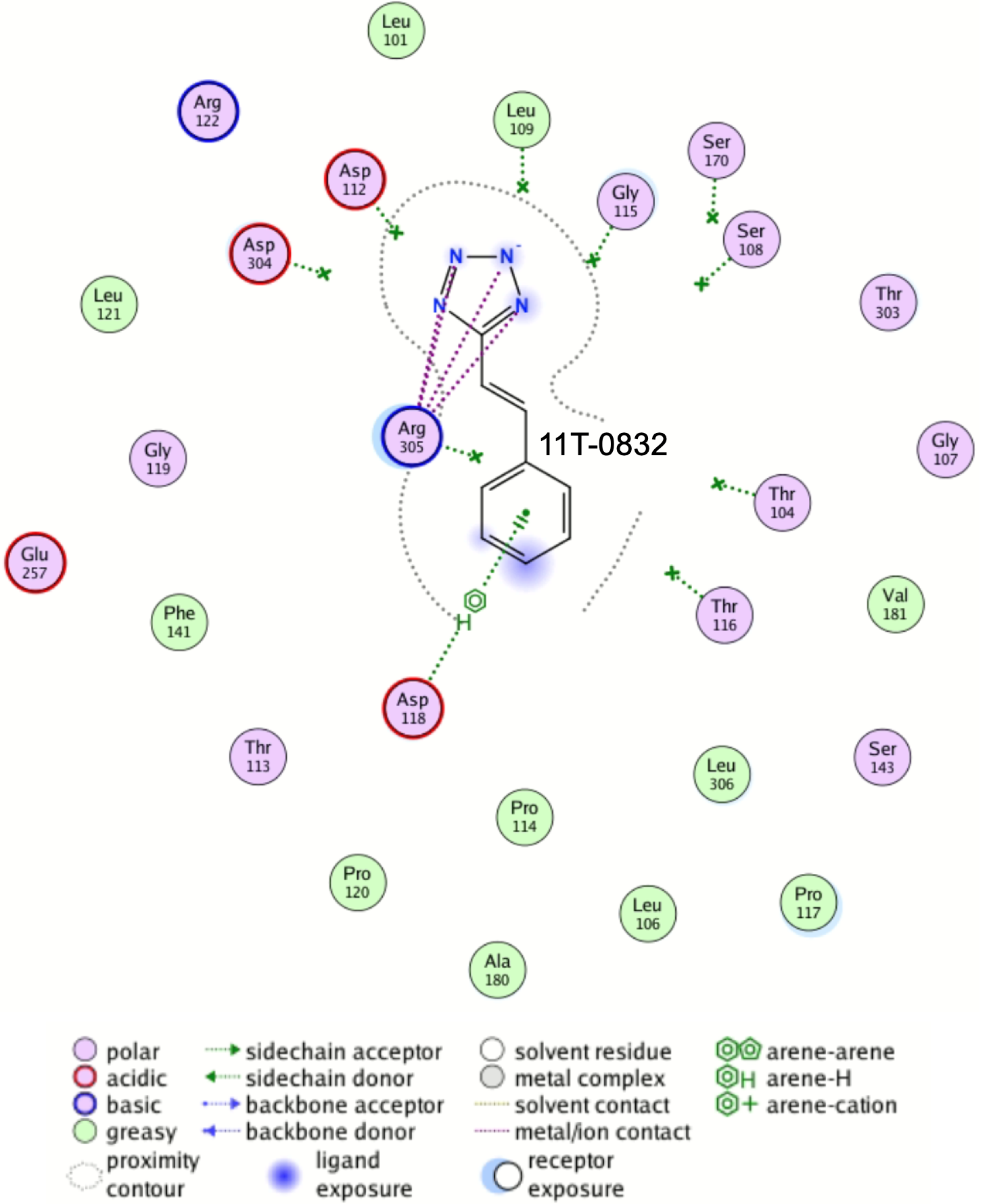

**Fig. S4.** 2D interaction map of 11T-0832 in the main hotspot of WzpX.

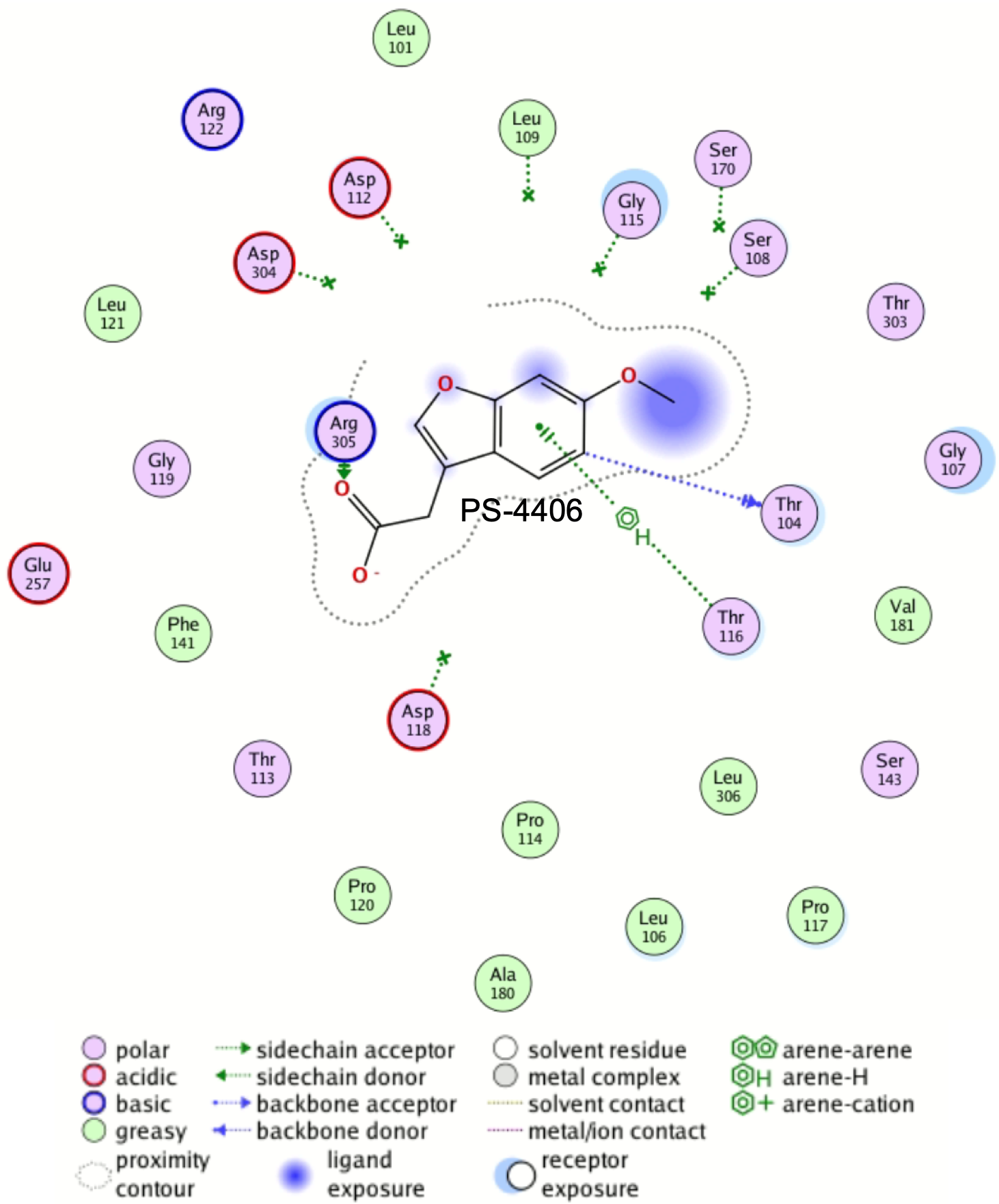

**Fig. S5.** 2D interaction map of PS-4406 in the main hotspot of WzpX.

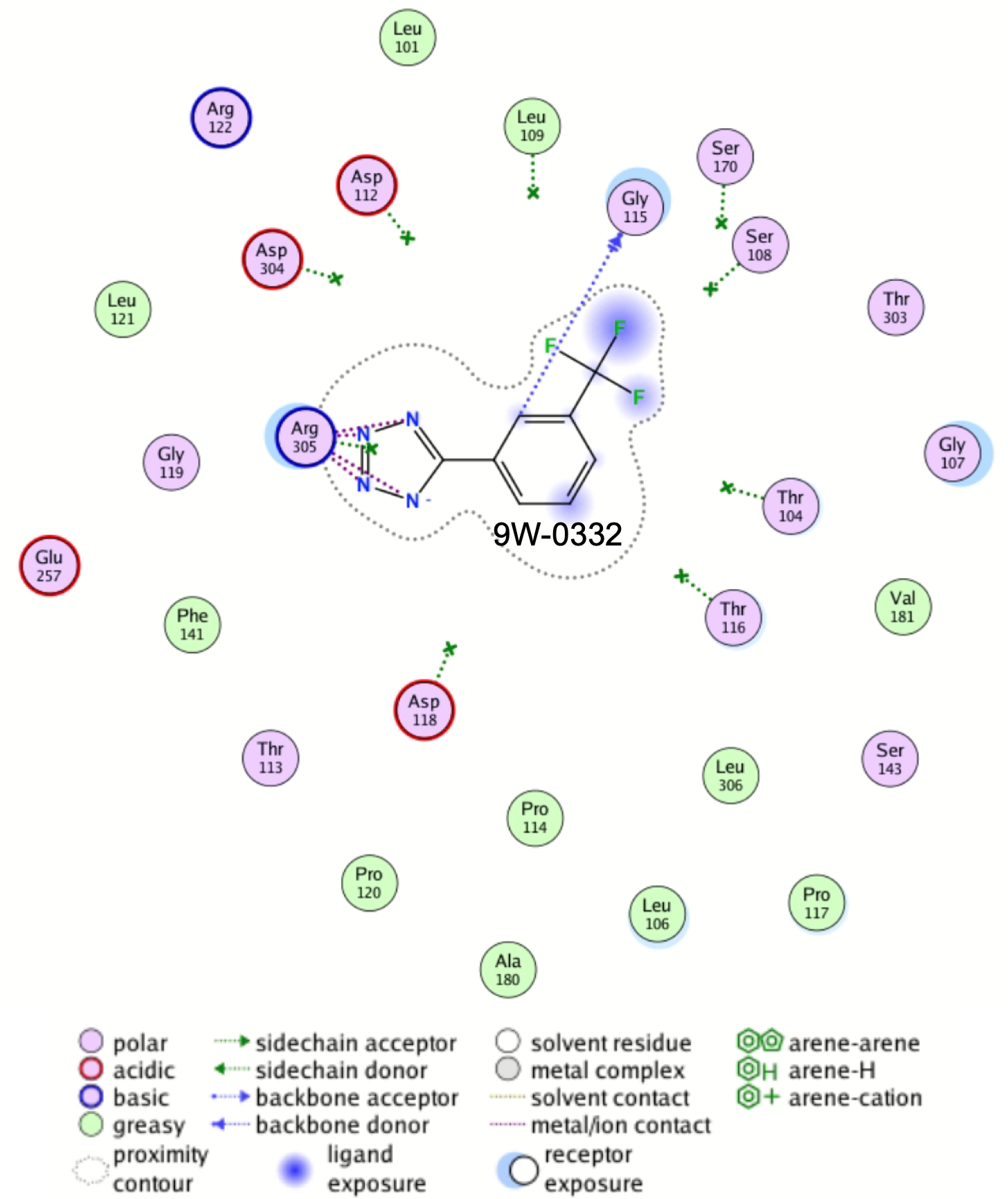

**Fig. S6.** 2D interaction map of 9W-0332 in the main hotspot of WzpX.

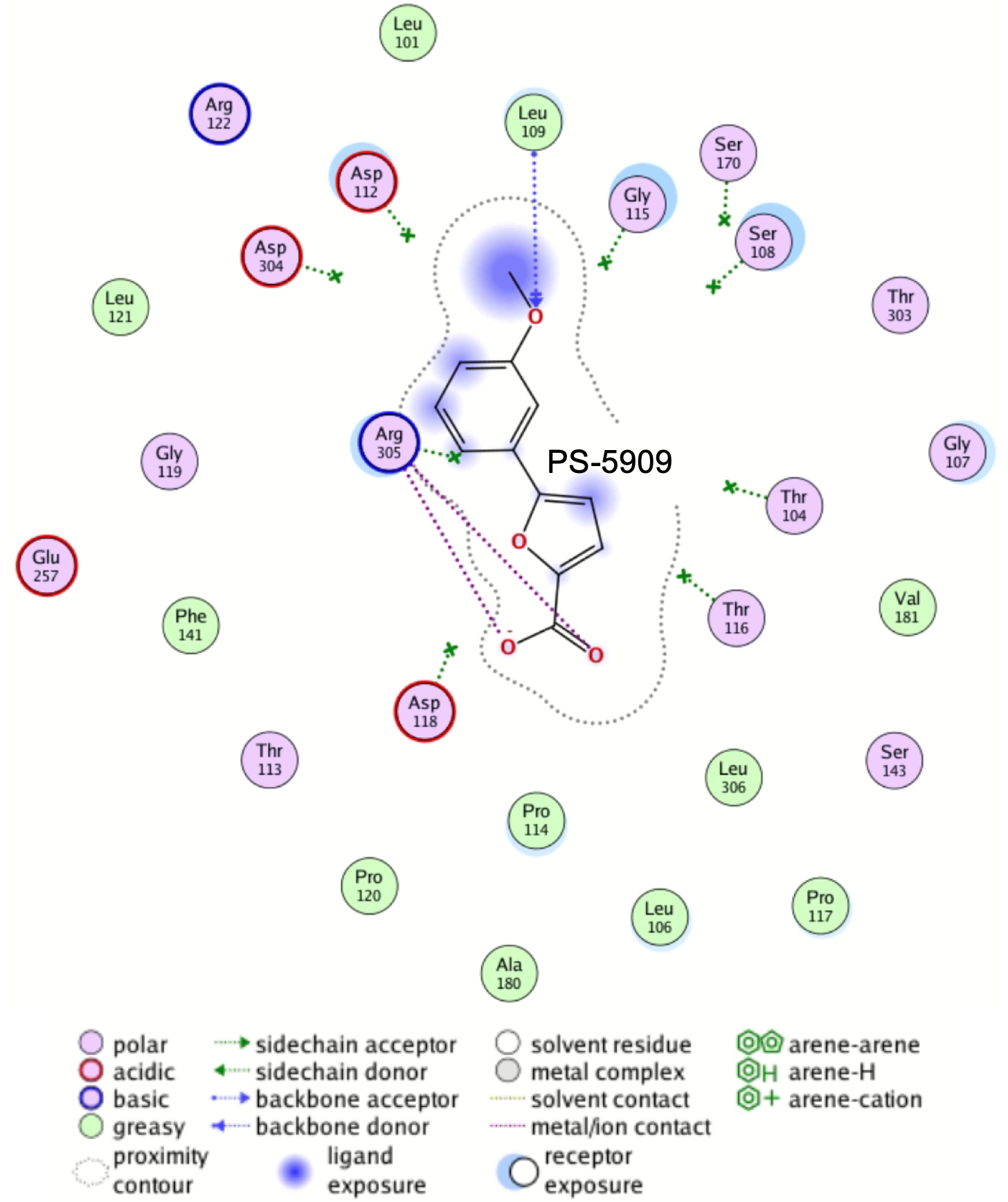

**Fig. S7.** 2D interaction map of Ps-5909 in the main hotspot of WzpX.

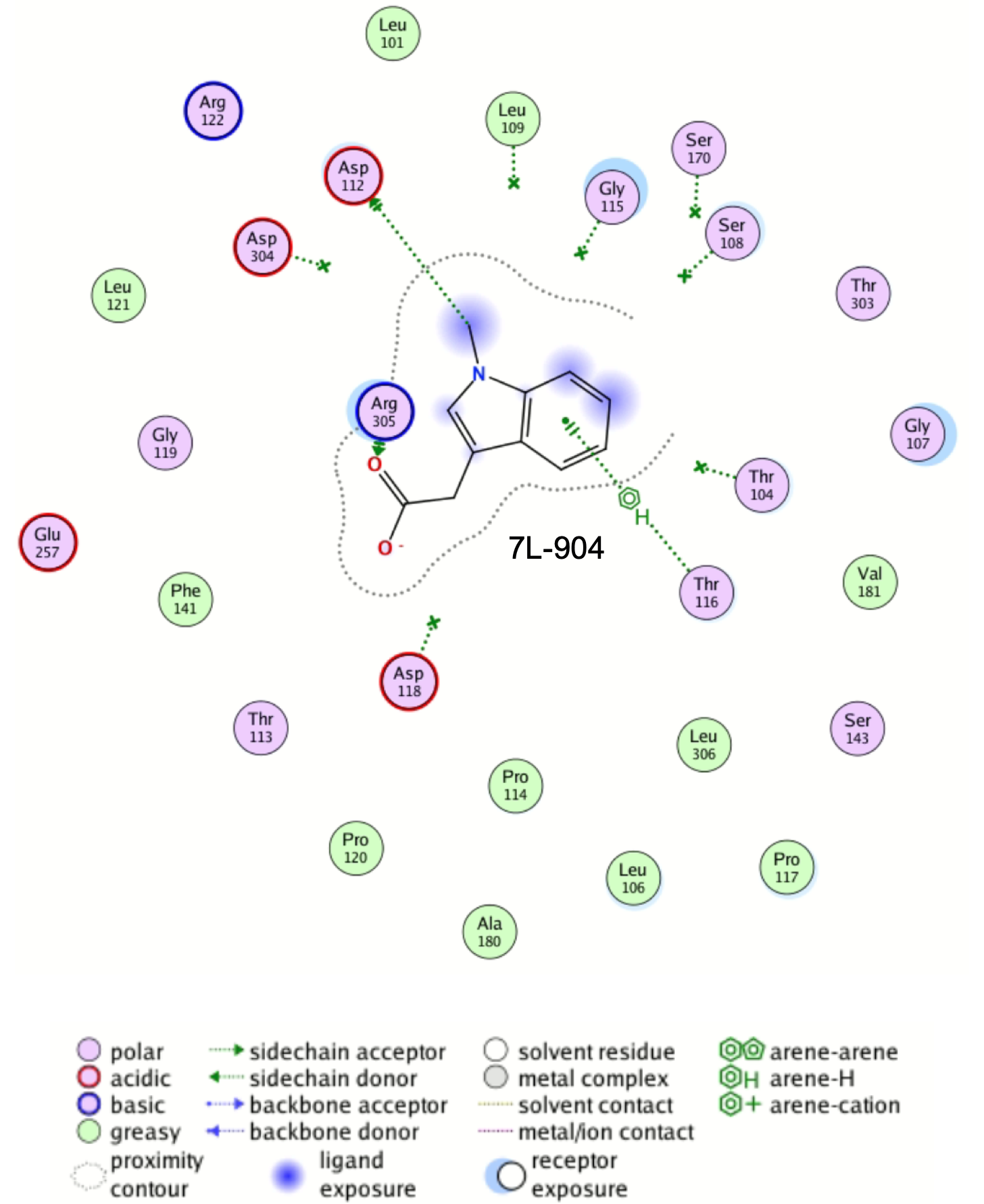

**Fig. S8.** 2D interaction map of 7L-904 in the main hotspot of WzpX.

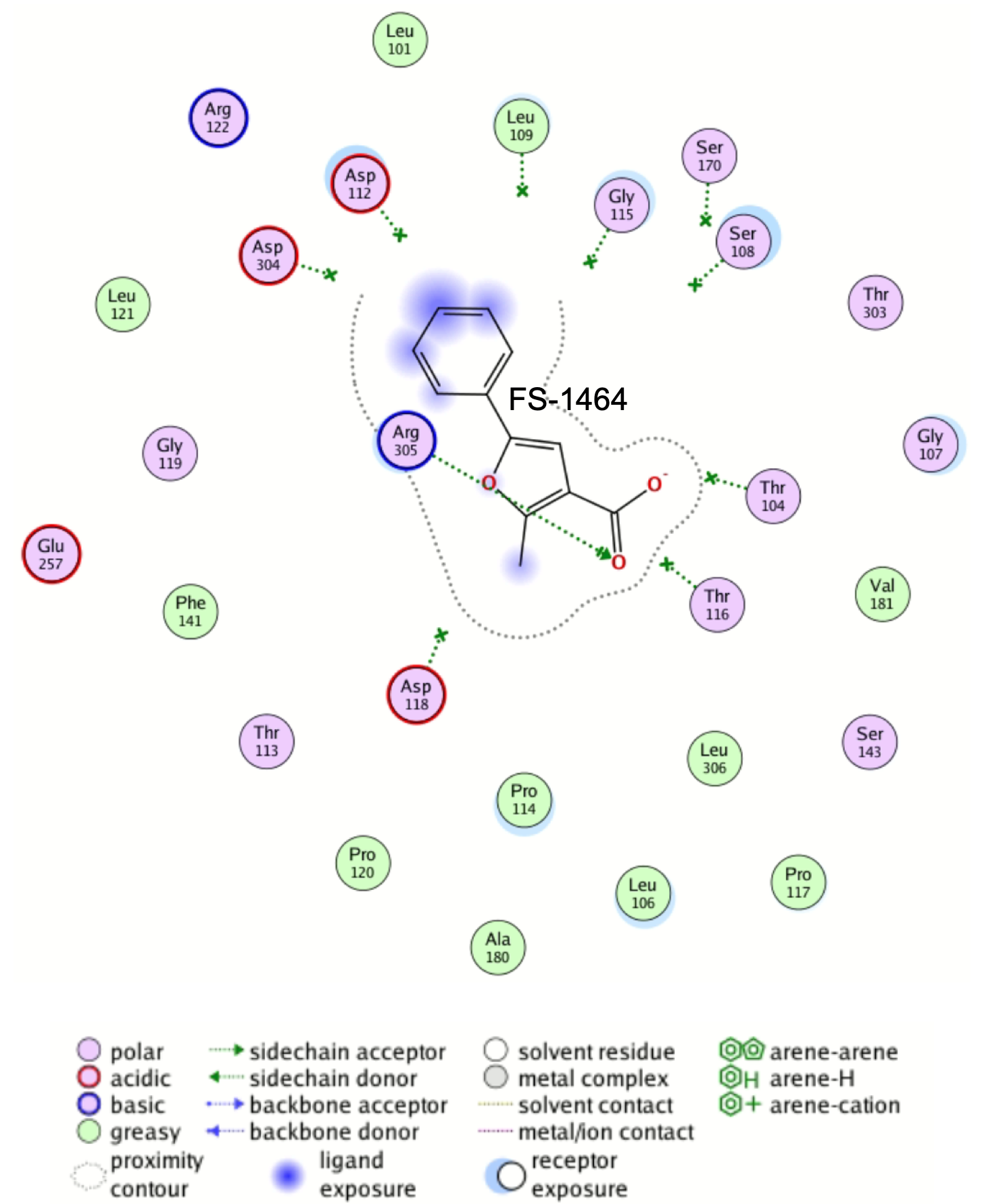

**Fig. S9.** 2D interaction map of FS-1464 in the main hotspot of WzpX.

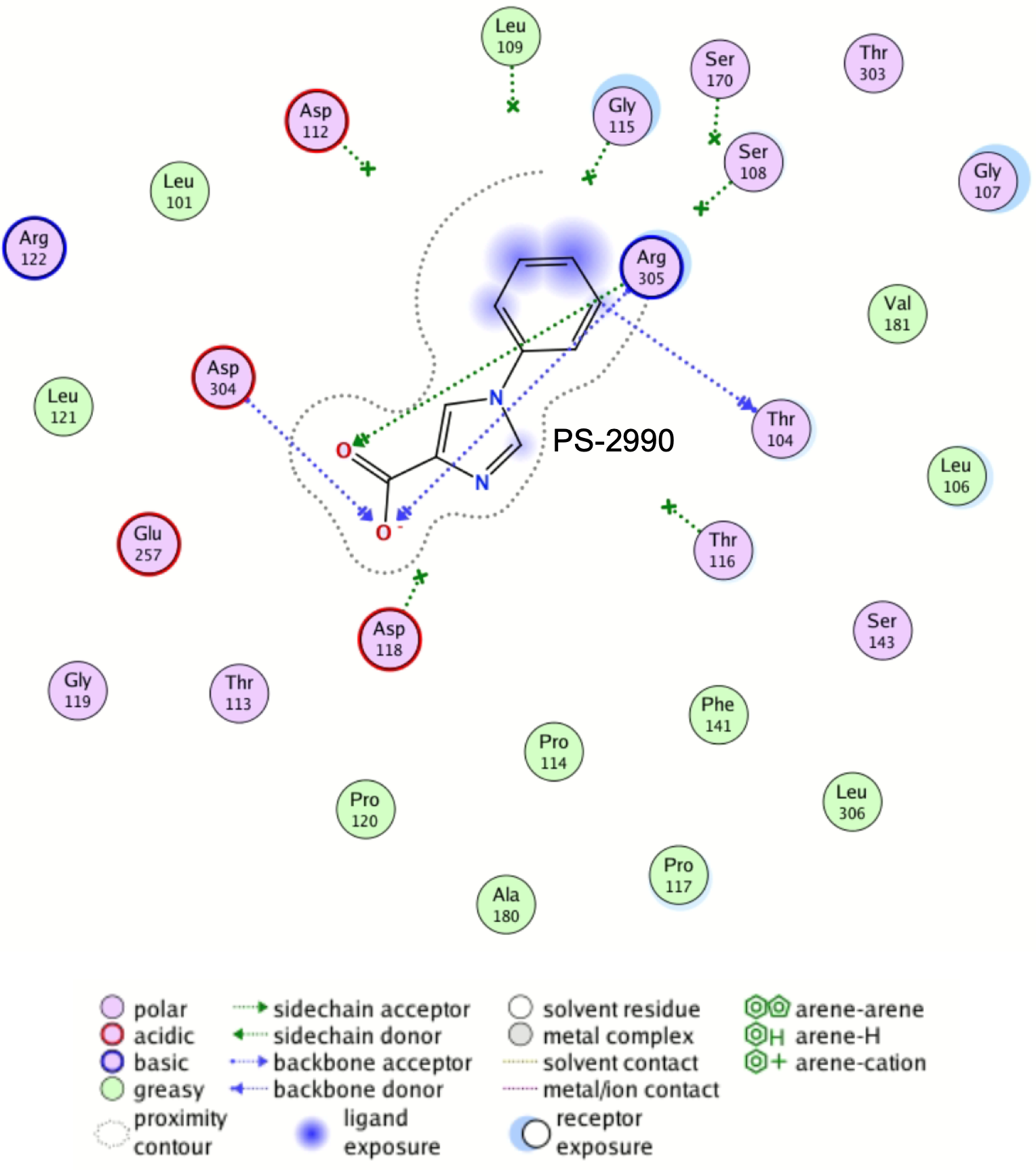

**Fig. S10.** 2D interaction map of PS-2990 in the main hotspot of WzpX.

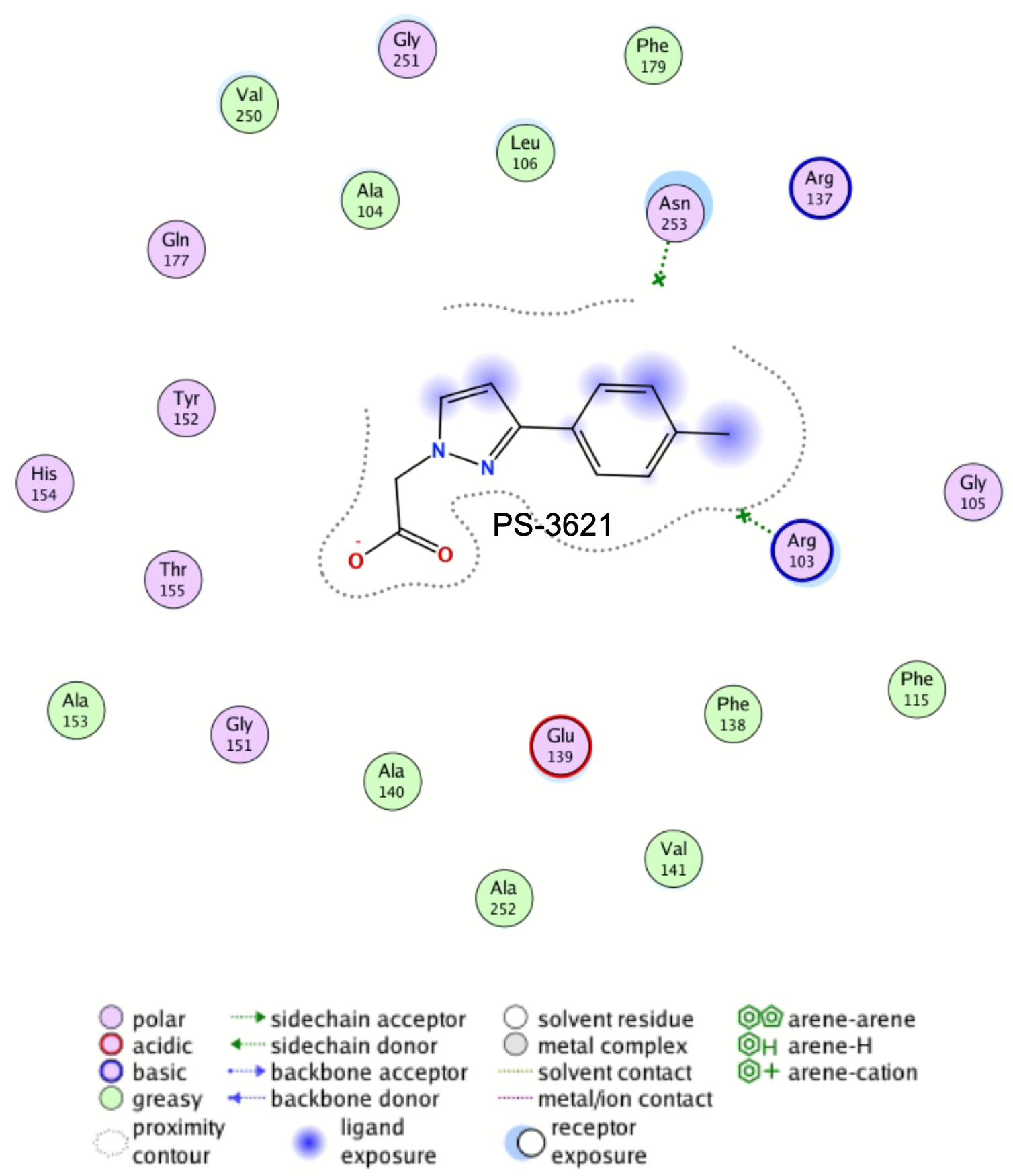

**Fig. S11.** 2D interaction map of PS-3621 in one of the main hotspots of WzpS.

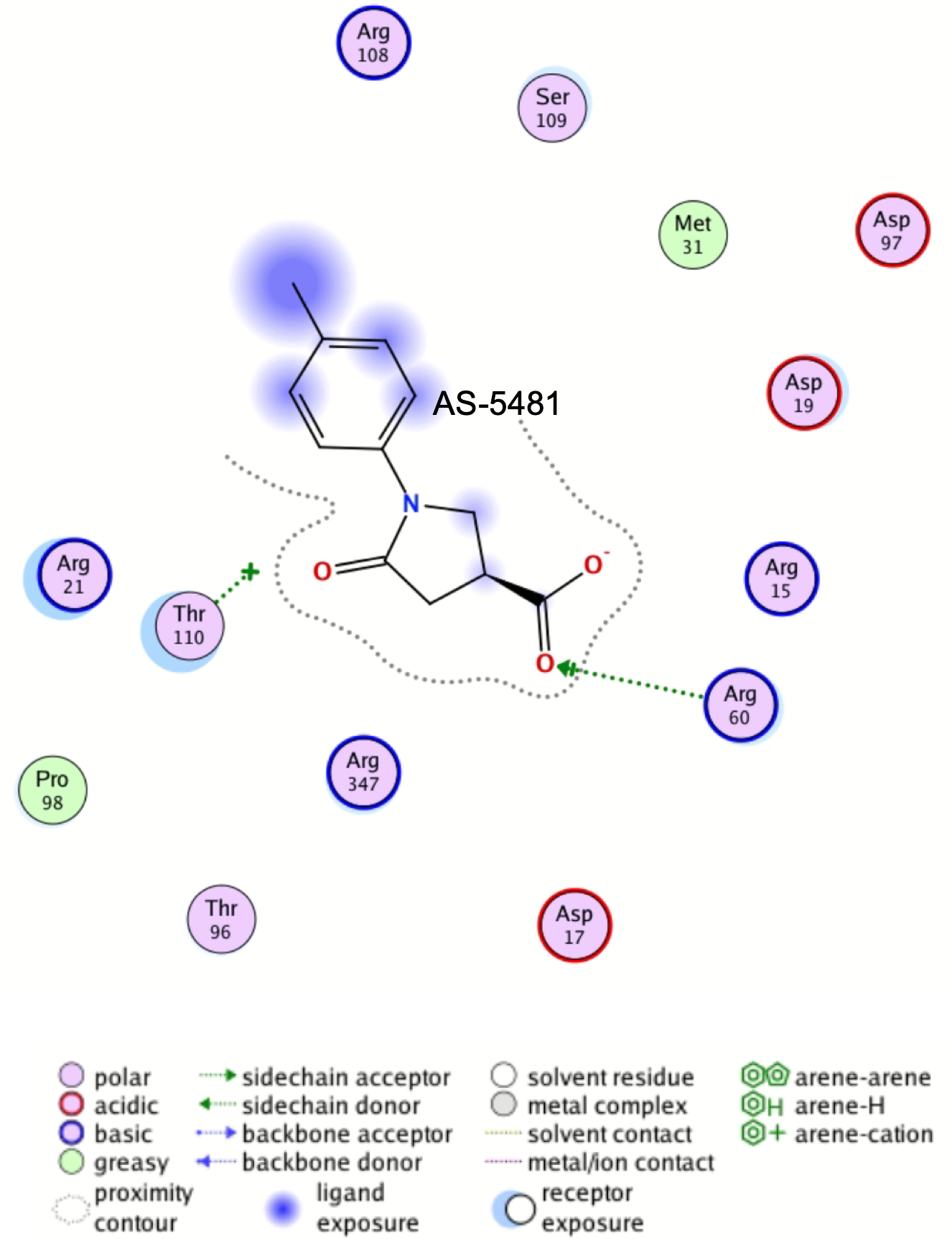

**Fig. S12.** 2D interaction map of AS-5481 in one of the main hotspots of WzpS.

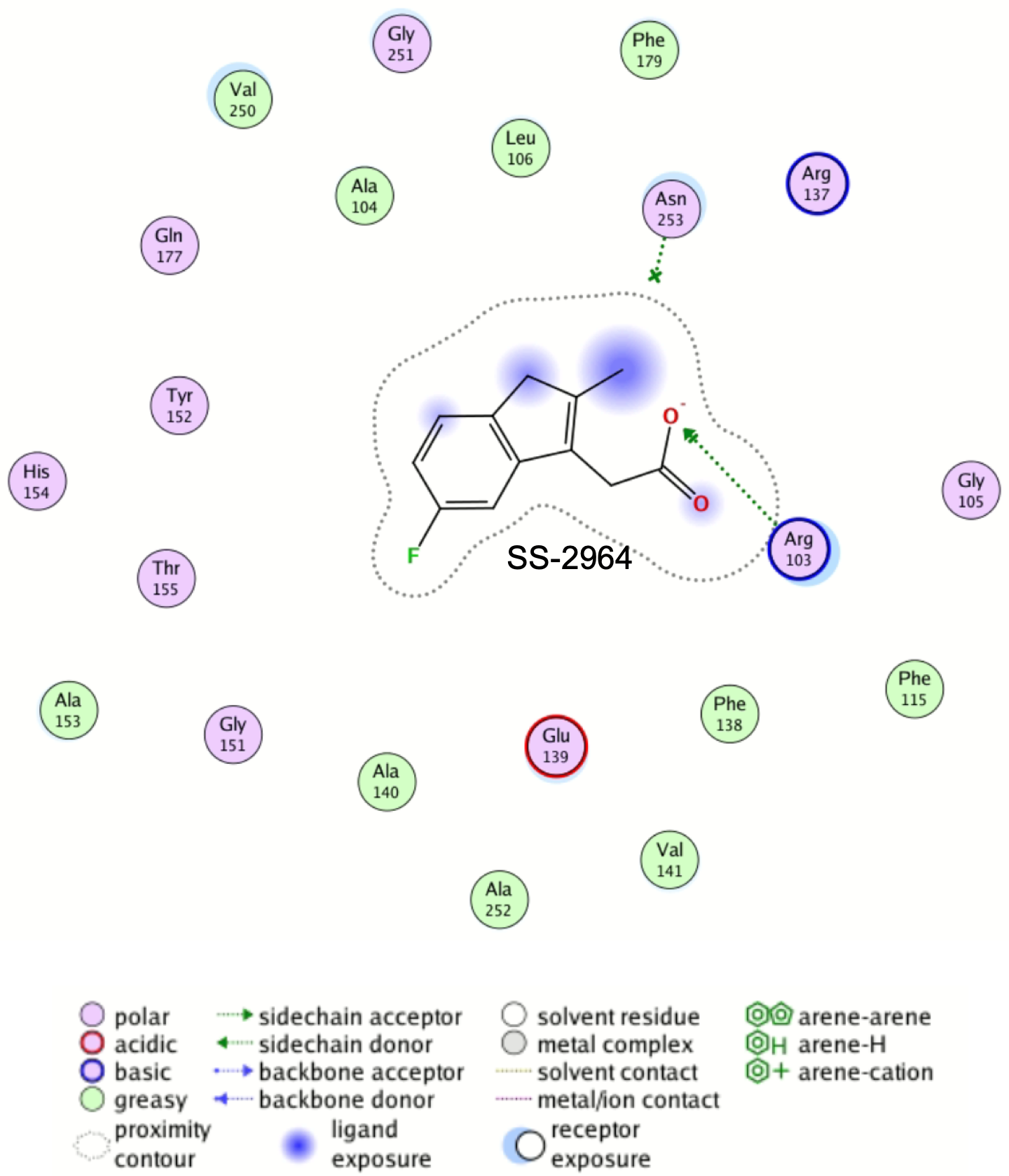

**Fig. S13.** 2D interaction map of SS-2964 in one hotspot of WzpS.

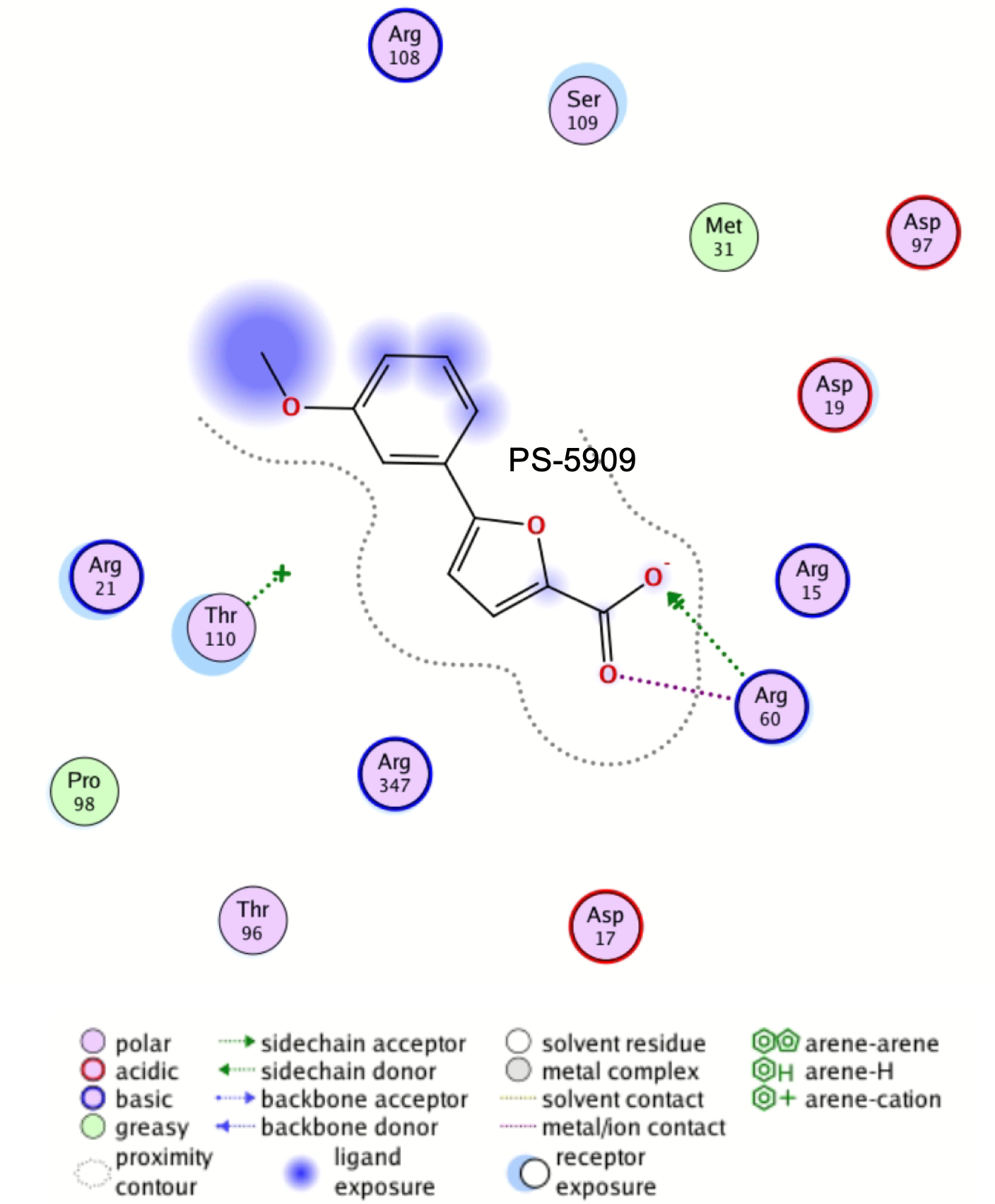

**Fig. S14.** 2D interaction map of PS-5909 in one hotspot of WzpS.

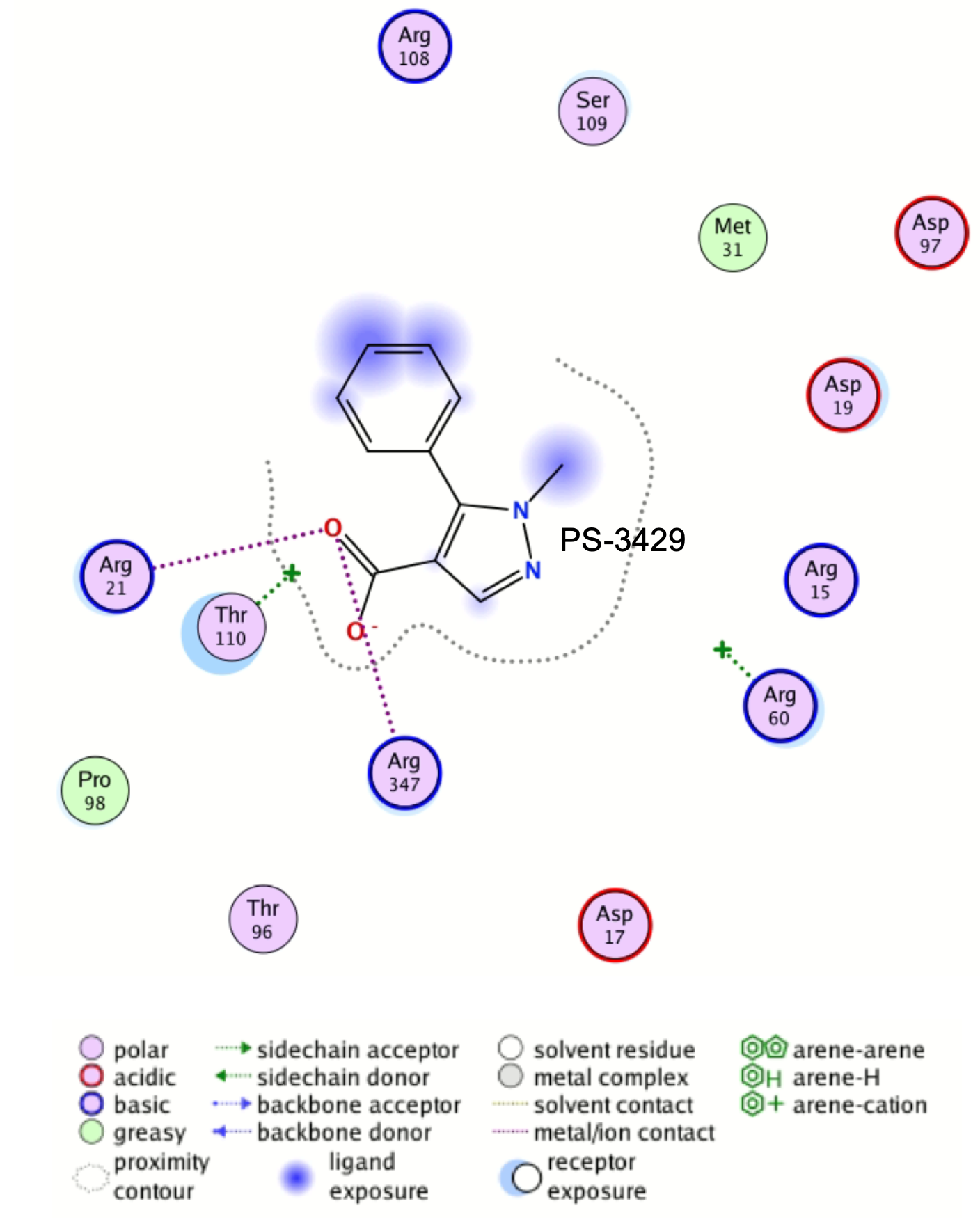

**Fig. S15.** 2D interaction map of PS-3429 in one hotspot of WzpS.

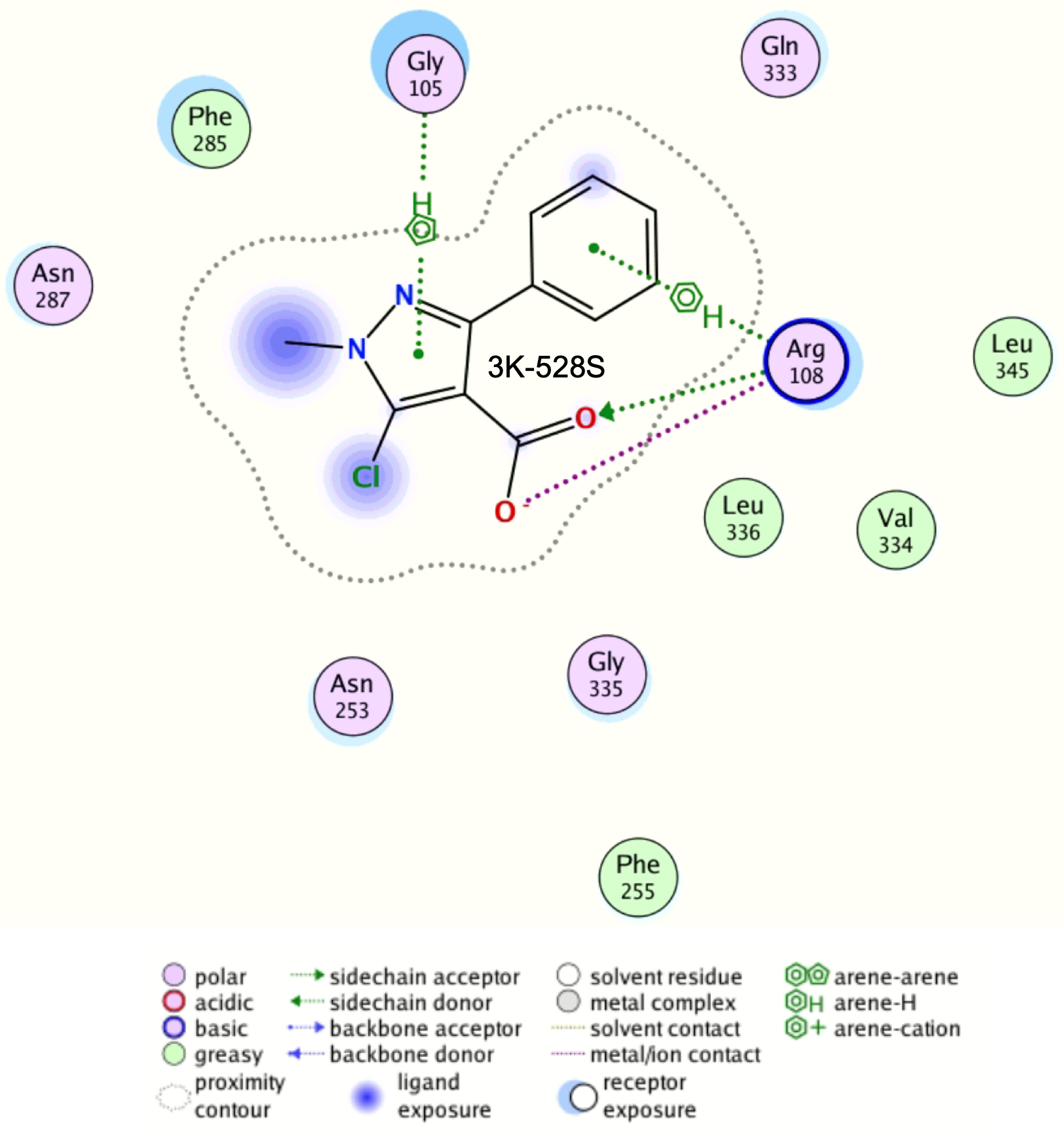

**Fig. S16.** 2D interaction map of 3K-528S in the main hotspot of WzpS.

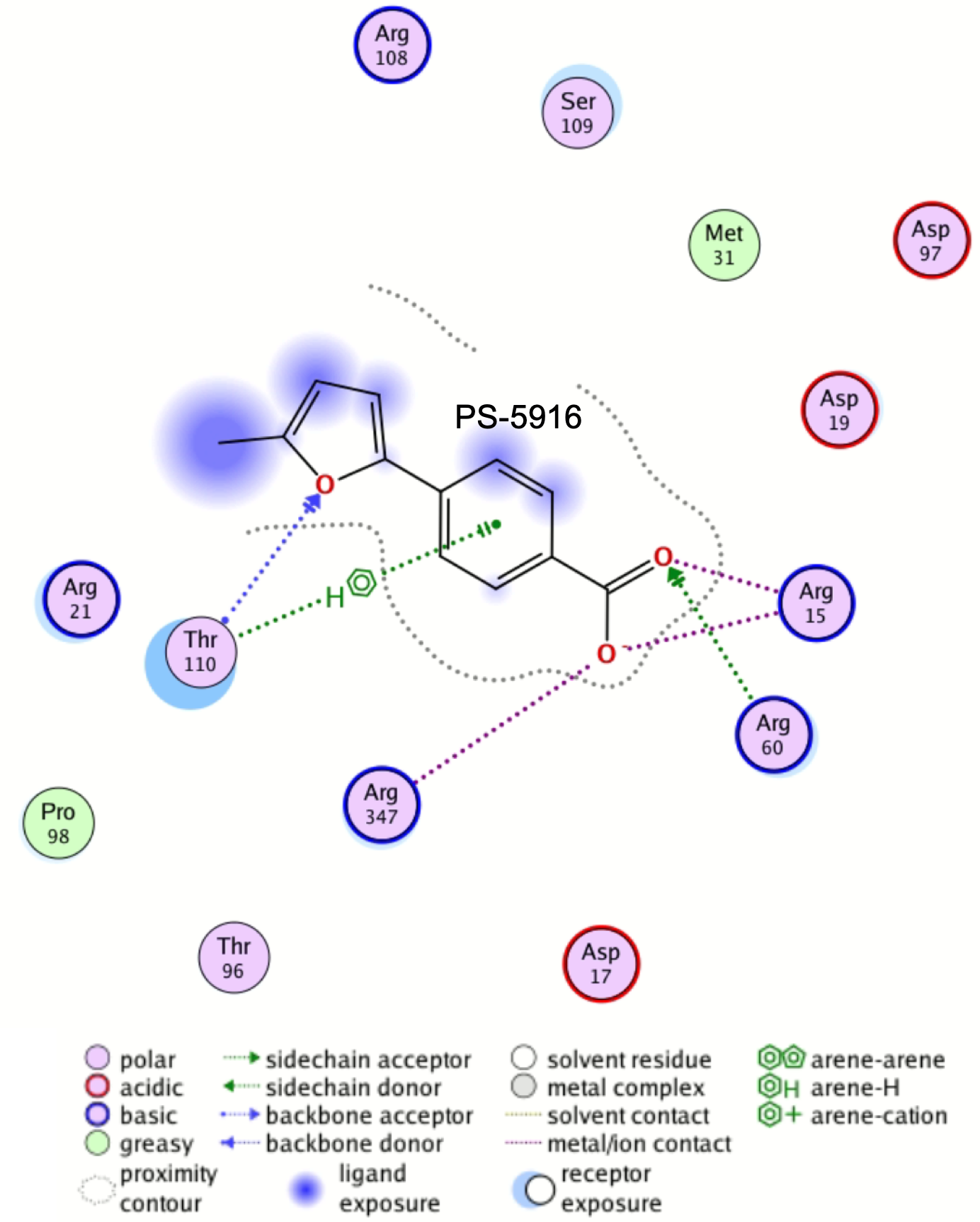

**Fig. S17.** 2D interaction map of PS-5916 in the main hotspot of WzpS.

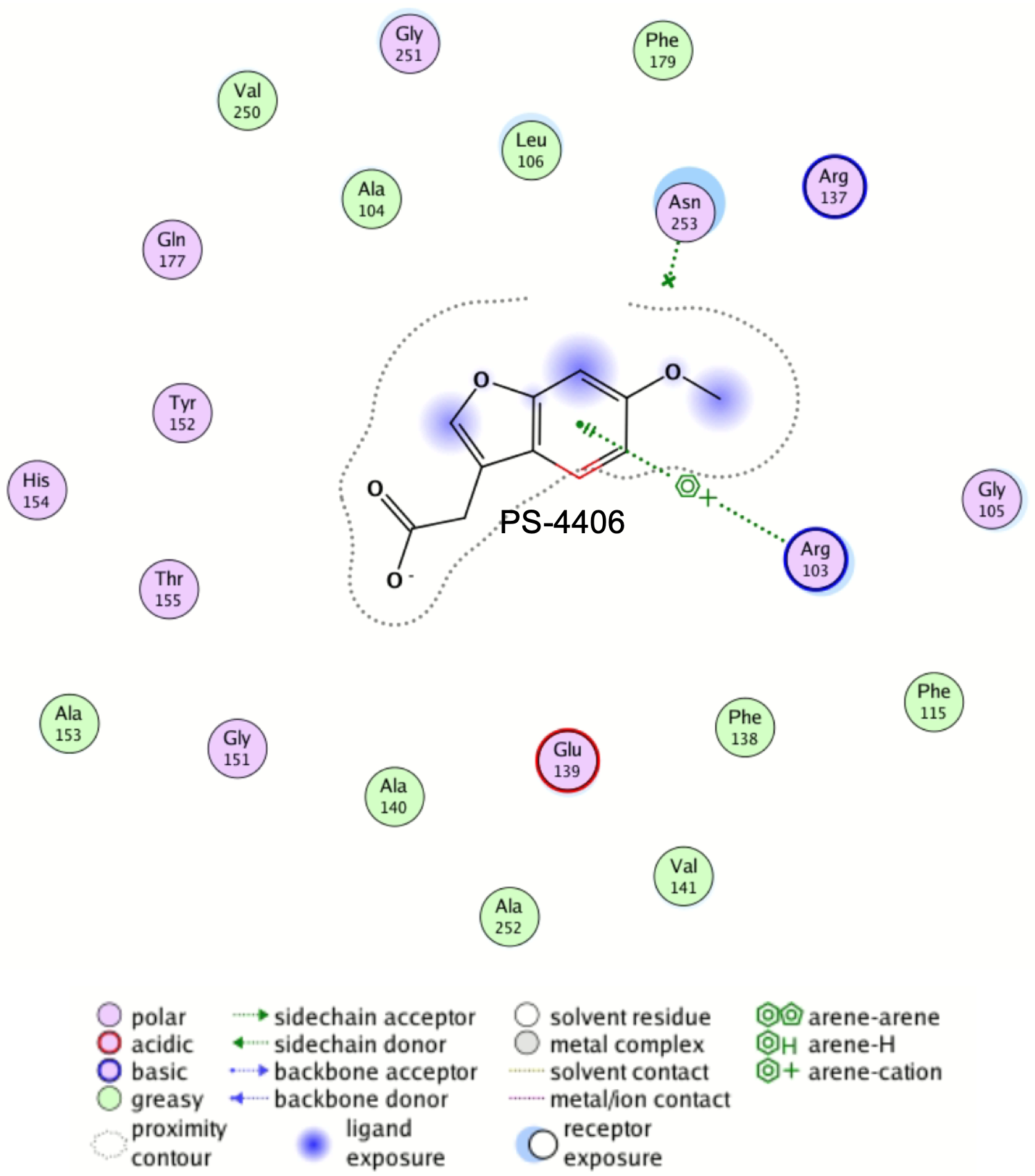

**Fig. S18.** 2D interaction map of PS-4406 in one of the hotspot of WzpS.

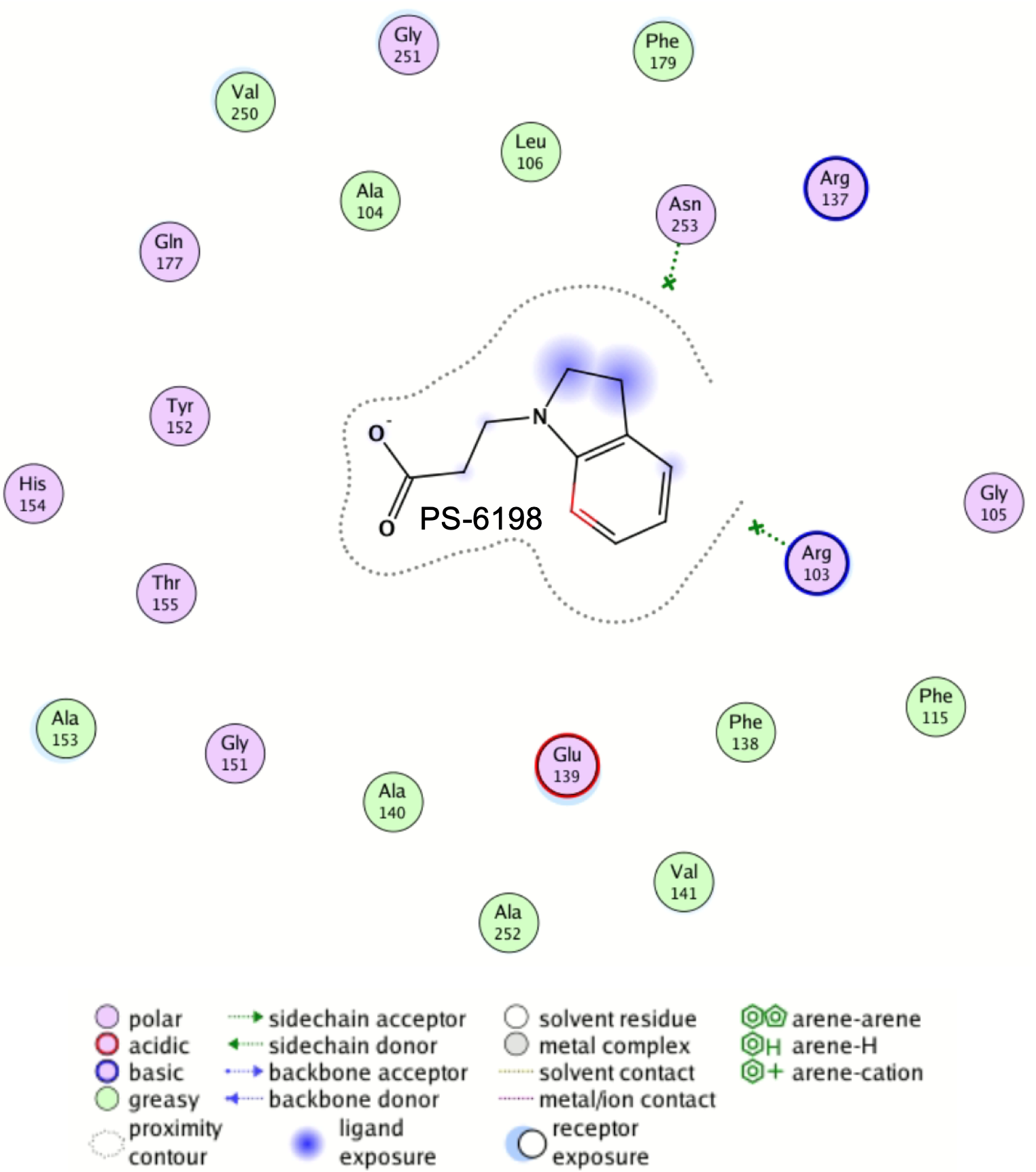

**Fig. S19.** 2D interaction map of PS-6198 in one of the hotspot of WzpS.

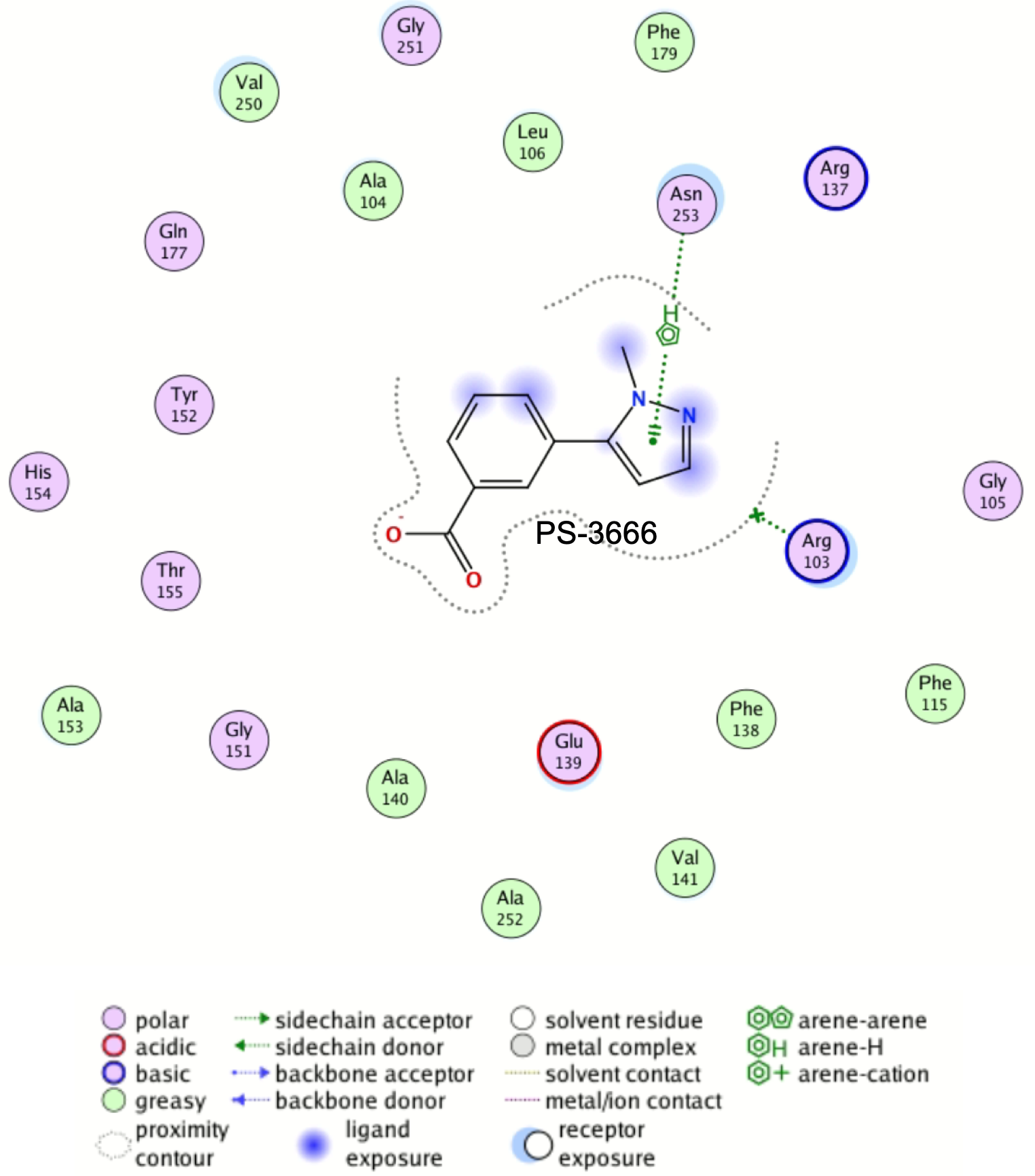

**Fig. S20.** 2D interaction map of PS-3666 in one of the hotspots of WzpS.

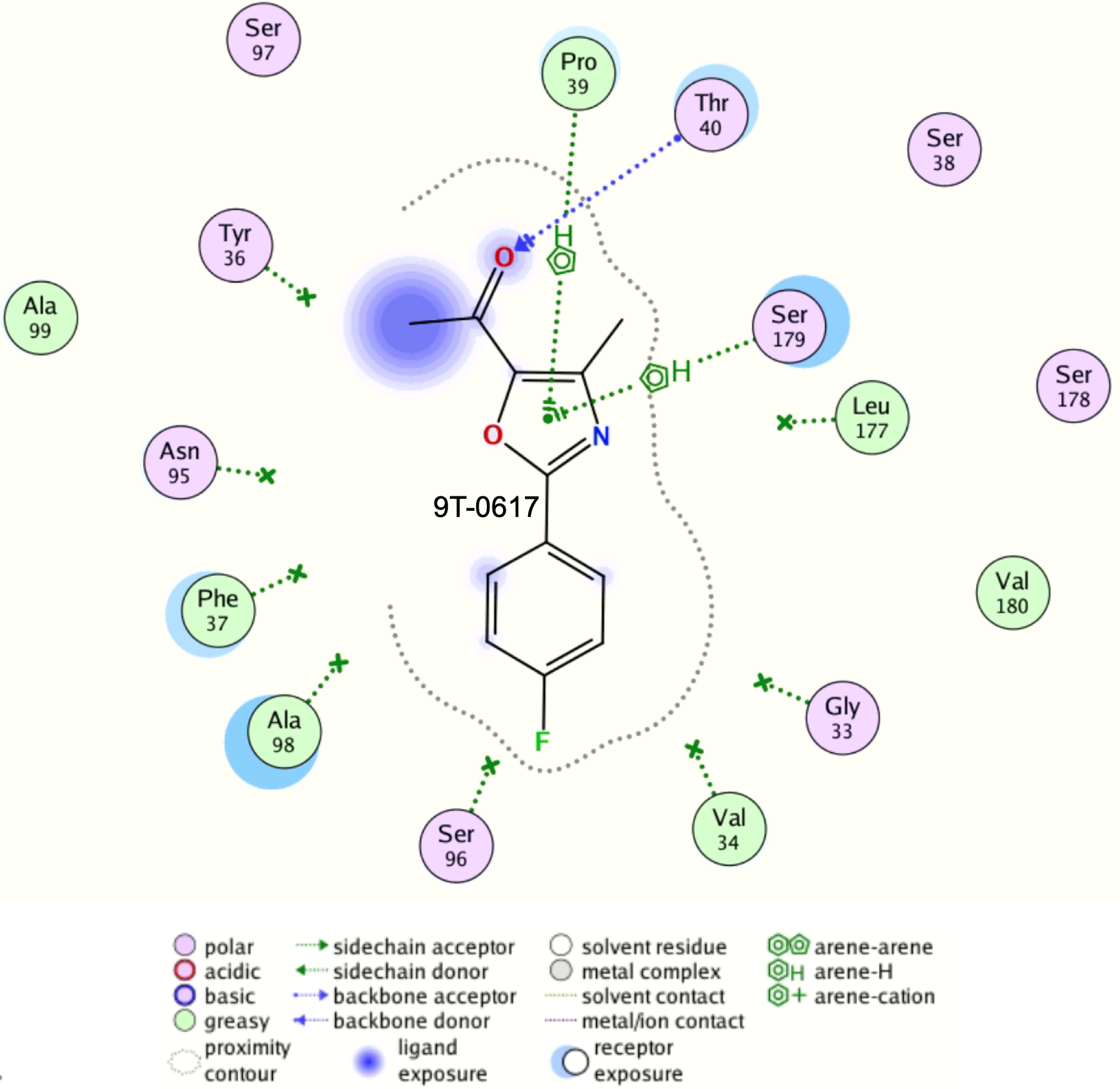

**Fig. S21.** 2D interaction map of 9T-0617 in the main hotspot of WzpB.

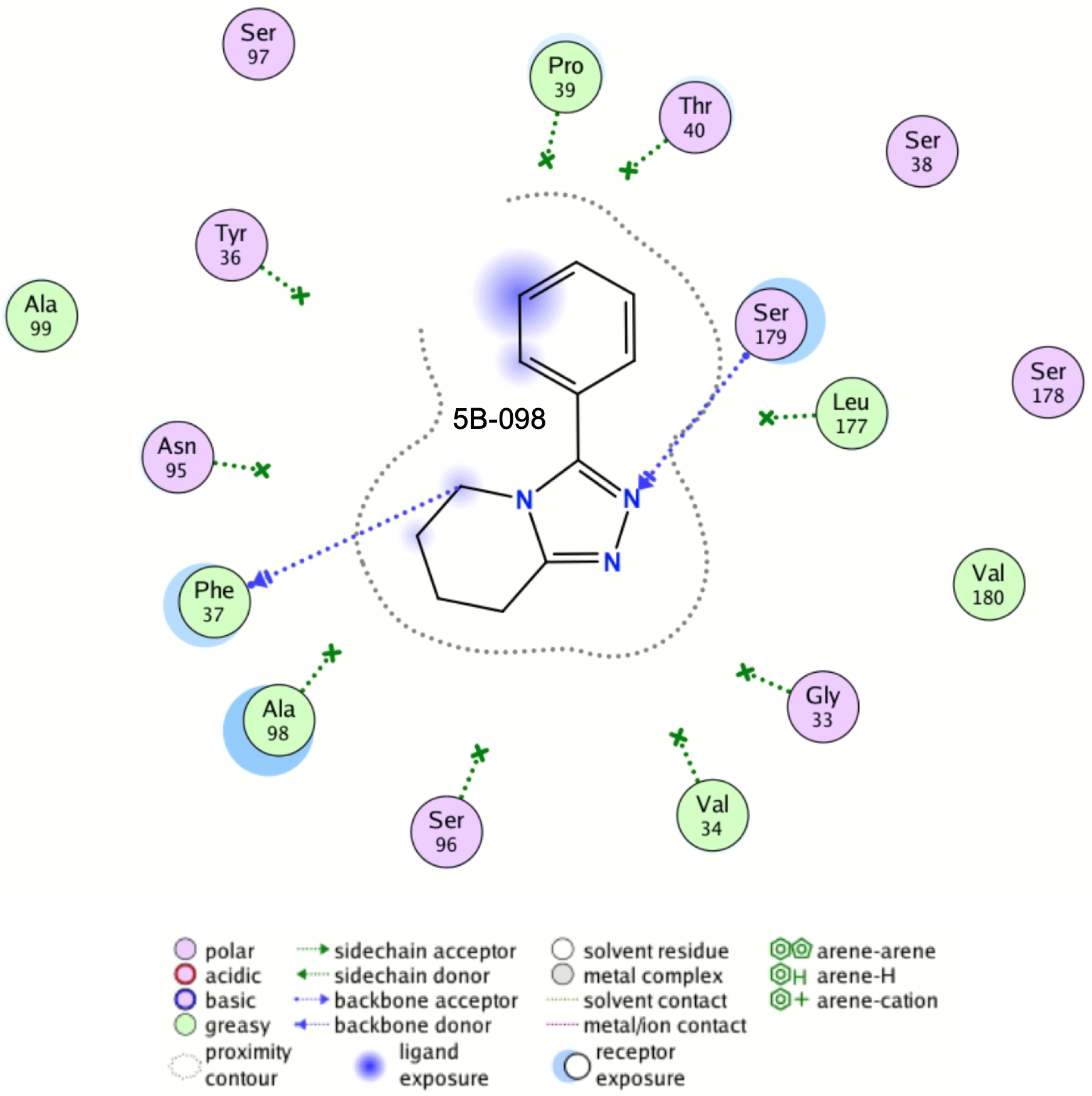

**Fig. S22.** 2D interaction map of 5B-098 in the main hotspot of WzpB.

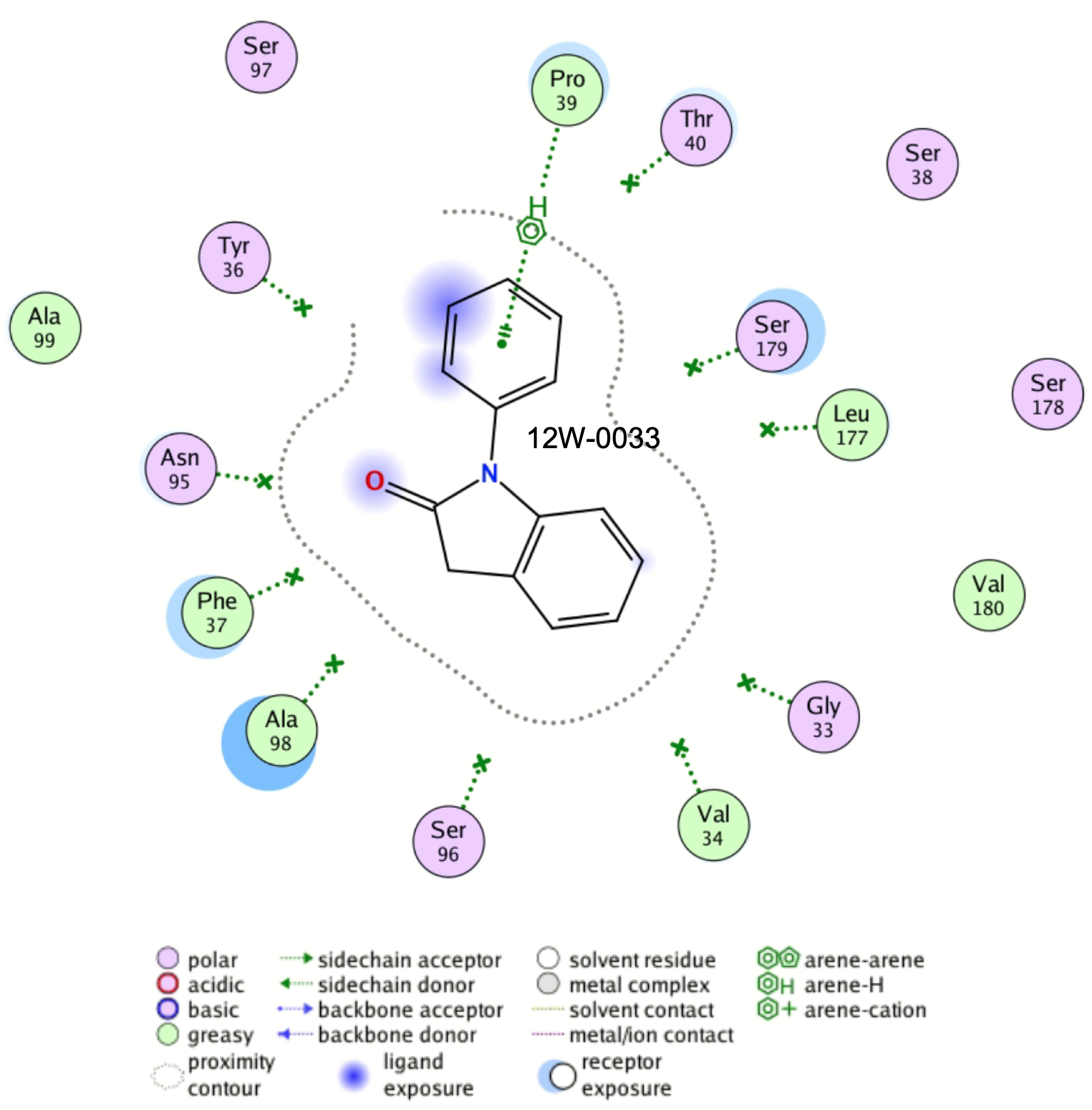

**Fig. S23.** 2D interaction map of 12W-0033 in the main hotspot of WzpB.

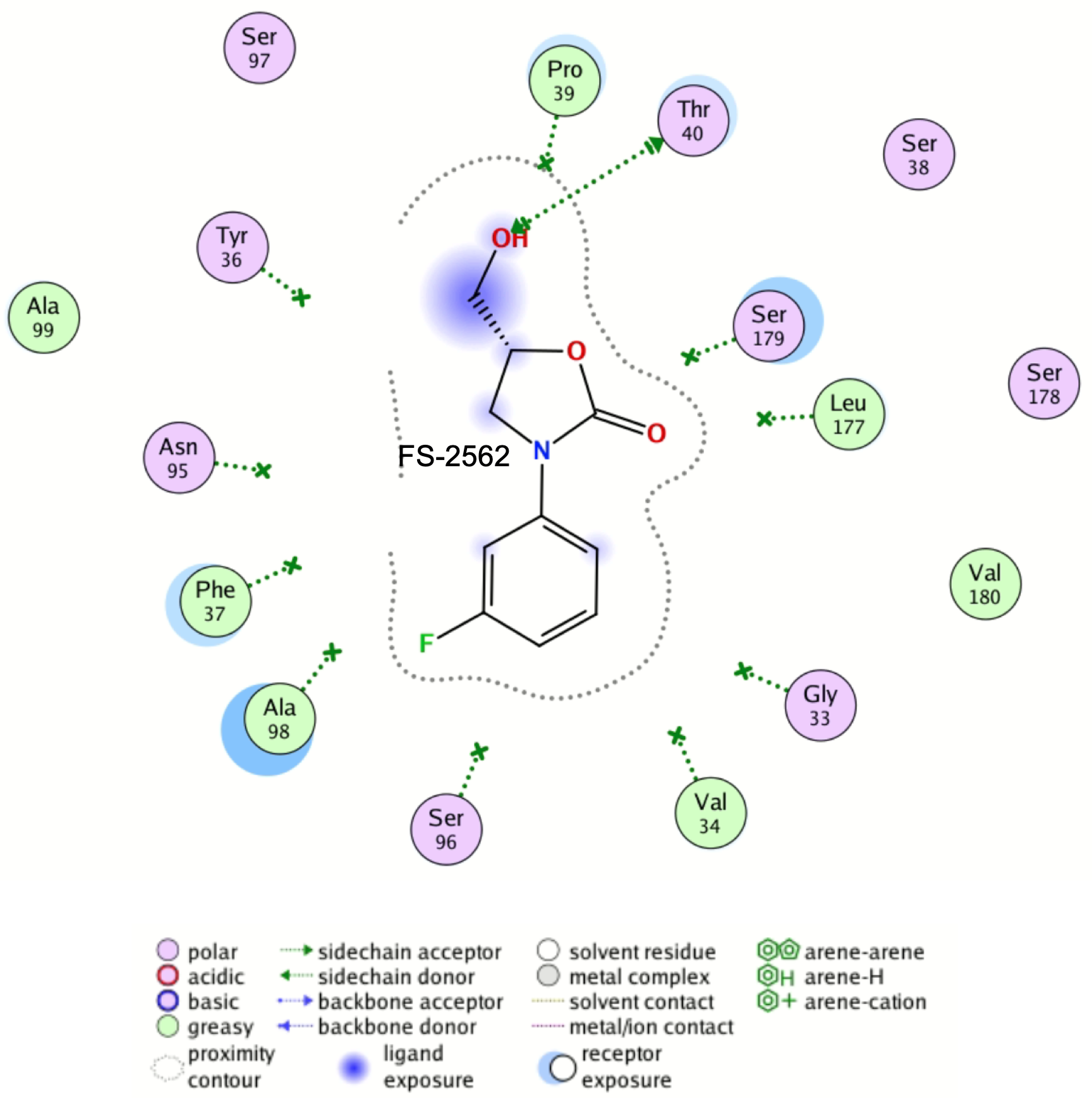

**Fig. S24.** 2D interaction map of FS-2562 in the main hotspot of WzpB.

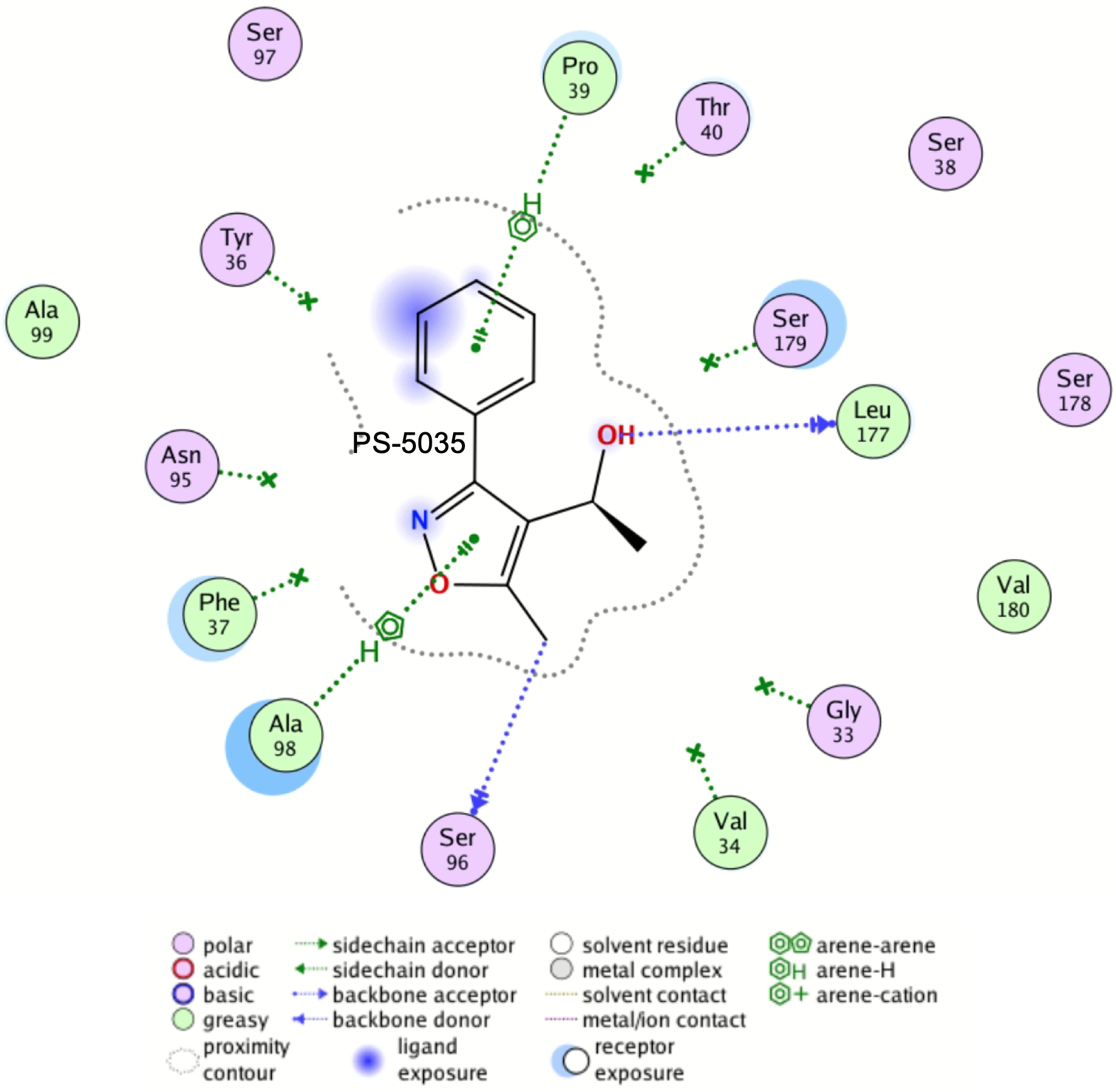

**Fig. S25.** 2D interaction map of PS-5035 in the main hotspot of WzpB.

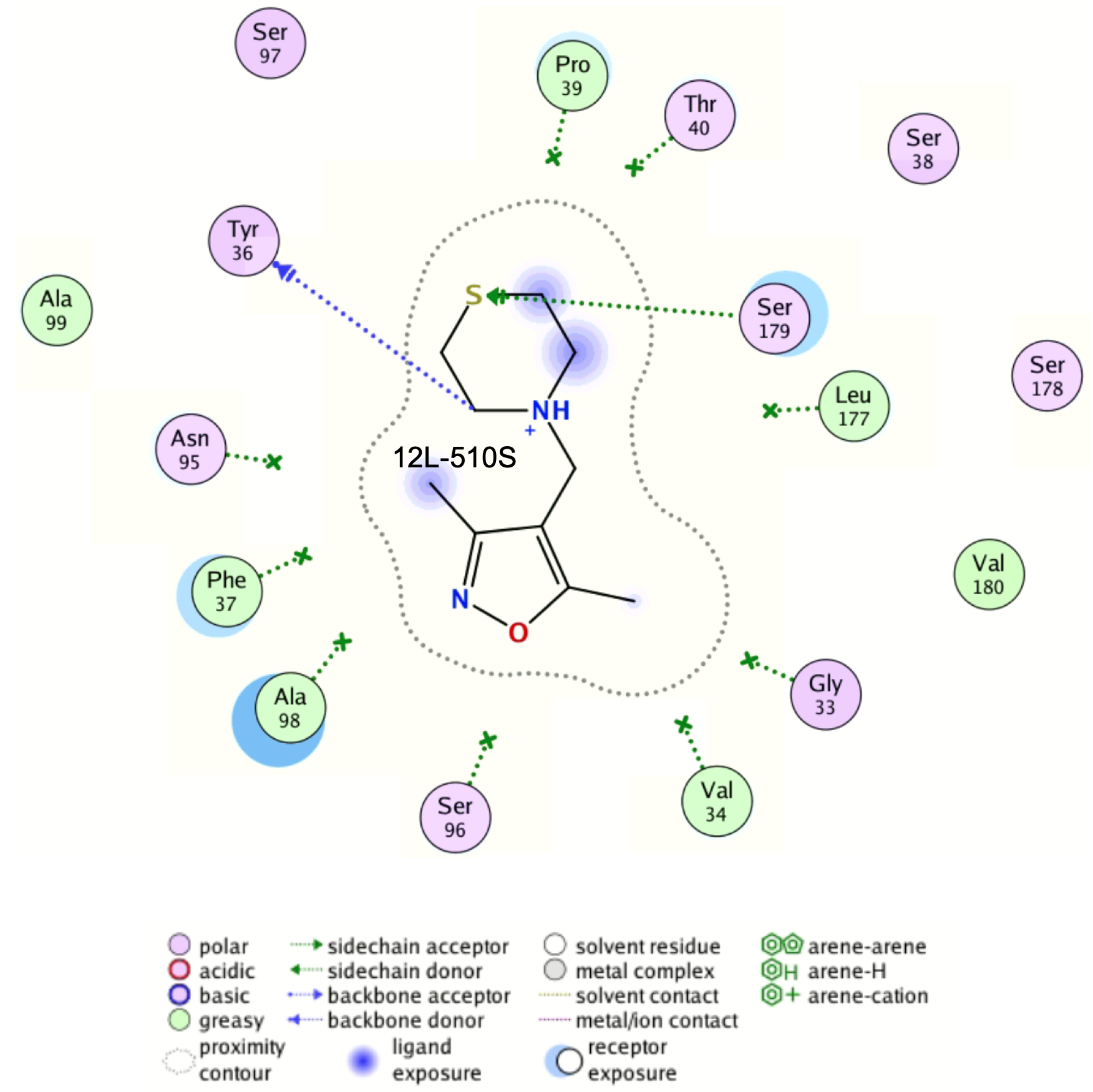

**Fig. S26.** 2D interaction map of 12L-510S in the main hotspot of WzpB.

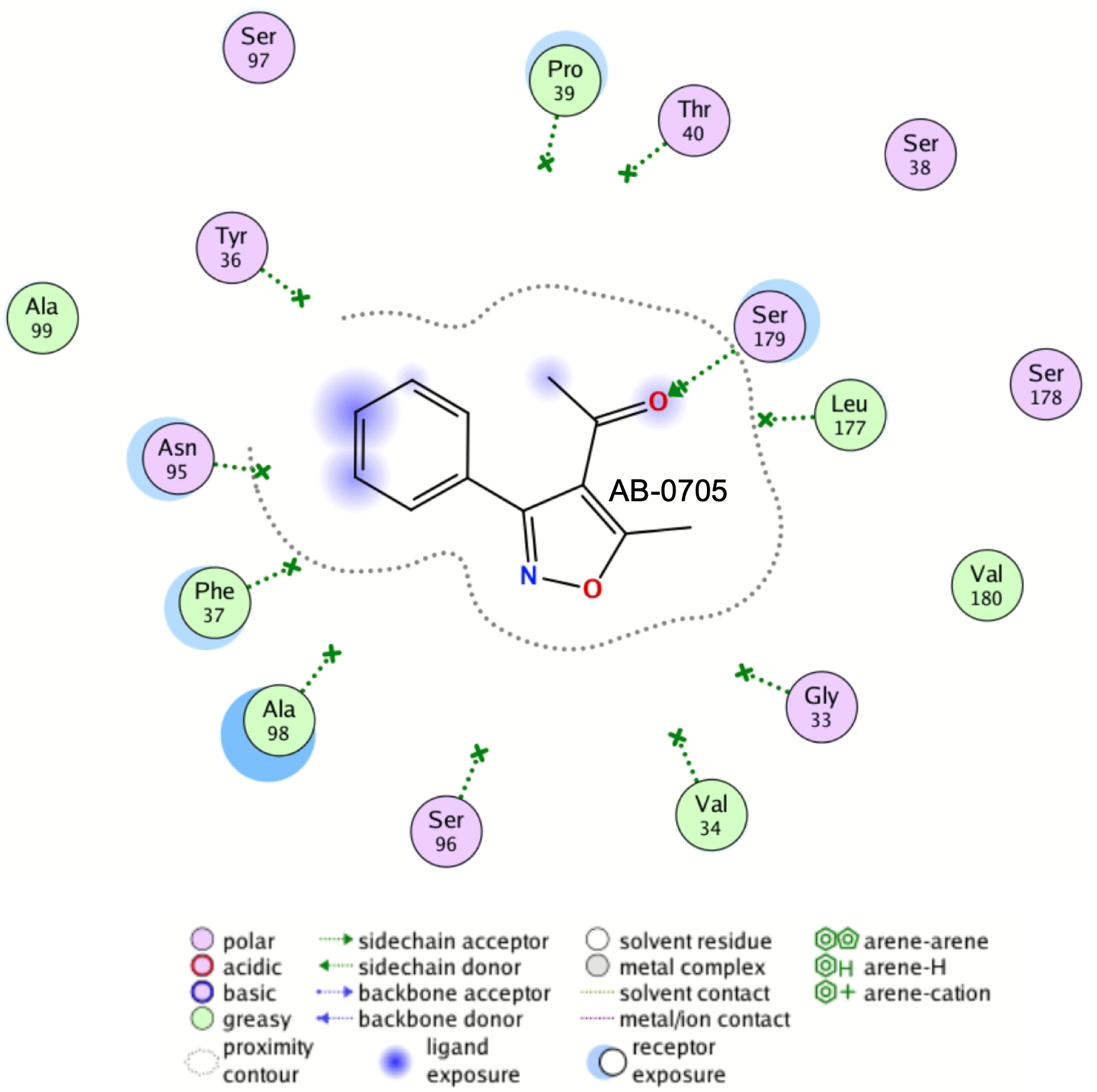

**Fig. S27.** 2D interaction map of AB-0705 in the main hotspot of WzpB.

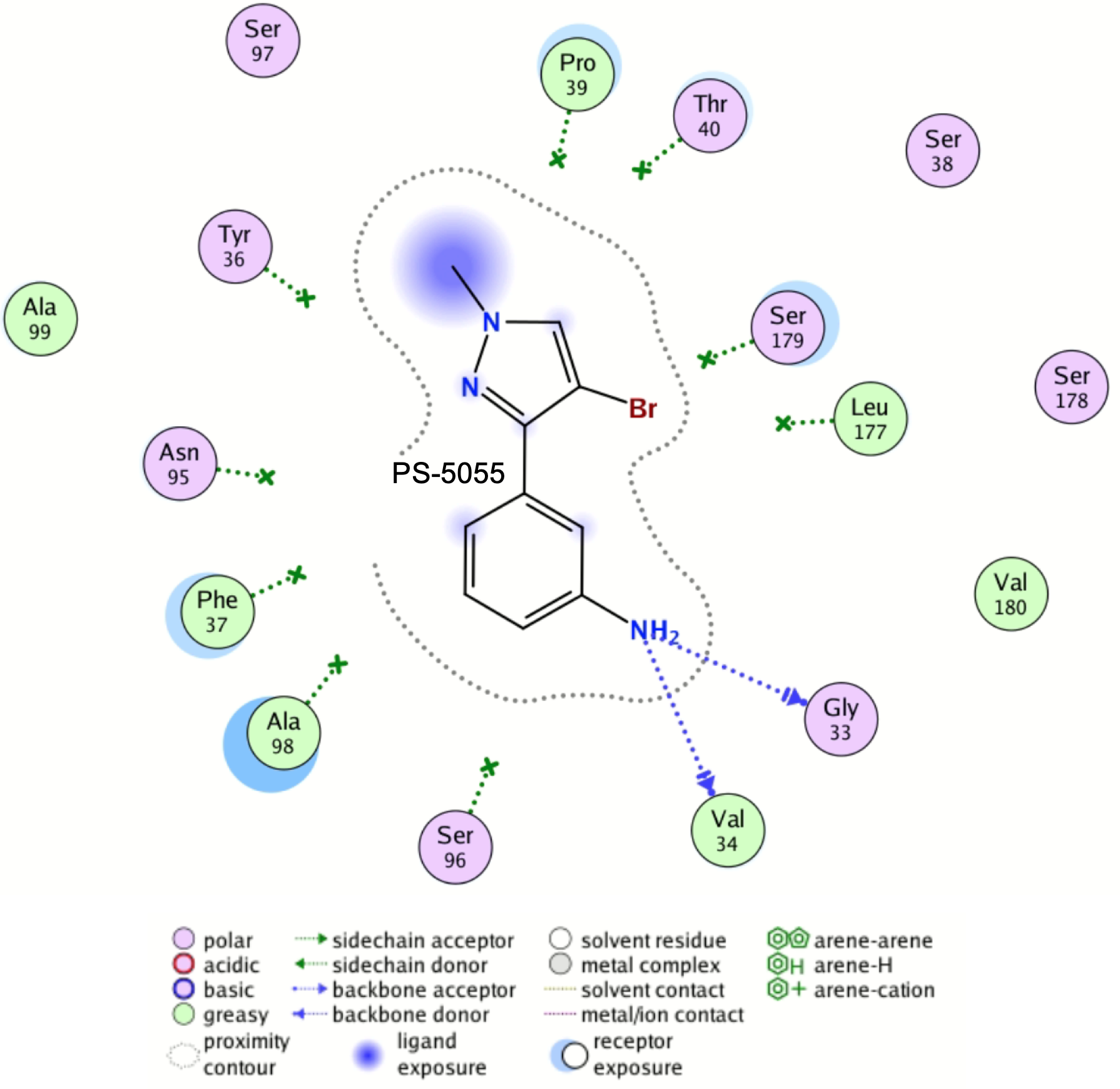

**Fig. S28.** 2D interaction map of PS-5055 in the main hotspot of WzpB.

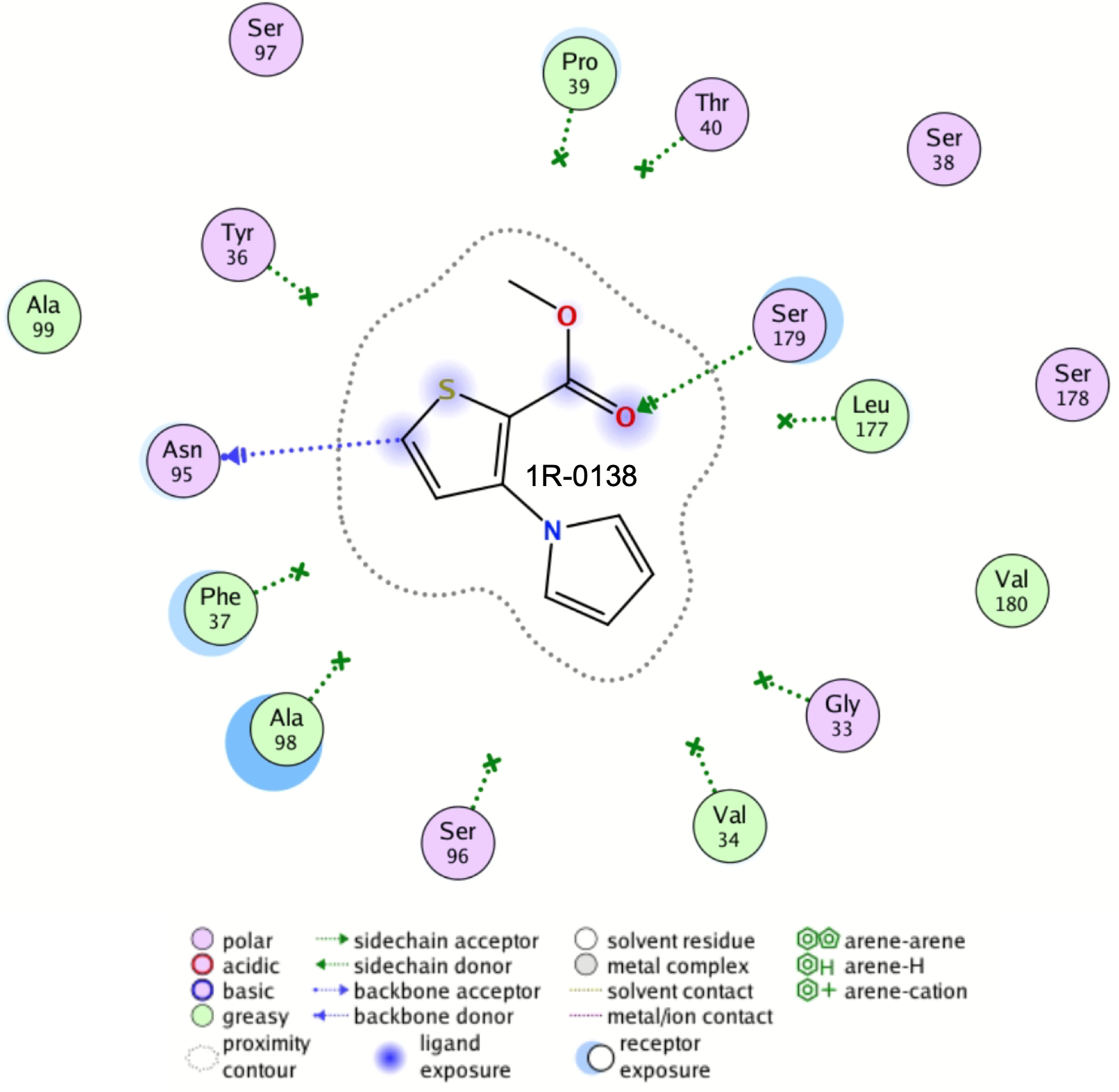

**Fig. S29.** 2D interaction map of 1R-0138 in the main hotspot of WzpB.

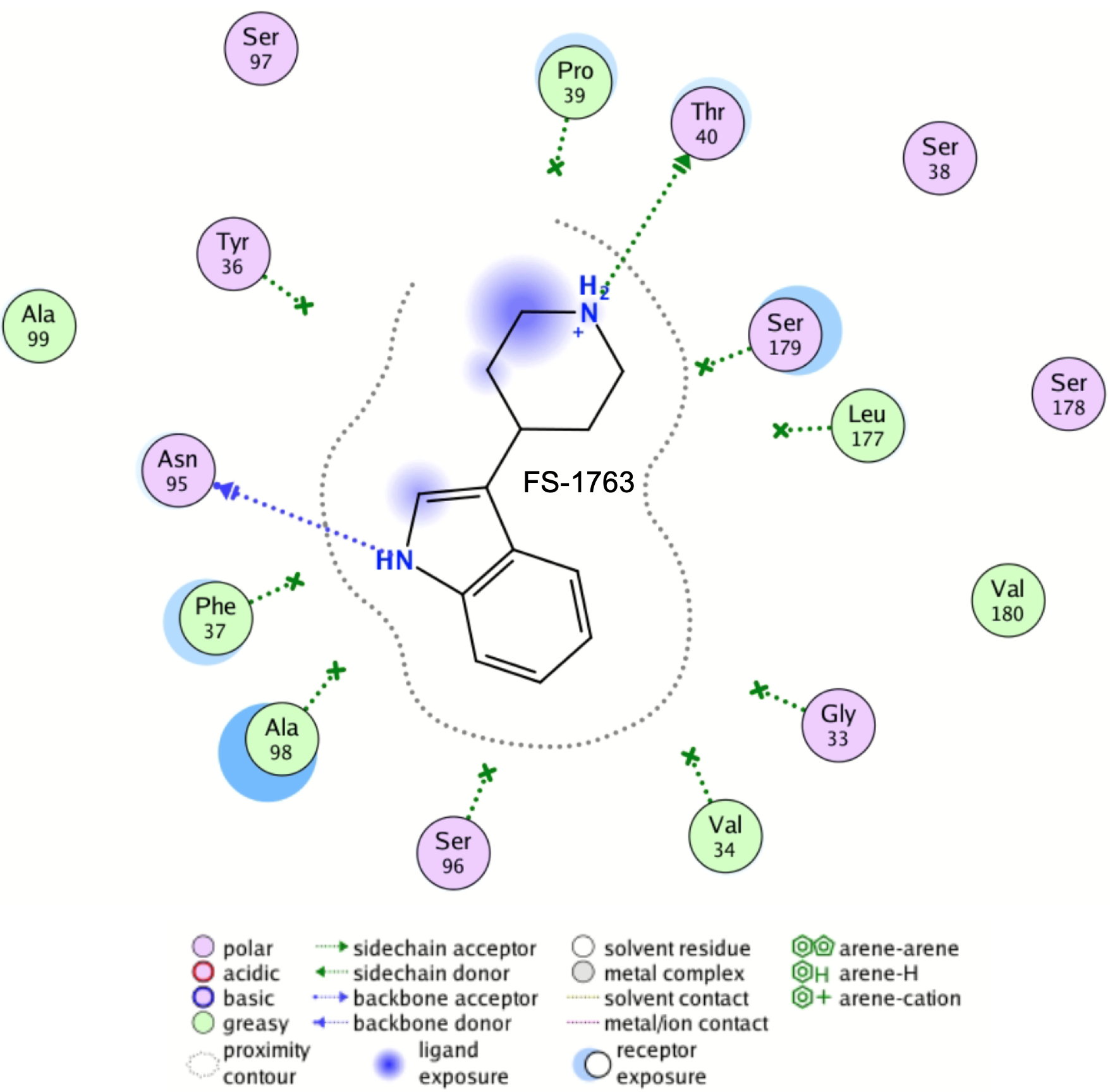

**Fig. S30.** 2D interaction map of FS-1763 in the main hotspot of WzpB.

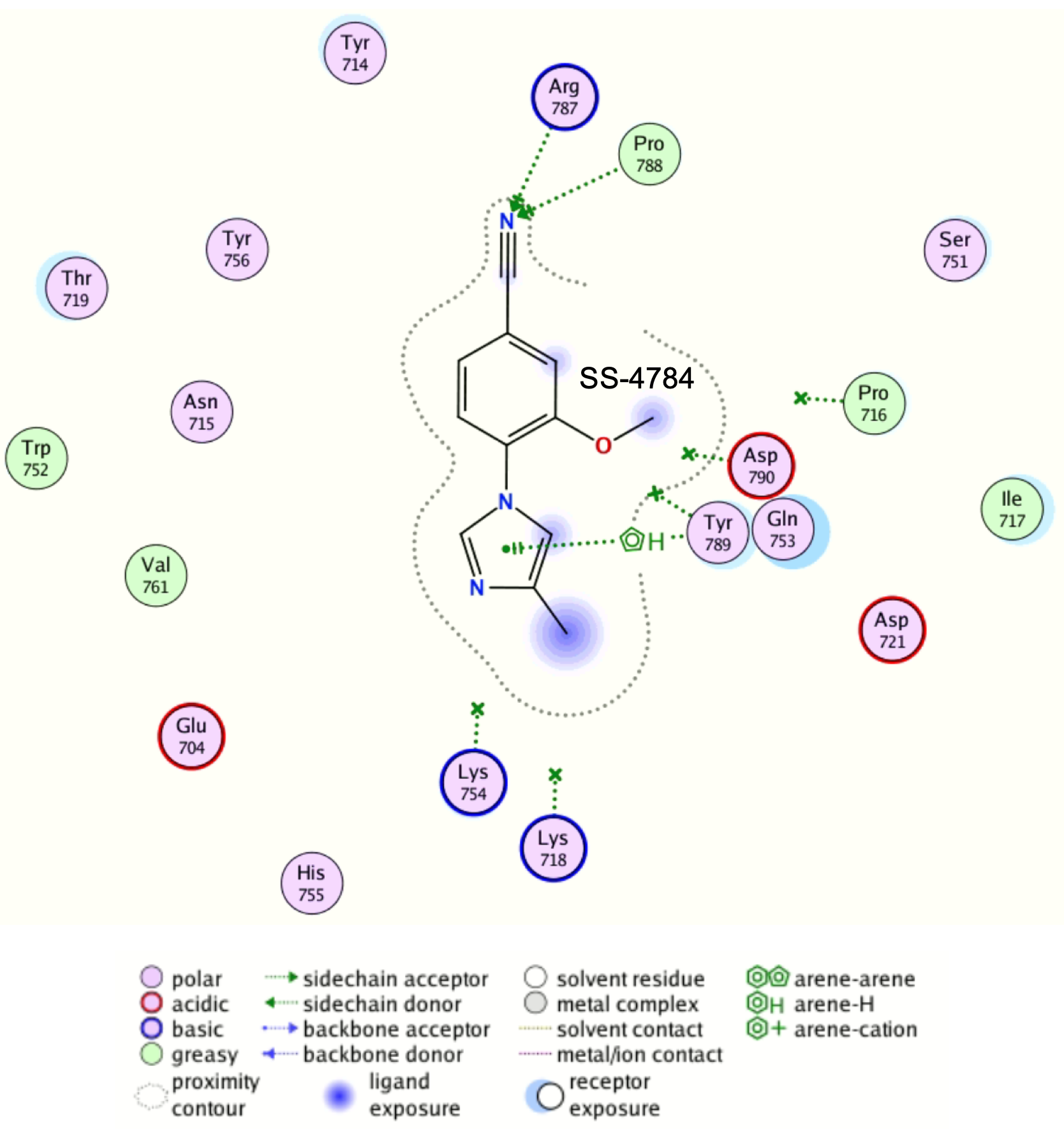

**Fig. S31.** 2D interaction map of SS-4784 in the main hotspot of PgaA.

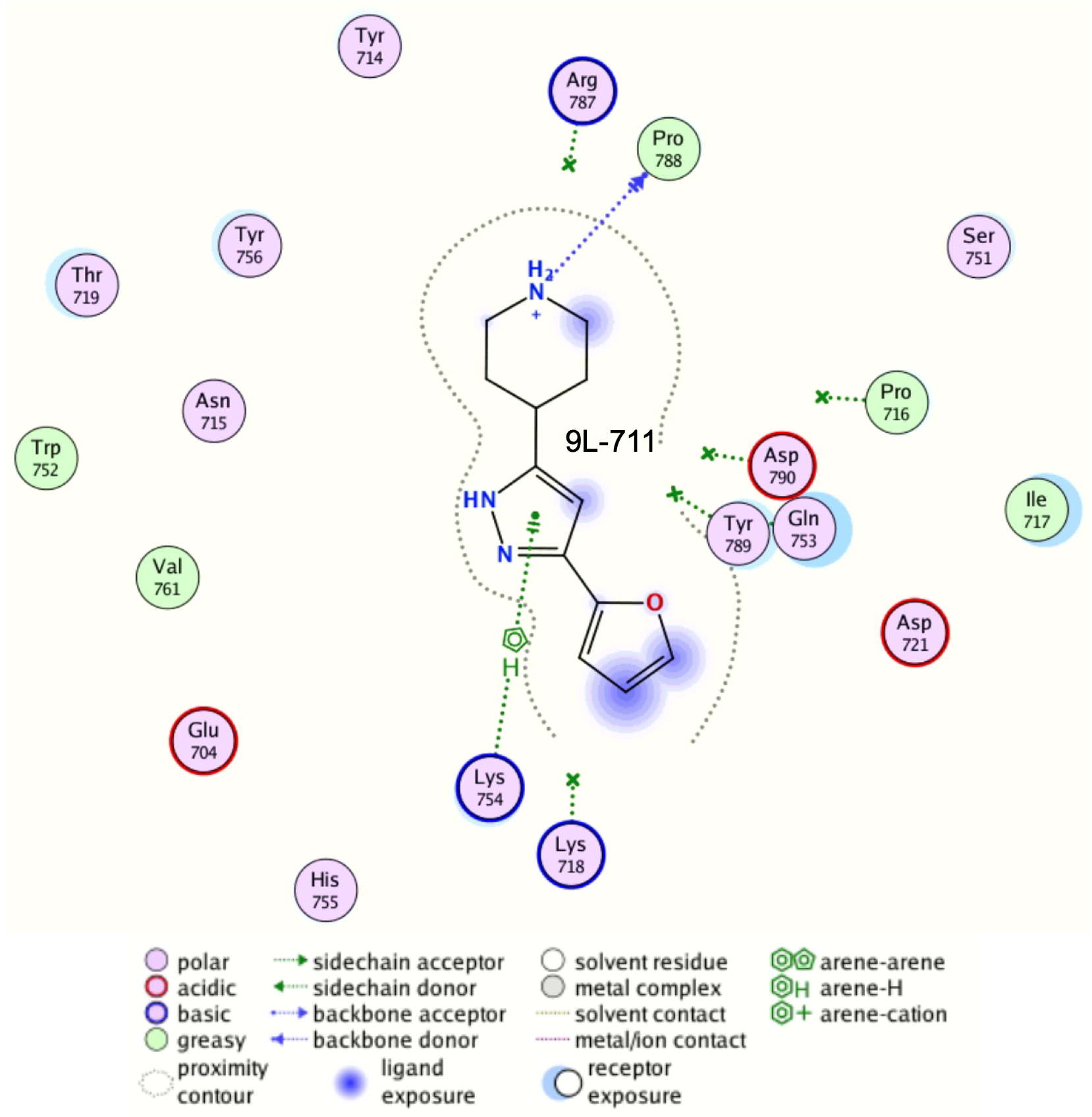

**Fig. S32.** 2D interaction map of 9L-711 in the main hotspot of PgaA.

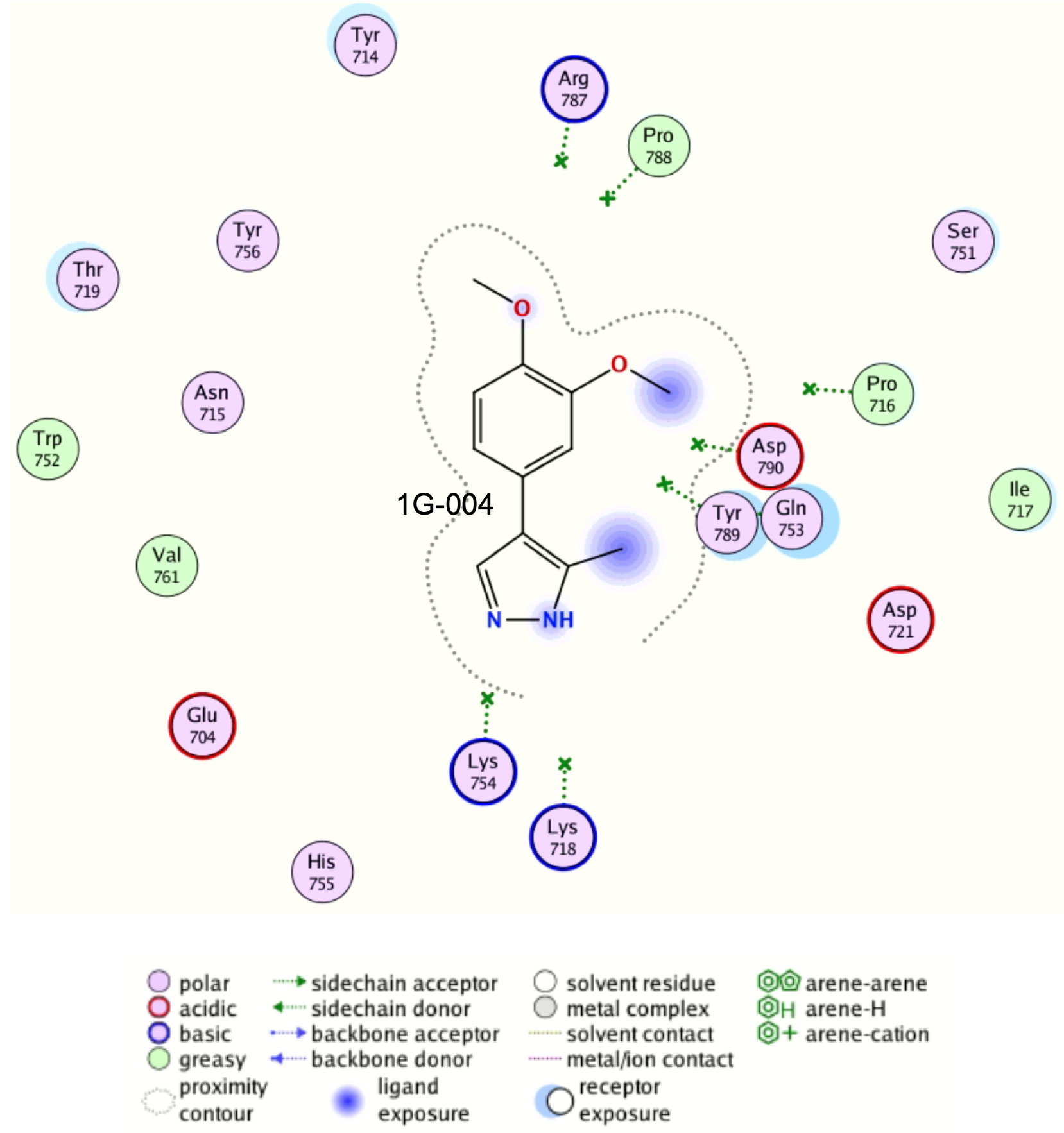

**Fig. S33.** 2D interaction map of 1G-004 in the main hotspot of PgaA.

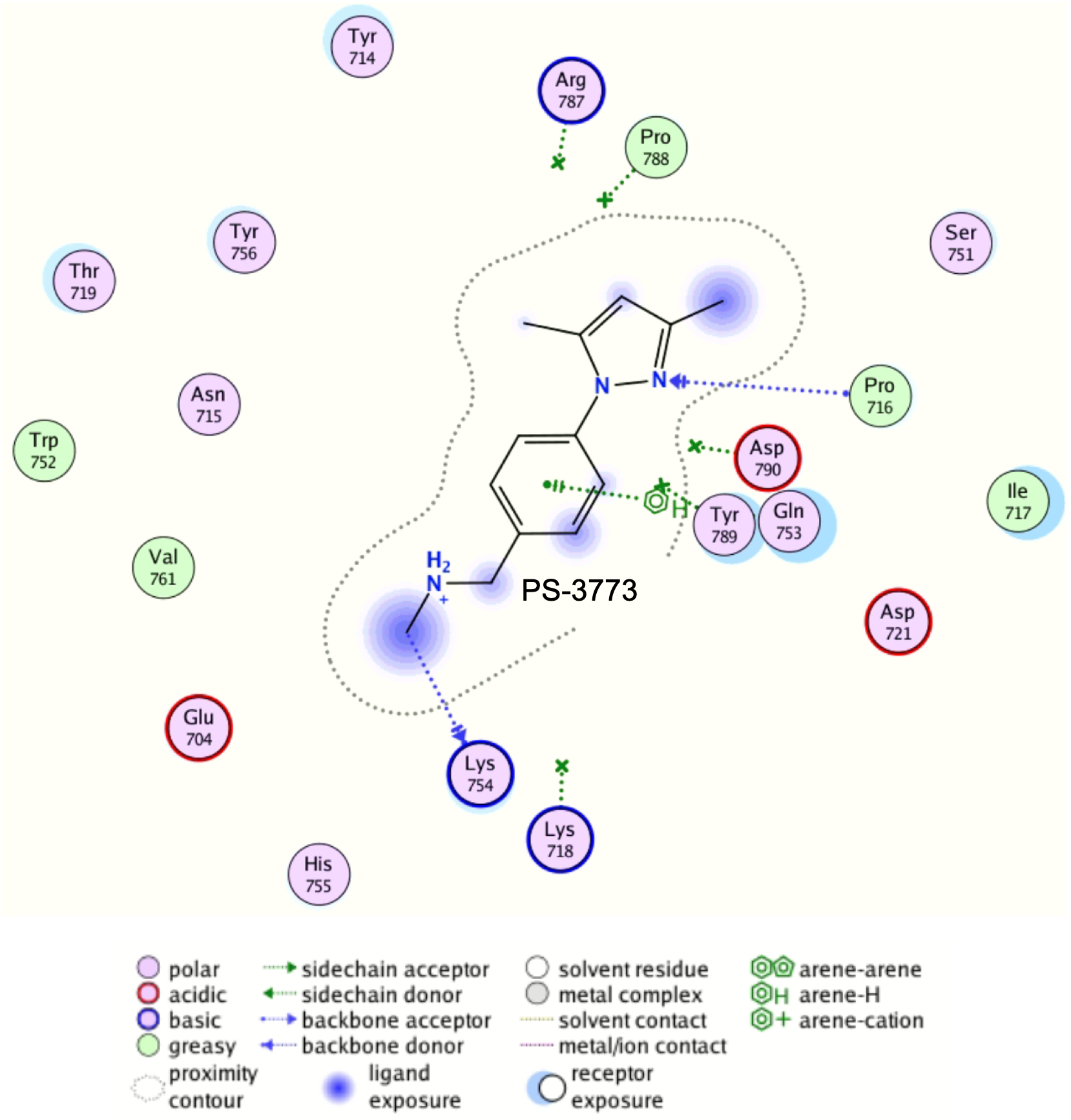

**Fig. S34.** 2D interaction map of PS-3773 in the main hotspot of PgaA.

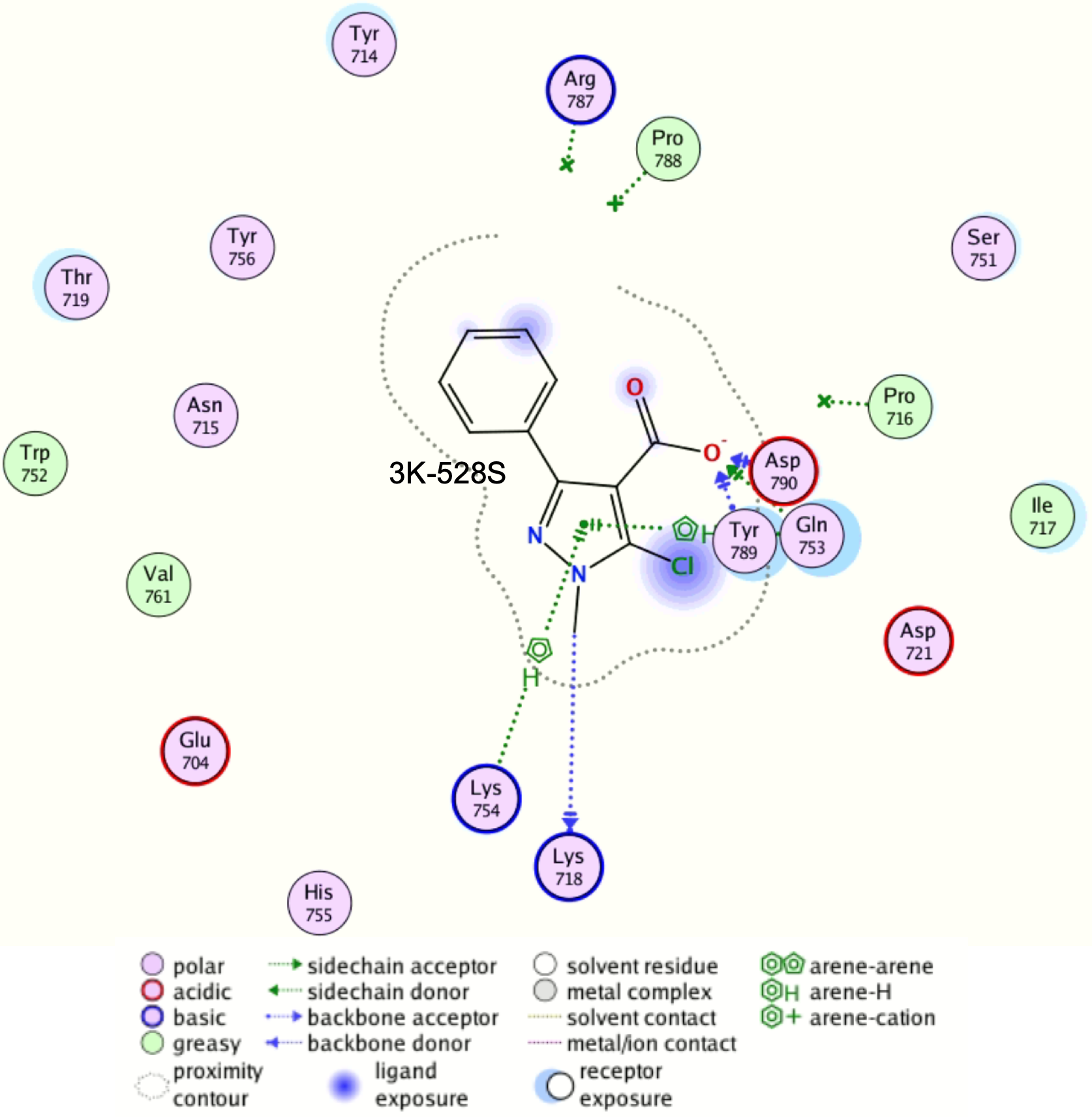

**Fig. S35.** 2D interaction map of 3K-528S in the main hotspot of PgaA.

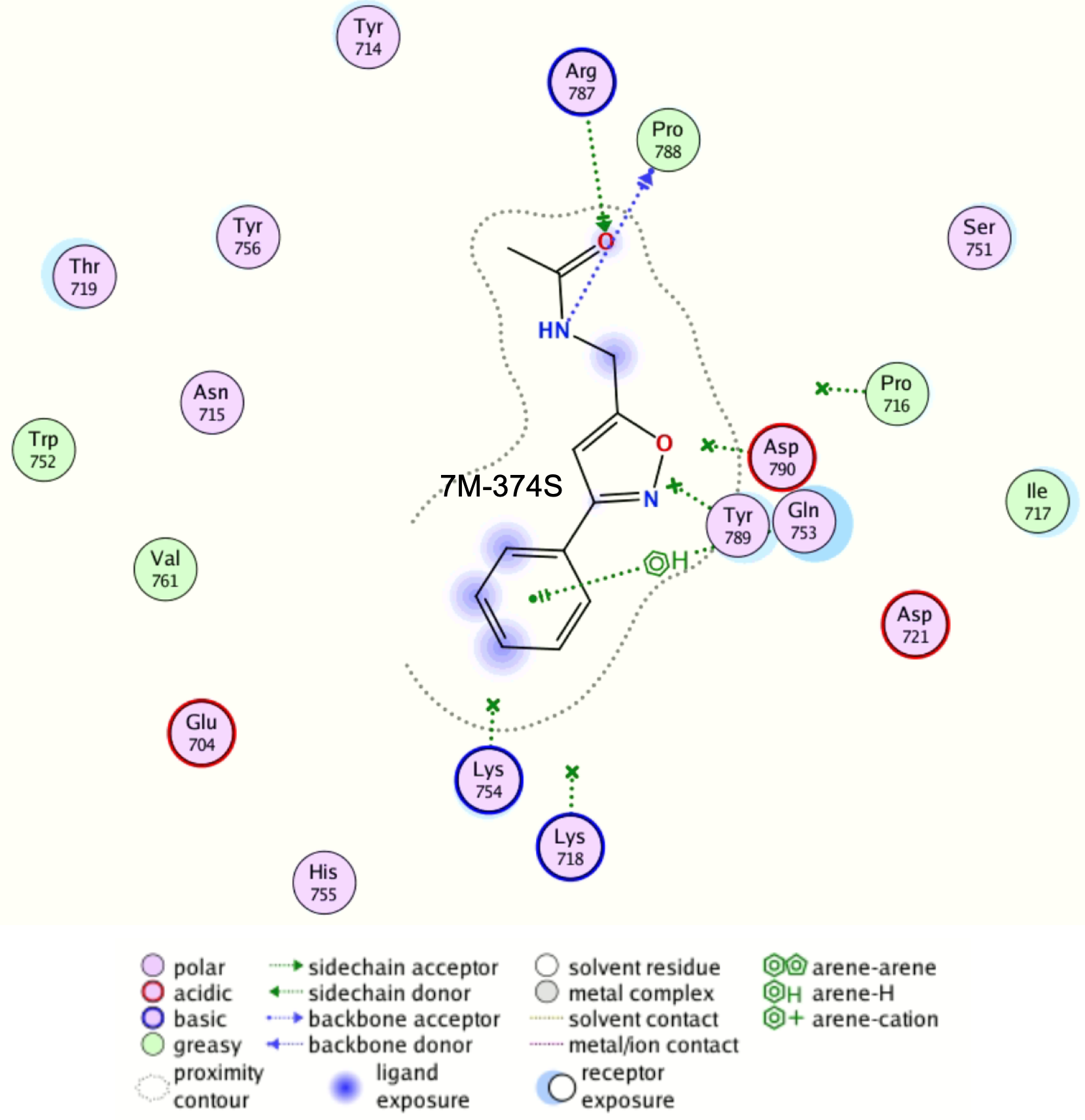

**Fig. S36.** 2D interaction map of 7M-374S in the main hotspot of PgaA.

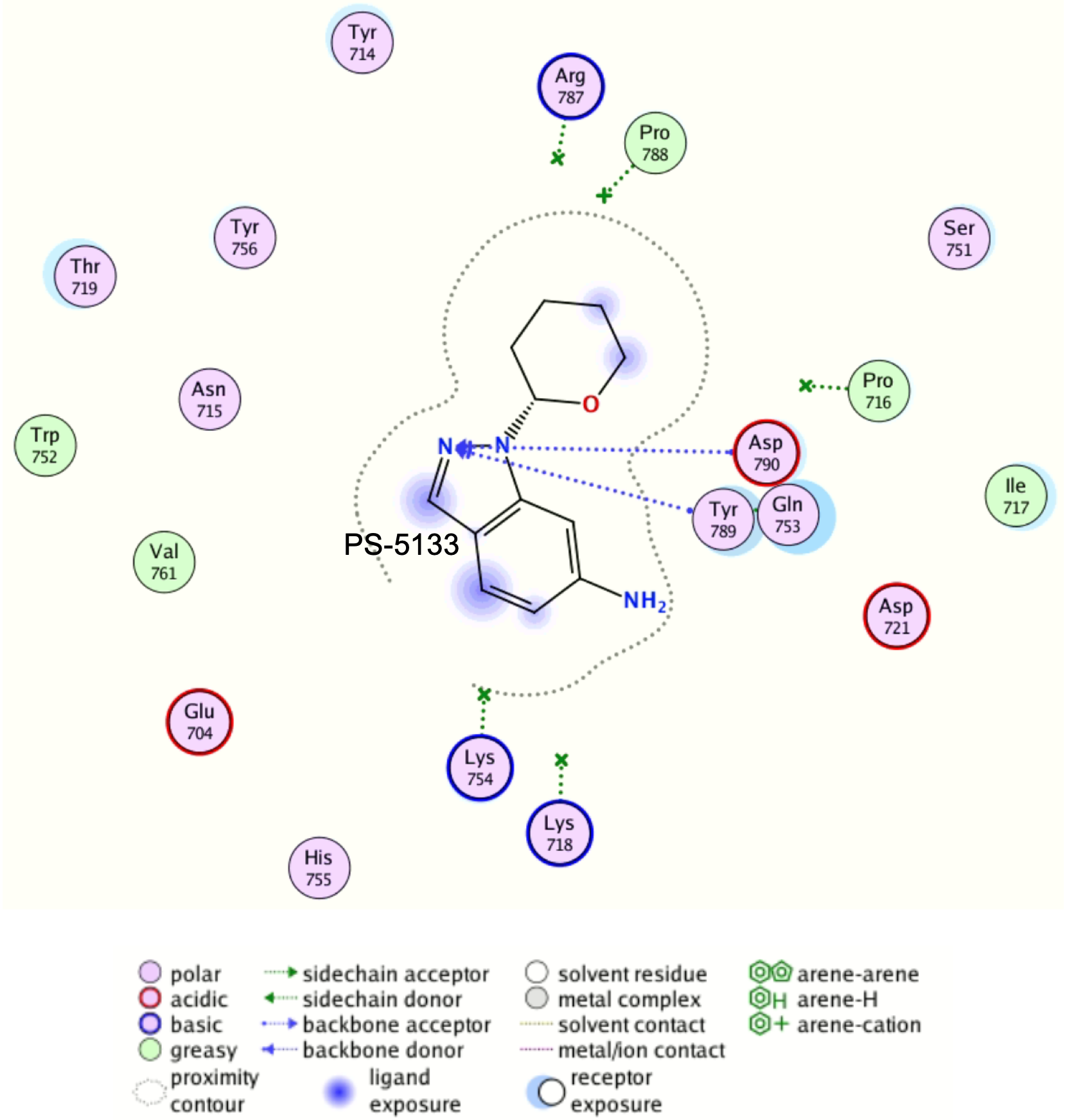

**Fig. S37.** 2D interaction map of PS-5133 in the main hotspot of PgaA.

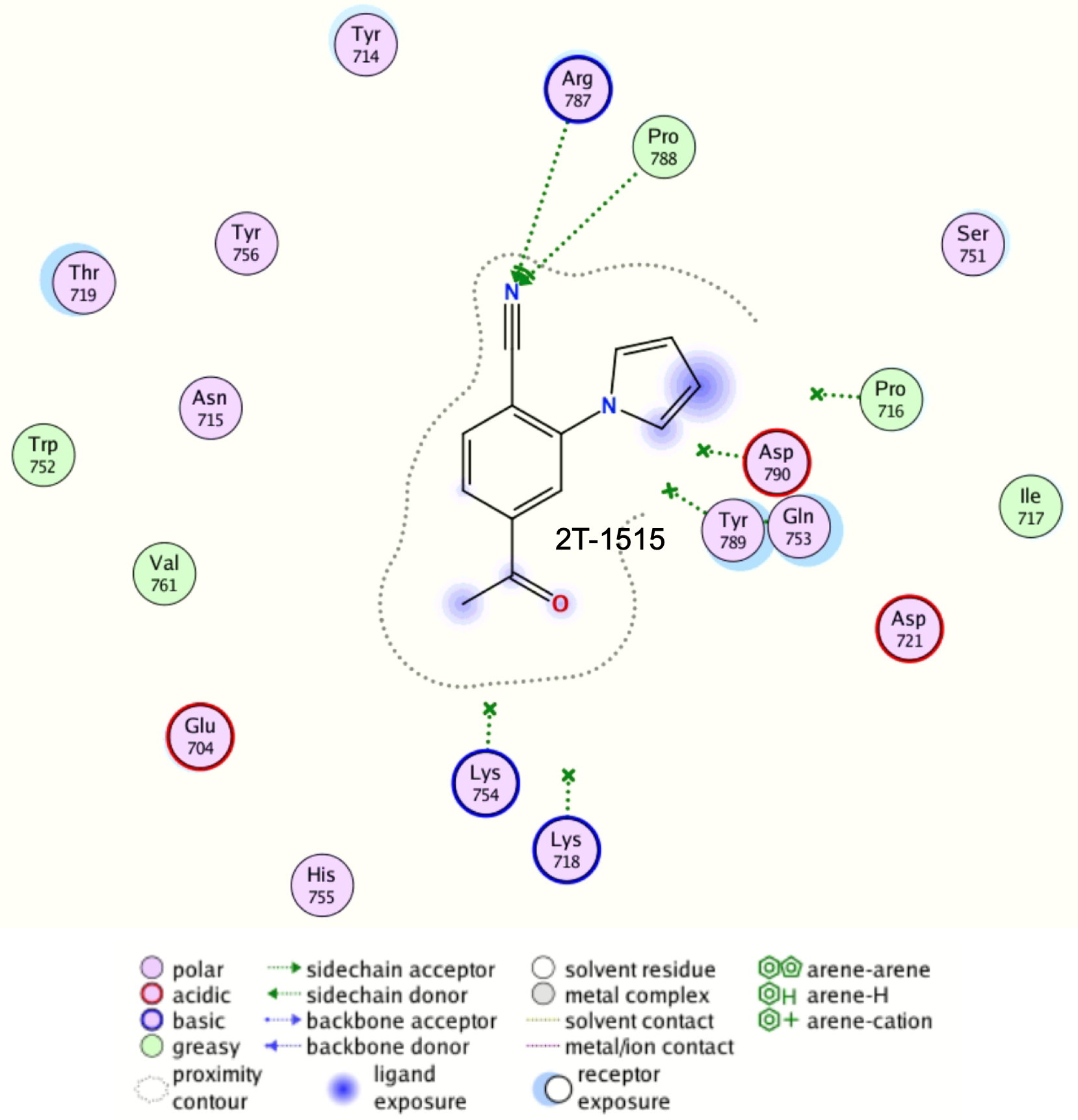

**Fig. S38.** 2D interaction map of 2T-1515 in the main hotspot of PgaA.

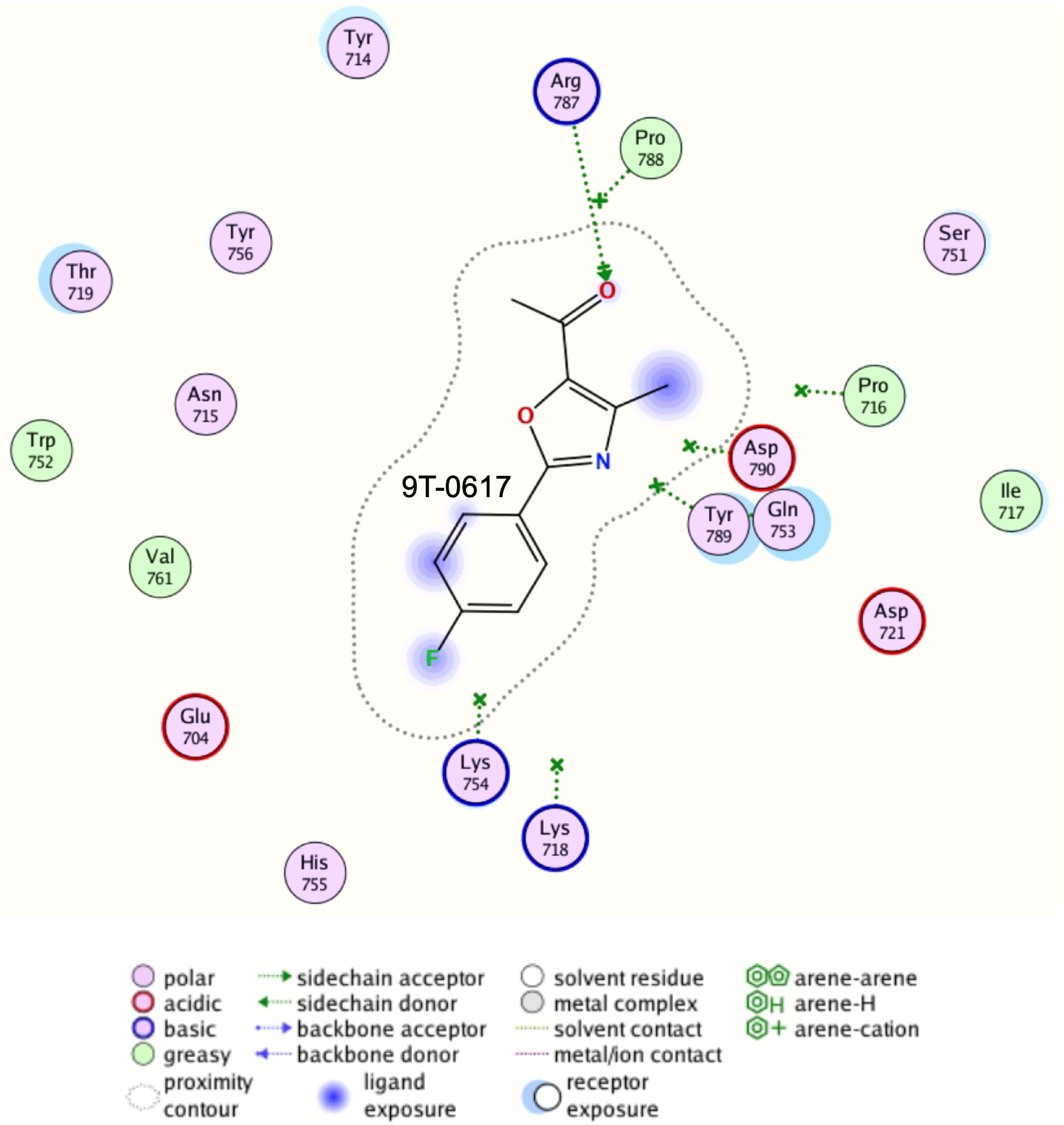

**Fig. S39.** 2D interaction map of 9T-0617 in the main hotspot of PgaA.

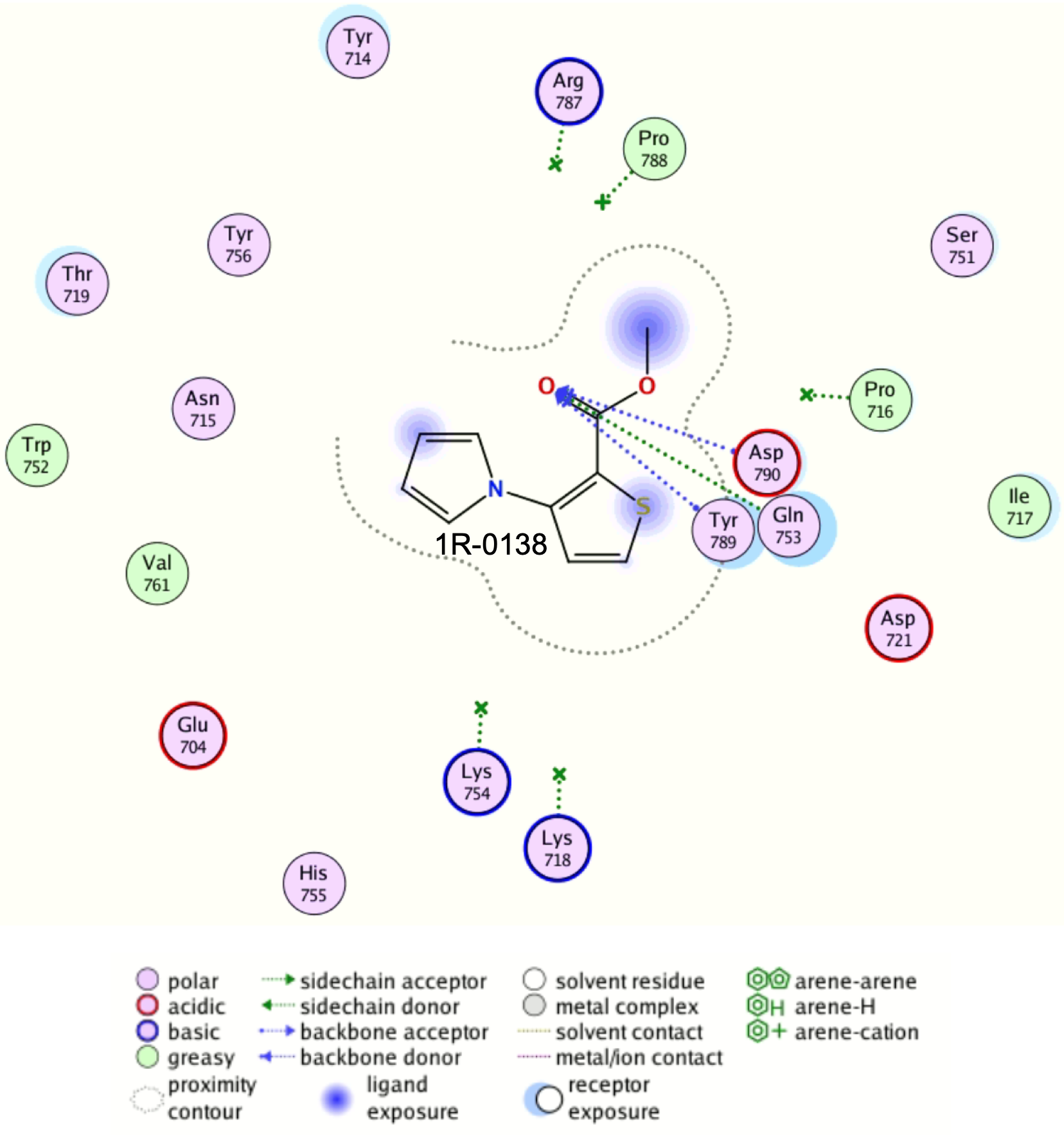

**Fig. S40.** 2D interaction map of 1R-0138 in the main hotspot of PgaA.
